# Directed Differentiation of Human iPSCs to Functional Ovarian Granulosa-Like Cells via Transcription Factor Overexpression

**DOI:** 10.1101/2022.07.04.498717

**Authors:** Merrick Pierson Smela, Christian Kramme, Patrick Fortuna, Jessica Adams, Edward Dong, Mutsumi Kobayashi, Garyk Brixi, Emma Tysinger, Richie. E. Kohman, Toshi Shioda, Pranam Chatterjee, George M. Church

**Affiliations:** Wyss Institute, Harvard Medical School; Department of Genetics, Harvard Medical School; Massachusetts General Hospital Center for Cancer Research, Harvard Medical School; Department of Biomedical Engineering, Duke University; Center for Bits and Atoms, Massachusetts Institute of Technology; Media Lab, Massachusetts Institute of Technology

## Abstract

An *in vitro* model of human ovarian follicles would greatly benefit the study of female reproduction. Ovarian development requires the combination of germ cells and their supporting somatic cells, known as granulosa cells. Whereas efficient protocols exist for generating human primordial germ cell-like cells (hPGCLCs) from human iPSCs, a method of generating granulosa cells has been elusive. Here we report that simultaneous overexpression of two transcription factors (TFs) can direct the differentiation of human iPSCs to granulosa-like cells. We elucidate the regulatory effects of several granulosa-related TFs, and establish that overexpression of *NR5A1* and either *RUNX1* or *RUNX2* is necessary and sufficient to generate granulosa-like cells. Our granulosa-like cells form ovary-like organoids (ovaroids) when aggregated with hPGCLCs, and recapitulate key ovarian phenotypes including support of germ cell maturation, follicle formation, and steroidogenesis. This model system will provide unique opportunities for studying human ovarian biology, and may enable the development of therapies for female reproductive health.

## Introduction

Oogenesis is the central process of female reproduction, yet relative to other animals, little is understood about this process in humans. This is in part due to the lack of a suitable model system to study human ovarian development *in vitro.* The potential power of such a system is illustrated by a recent study in mice, which differentiated mouse embryonic stem cells (mESCs) into primordial germ cell-like cells (PGCLCs) and ovarian granulosa-like cells, and combined these cell types. Reconstituted follicles were capable of producing oocytes and live offspring^1^. Using mouse fetal ovarian somatic cells, a similar system was published that allowed the maturation of human induced pluripotent stem cell (hiPSC)-derived PGCLCs to the oogonia stage^2^. However, as differences between mouse and human ovarian development are still poorly understood, xenobiotic systems involving mouse ovarian somatic cells and hPGCLCs have apparent limits in supporting human oogenesis. Clearly, the construction of fully human ovarian organoids from pluripotent stem cells could enable important advances in the study of female reproduction.

Within the ovary, granulosa cells surround and support developing oocytes^3–6^, and additionally play key roles in hormonal signaling throughout adult life^7^. Therefore, these cells are a crucial component of any *in vitro* model of the human ovary. Although a few studies have attempted differentiating hiPSCs to granulosa-like cells by treatment with growth factors^8–10^, none has resulted in an efficient method, nor has any study evaluated their ability to support germ cell development. Therefore, we took a different approach based on the principle that cell identity can be manipulated through transcription factor (TF) overexpression^11^. Encouragingly, a recent study demonstrated that overexpression of the TFs NR5A1 and GATA4 is sufficient to transdifferentiate human fibroblasts into Sertoli-like cells^12^, the male equivalent of granulosa-like cells. Here, we report a set of TFs that enables an efficient, scalable method of generating granulosa-like cells from hiPSCs, and show that these granulosa-like cells engage in hormonal signaling, support germ cell development, and form follicle-like structures.

## Results

### *In silico* prediction of granulosa cell-regulating transcription factors

We began by predicting candidate TFs that could direct the differentiation of hiPSCs to granulosa-like cells. We first selected 22 TFs that were differentially expressed in granulosa cells compared to hESCs and early mesoderm, using previously published datasets for these cell types^13,14^. We also included 5 TFs based on previous developmental biology studies, mainly showing that these TFs were important for mouse ovarian development^15–19^. Finally, we identified 9 additional TFs that we predicted to be upstream of the others on the list, based on a gene regulatory network analysis taking into account co-expression data as well as binding motifs^20^. The list of TFs (35 in total) is given in Table S1. Next, we assembled a PiggyBac library by cloning cDNAs into a barcoded destination plasmid. The TFs were expressed under the control of a doxycycline-inducible promoter.

### Overexpression screening identifies TFs that drive granulosa-like cell formation

To enable identification of granulosa-like cells, we used CRISPR/Cas9 and HDR to engineer an hiPSC line with a homozygous knock-in of a T2A-tdTomato reporter at the C-terminus of FOXL2 (Figure S1). We chose FOXL2 due to its specific expression in granulosa cells^21,22^. Next, the pooled cDNA library was co-electroporated with a PiggyBac transposase expression plasmid, and a population of cells with integrated transposons was selected by treatment with puromycin. In these experiments, we tested several different library compositions and DNA concentrations (see Methods, Table S1, and Figure S2).

For screening, we induced TF expression with doxycycline, and sorted reporter-positive cells after 5 days of treatment. In addition to screening TF expression in the pluripotency-supporting mTeSR Plus medium, we also expressed TFs following differentiation of hiPSCs to early-stage mesoderm by treatment with CHIR99021. The TF expression resulted in a small fraction of FOXL2+ cells, from which we extracted gDNA and sequenced barcodes. In both conditions, we observed barcodes for *NR5A1* to be strongly enriched in FOXL2+ cells relative to negative cells, as well as relative to the barcodes in the pre-induced hiPSC population (Figure 1A). Other TFs showed more modest barcode enrichment *(RUNX2, GATA4, TCF21),* or enrichment in only one condition *(KLF2, NR2F2).* Interestingly, barcodes for *FOXL2* were strongly depleted, suggesting negative feedback whereby exogenous FOXL2 directly or indirectly suppresses the expression of the endogenous FOXL2 reporter allele.

**Figure 1.**
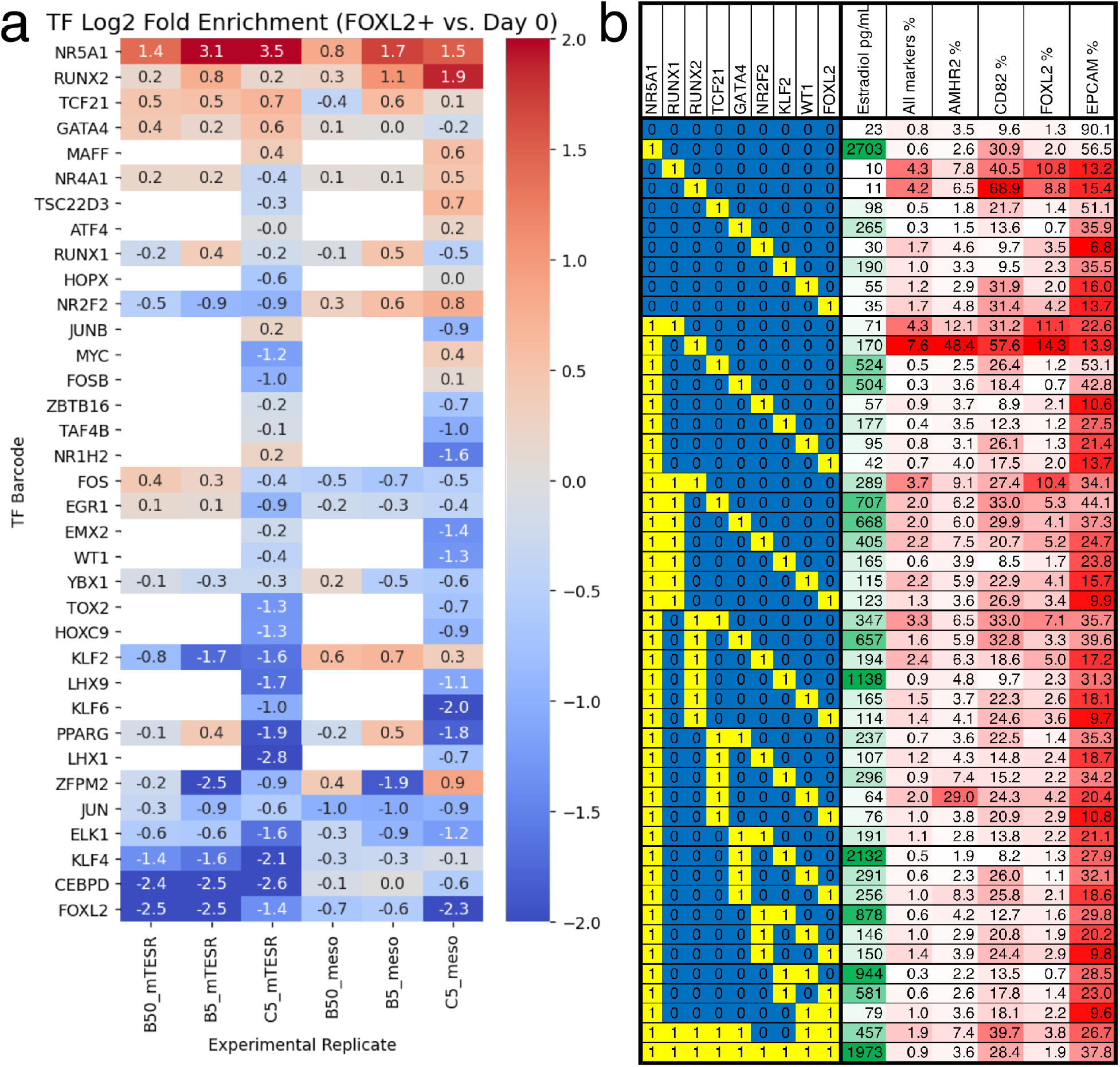
Identification of TFs whose overexpression generates granulosa-like cells. **(A)** Pooled screening of barcoded TF cDNA libraries (see Methods for details) identifies TFs enriched in FOXL2-T2A-tdTomato+ cells. **(B)** Combinatorial screening identifies minimal TF combinations for inducing granulosa-like cells. TF combinations were integrated into hiPSCs; a “1” in the left-hand box signifies presence of the TF in the combination corresponding to that row. For each combination, the polyclonal hiPSC population was differentiated with TF induction (see Methods). For the last 24hr of differentiation, cells were additionally treated with FSH and androstenedione. Estradiol production and granulosa markers were measured by ELISA and flow cytometry after a total of 5 days. NR5A1 expression induced high levels of estradiol synthesis, but the combination of NR5A1 with RUNX1 or RUNX2 was required to give the best results for granulosa markers.

Using the top TFs *(NR5A1, RUNX1/RUNX2, TCF21, GATA4)* we then further optimized the conditions for generation of FOXL2+ cells. We included *RUNX1* in this list because the two TFs are structurally and functionally similar, and *RUNX1* is known to play an important role for granulosa cell maintenance in the mouse^19^. We integrated these TFs into hiPSCs and established monoclonal lines, which we screened by flow cytometry after TF induction. We monitored FOXL2-tdTomato as well as the surface markers CD82, follicle stimulating hormone receptor (FSHR), and EpCAM. CD82 is absent in hiPSCs, but highly expressed in granulosa cells beginning at the primordial follicle stage^14^. FSHR is specific for late-stage (secondary/antral) granulosa cells and Sertoli cells (the male equivalent). By contrast, EpCAM is expressed in hiPSCs and epithelial cells, but not granulosa cells^14^.

Out of 23 lines tested, 4 showed >50% FOXL2-tdTomato+ CD82+ EpCAM–cells after 5 days of differentiation (Figure S3A). Further optimization by adding the GSK3 inhibitor CHIR99021 for the first 2 days resulted in a near-homogeneous FOXL2-tdTomato+ CD82+ EPCAM–population for the top cell lines (Figure S4A), with this effect being doxycycline-responsive (Figure S4B). We observed low levels of FSHR expression, suggesting that we were mainly generating cells corresponding to early (primordial/primary follicle) granulosa cells, given that FSHR only becomes strongly expressed at later stages of ovarian follicle development^23^. Genotyping of two top lines revealed that both had *NR5A1* and *TCF21* expression cassettes integrated, with the first line also having *RUNX1* and the second line *RUNX2*.

Subsequently, we set out to determine which TFs were necessary and sufficient for induction of granulosa-like cells. We tested combinations of hit TFs from our screening *(NR5A1, RUNX2, TCF21, GATA4, KLF2, NR2F2)* as well as TFs reported to be important for ovarian function *(RUNX1, WT1* [–KTS isoform], *FOXL2*). We tested expression of each TF individually, as well as combinations of other TFs and our top hit *NR5A1.* In addition to flow cytometry, we also measured production of estradiol after treatment of the cells with FSH and androstenedione. We observed that combinations containing *NR5A1* and *RUNX1* or *RUNX2* upregulated FOXL2-tdTomato expression, as well as granulosa surface markers AMHR2 and CD82 (Figure 1B). Estradiol production was strongly induced by *NR5A1,* and weakly by *GATA4* and *KLF2. RUNX1* and *RUNX2* expression somewhat decreased estradiol production, suggesting a regulatory role of these factors. Overall, these results indicate that *NR5A1* and either *RUNX1* or *RUNX2* are necessary and sufficient to induce a granulosa-like phenotype.

### TF-mediated differentiation drives granulosa-like cell formation based on gene expression signatures

We next examined the gene expression of our granulosa-like cells. We compared the bulk transcriptomes of hiPSCs, COV434 ovarian tumor cells, and granulosa-like cells (sorted FOXL2+ CD82+ cells from day 5 of a polyclonal differentiation with expression of NR5A1, TCF21, GATA4, and RUNX1). As an additional control, we included hiPSCs differentiated under the same conditions but without TF induction.

In the absence of TF expression, the hiPSCs differentiated into cells expressing mesoderm markers (full gene expression data are provided in Supplementary File 2). At day 5 of differentiation, we observed strong upregulation relative to hiPSCs of genes associated with both lateral mesoderm *(HAND1, BMP5)* as well as paraxial mesoderm *(PAX3),* which potentially indicates the presence of a heterogeneous population. Upregulated genes were enriched for GO terms related to heart, blood vessel, muscle, and skeletal development (Supplementary File 3).

With TF induction, we observed expression of gonadal and granulosa markers, notably including *AMHR2, CD82, FOXL2, FSHR, IGFBP7, KRT19, STAR,* and *WNT4* (Figure 2A). The ovarian stromal/theca cell marker *NR2F2* was also upregulated. The expression levels (TPM) were generally comparable to those observed in previously published data from granulosa cells and human fetal gonad^14,24^. The only major exception was *WT1,* which had much weaker expression than *in vivo* (although still greater than hiPSCs). We also examined the expression of pluripotent and adrenal markers to check for incomplete or off-target differentiation. We did not observe adrenal marker expression, although we did note some residual *POU5F1* expression that may indicate that our polyclonal population did not fully differentiate. Finally, COV434 cells, which were commonly considered as granulosa-like cells but recently reclassified as small cell ovarian carcinoma^25–27^, did not express most granulosa markers, but did express high levels of *WT1* and *NR2F2*.

**Figure 2.**
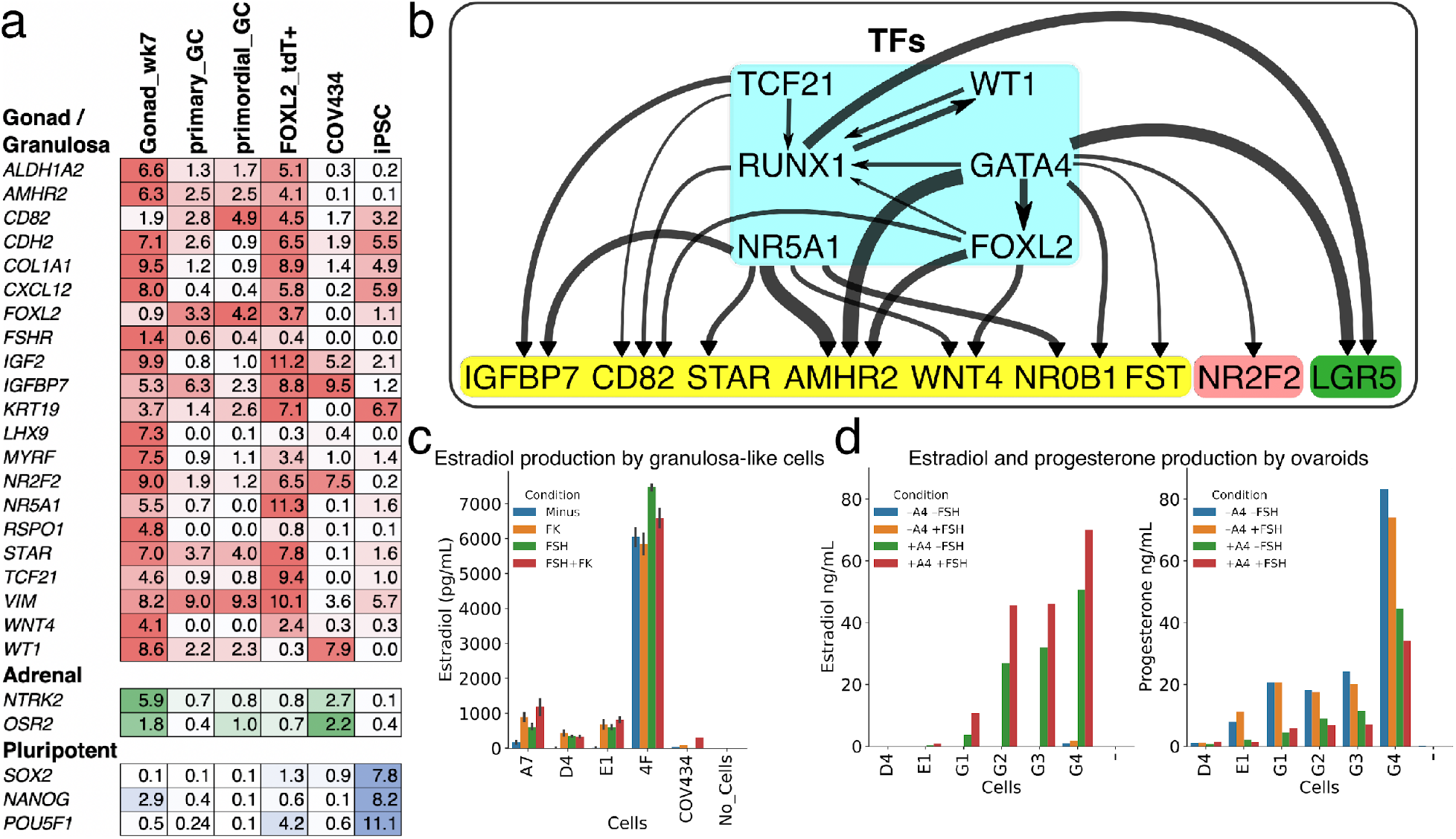
Phenotypic validation of TF-induced granulosa-like cells. **(A)** Gene expression in granulosa-like cells. Log_2_(TPM) values for gondal/granulosa, adrenal, and pluripotent marker genes were compared between week 7 fetal gonad, primary and primordial granulosa cells, TF-induced FOXL2+ cells, COV434 cells, and hiPSCs. **(B)** Regulatory effects of granulosa-related TFs. Expression of TFs (in blue box) was induced by 2 days of doxycycline treatment in hiPSCs, after which time RNA was harvested and sequenced. A differential gene expression analysis was performed for all samples (n = 2 biological replicates per TF) relative to the hiPSC control. Black arrows represent significant (FDR < .05) upregulation, with the width proportional to the log_2_-fold change. Interactions are shown between TFs (blue box) and granulosa marker genes (yellow box), as well as the stromal/theca marker *NR2F2* (red box) and the pre-granulosa marker *LGR5* (green box). **(C)** Granulosa-like cells produce estradiol in the presence of androstenedione and either FSH or forskolin (FK). Results are shown from four monoclonal populations of granulosa-like cells (n = 2 biological replicates for each of 4 clones, error bars are 95% CI), as well as the COV434 cancer cell line. **(D)** Ovaroids produce both estradiol and progesterone. Estradiol production requires androstenedione and is stimulated by FSH. Results are shown for ovaroids formed with six different monoclonal samples of granulosa-like cells (n = 1 sample per ovaroid per condition), at 3 days post-aggregation.

To further elucidate the regulatory effects of individual TFs that we identified as granulosa-related (FOXL2, WT1, NR5A1, GATA4, TCF21, RUNX1), we integrated their cDNA plasmids into hiPSCs and induced expression with doxycycline. For these experiments, we harvested RNA after 2 days of TF expression, since we were interested in short-term effects. We performed differential gene expression analysis relative to hiPSCs, and examined regulatory effects among TFs, and between TFs and marker genes (Figure 2B). Overall, we observed that FOXL2, NR5A1, GATA4, and RUNX1 had the greatest effects on granulosa marker genes. Therefore we generated additional hiPSC lines by integrating these TFs. To rule out any effects of the T2A-tdTomato reporter allele potentially perturbing FOXL2 function, we generated these in a wild-type background (F66 hiPSC line). We measured surface marker expression by flow cytometry after induction, confirmed expression of the granulosa marker *AMHR2* by qPCR, and additionally measured estradiol production in presence of FSH and androstenedione (Figure S3B). We selected the clones (A7, D4, E1) that showed the best overall phenotype.

In the differential expression analysis, we additionally found that GATA4 upregulated *NR2F2,* a marker of ovarian stromal and theca cells. *LGR5,* previously reported as a marker of pre-granulosa cells^16^, was upregulated by GATA4 and RUNX1. We also confirmed a previous report that FOXL2 expression upregulated the cyclin-dependent kinase inhibitor *CDKN1B*^28^, as well as downregulating *ORC1* and *MYC* (Supplementary File 2), suggesting suppression of cellular proliferation.

Finally, we conducted a gene ontology enrichment analysis of upregulated and downregulated genes for each TF^29^. Significantly enriched terms were mainly related to generic developmental processes (e.g. “tissue development”, see Supplementary File 3) although terms related to gonad development were also significantly enriched for NR5A1-upregulated genes.

### Granulosa-like cells respond to FSH and perform steroidogenesis

We next validated the ability of our granulosa-like cells to carry out one of the key endocrine functions of granulosa cells: the production of estradiol. In the ovary, theca cells convert cholesterol to androstenedione, which is the substrate for estradiol production in granulosa cells. The rate-limiting step is oxidative decarboxylation by CYP19A1 (aromatase), producing estrone, which is subsequently reduced to estradiol by enzymes in the HSD17B family, typically HSD17B1 in granulosa cells^30^. *In vivo,* this pathway of estrogen synthesis is stimulated by FSH^31,32^.

We treated our granulosa-like cells with androstenedione, in the presence or absence of FSH or forskolin (which directly increases levels of the FSHR second messenger cAMP). As a control, we used COV434 cells, which were previously reported to produce estradiol from androstenedione^25,26^. Our granulosa-like cells produced estradiol from androstenedione, and in three out of the four monoclonal lines we tested, this steroidogenic activity was dependent on stimulation with FSH or forskolin (Figure 2C). The fourth produced high levels of estradiol in all conditions, and this line lacked the *RUNX1* expression vector that was present in the others. In contrast, COV434 cells, which showed no *FSHR* expression in our RNA-seq data (Figure 2A), were unresponsive to FSH alone, producing estradiol only in the presence of forskolin.

We also investigated whether our granulosa-like cells maintained their steroidogenic activity during ovaroid co-culture with hPGCLCs. We measured hormone levels in ovaroid supernatants in the presence or absence of androstenedione and FSH. In addition to estradiol, we also measured progesterone, which granulosa cells produce *in vivo* after ovulation and formation of the corpus luteum. We observed production of both hormones in five out of six samples (Figure 2D). Estradiol was produced only in the presence of androstenedione supplementation, and levels increased with FSH treatment. Progesterone was produced in all conditions, but was highest in the absence of androstenedione.

### Granulosa-like cells support germ cell maturation within ovaroids

Current methods for inducing and culturing human PGC-like cells (hPGCLCs) produce cells corresponding to immature, premigratory PGCs that lack expression of mature markers such as DAZL^13^. During fetal development, PGCs mature through interactions with ovarian somatic cells, with DAZL playing a key role in downregulation of pluripotency factors and commitment to gametogenesis^33^. This process has recently been recreated *in vitro* using mouse fetal ovarian somatic cells^2^, which allowed the maturation of hPGCLCs to the oogonia-like stage. We hypothesized that *in vitro*-derived human granulosa-like cells could perform a similar role, with the potential for eliminating interspecies developmental mismatches. Therefore, we combined our granulosa-like cells with hPGCLCs to form ovaroids.

Following a previously described protocol (see Methods for details) we aggregated these two cell types in low-binding U-bottom wells, followed by transfer to air-liquid interface Transwell culture. By immunofluorescence, we observed expression of the mature marker DAZL beginning in a subset of OCT4+ hPGCLCs at 4 days of culture (Figure 5A). This is notable in comparison to a previous study using the same hPGCLC line and anti-DAZL antibody, in which DAZL staining was observed only after 77 days of co-culture with mouse fetal testis somatic cells^34^. Expression of TFAP2C, an early-stage PGC marker, declined during ovaroid culture (Figure 5B) and was almost entirely absent by day 8. By contrast, hPGCLCs continued expression of SOX17, OCT4, and DAZL (Figures 3A and 3B). At later time points (day 8 and especially day 16), DAZL+ OCT4– cells were also apparent (Figures 3A and S5). The downregulation of OCT4 in DAZL+ oogonia occurs *in vivo* during the 2^nd^ trimester of human fetal ovarian development^35^; however, we did not observe the transition of DAZL to exclusively cytoplasmic localization that was reported to take place at this stage. Although this system allowed the rapid maturation of hPGCLCs, the number of germ cells declined over prolonged culture, indicating that either they were dying or differentiating to other lineages. By 35 days of culture no germ cells remained, but we observed the formation of empty follicle-like structures composed of cuboidal AMHR2+ FOXL2+ granulosa-like cells (Figure 3C), suggesting that the TFs could drive folliculogenesis even in the absence of oocytes. Follicle-like structure formation was first visible at day 16 (Figure S5), and at day 26 the largest of these structures had grown to 1–2 mm diameter (Figure 3D).

**Figure 3.**
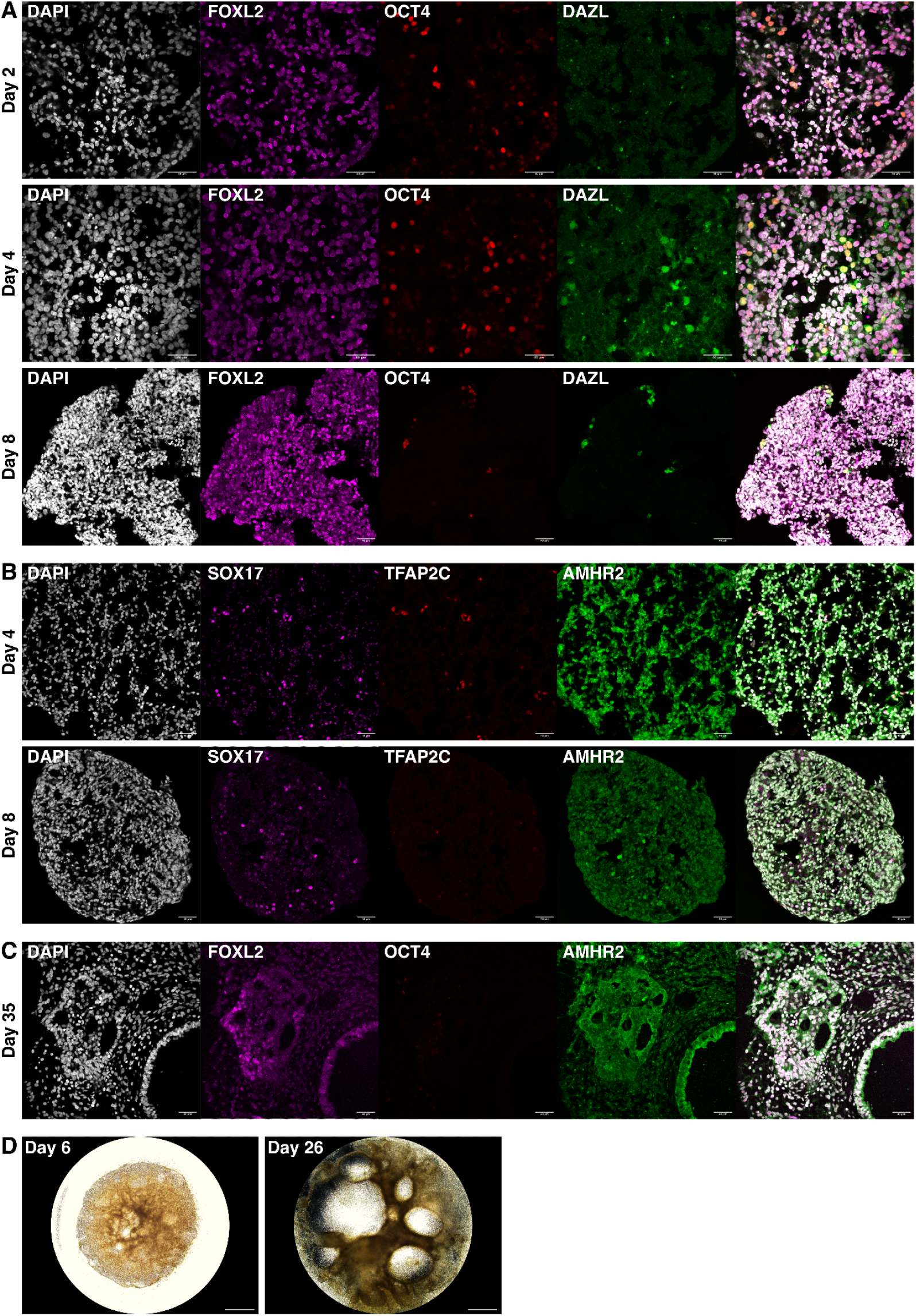
Ovaroid development and follicle formation. **(A)** Ovaroid sections stained for FOXL2 (granulosa), OCT4 (germ cell/pluripotent), and DAZL (mature germ cell). **(B)** Sections stained for SOX17 (germ cell), TFAP2C (early germ cell), and AMHR2 (granulosa). **(C)** Day 35 ovaroid sections stained for FOXL2, OCT4, and AMHR2. All scale bars are 40 μm. **(D)** Whole-ovaroid view of follicle-like structures. Scale bars are 1 mm.

To further examine the gene expression of hPGCLCs and somatic cells in this system, we performed scRNAseq on dissociated ovaroids at days 2, 4, 8, and 14 of culture, and clustered cells according to gene expression. As expected, the largest cluster contained cells expressing granulosa markers such as *FOXL2* and *WNT4* (Figure 4A). Cells expressing markers of secondary/antral granulosa cells such as *FSHR* and *CYP19A1* were also found within this cluster, although these were much less numerous. Full expression data for each cluster is given in Supplementary File 4.

**Figure 4:**
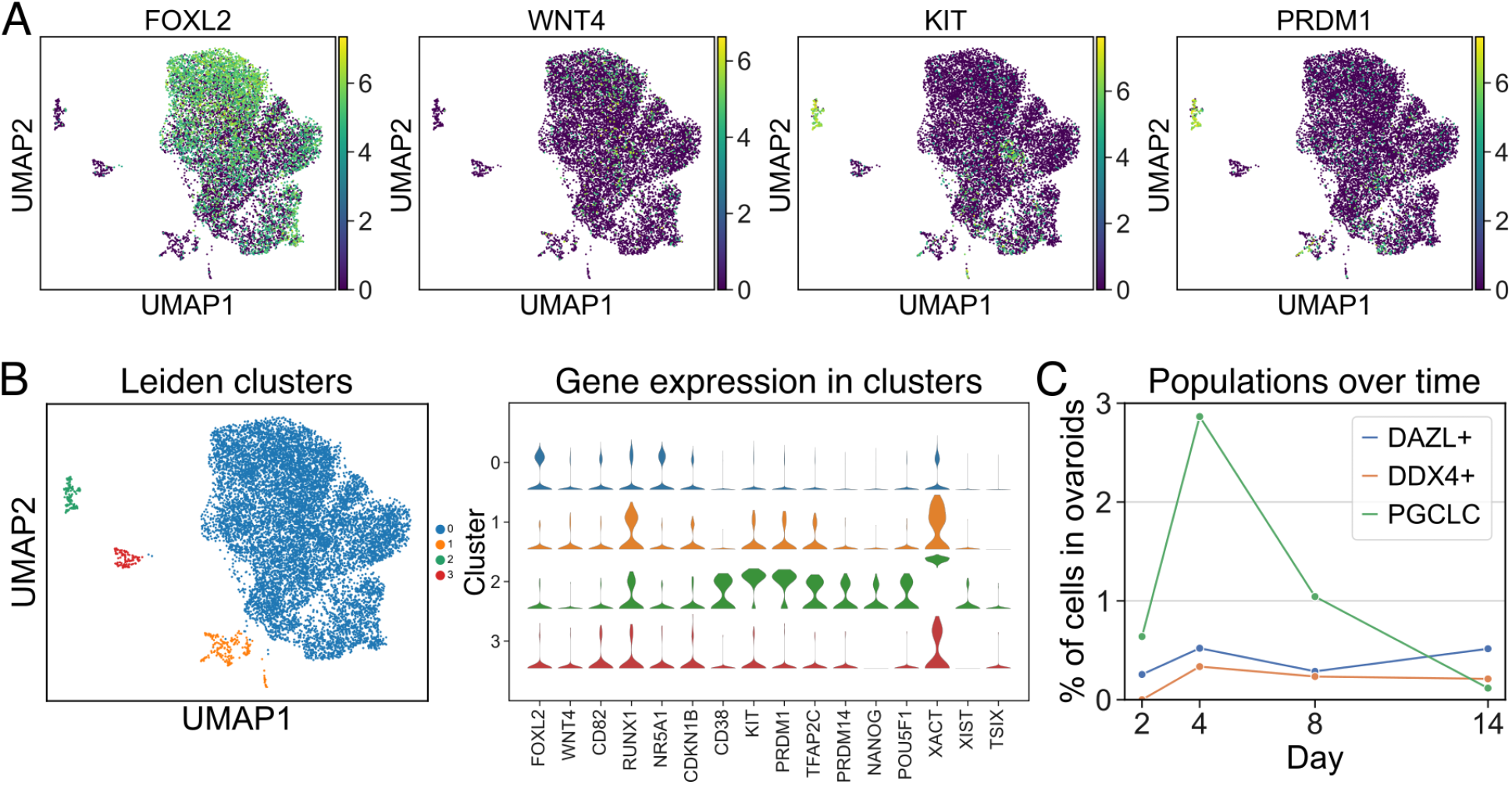
scRNAseq analysis of ovaroids. Data from all samples (days 2, 4, 8, and 14) were combined for joint dimensionality reduction and clustering. **(A)** Expression (log_2_ CPM) of selected granulosa *(FOXL2, WNT4)* and germ cell *(KIT, PRDM1)* markers. **(B)** Leiden clustering shows four main clusters; the expression (log_2_ CPM) of marker genes is plotted for each. **(C)** Proportion of hPGCLCs (cluster #2), *DAZL+* cells, and *DDX4*+ cells in ovaroids from each day.

We also observed a cluster of hPGCLCs expressing marker genes such as *CD38, KIT, PRDM1, TFAP2C, PRDM14, NANOG,* and *POU5F1*. Notably, X-chromosomal lncRNAs *XIST, TSIX,* and *XACT* were all more highly expressed (an average of ~80-fold, ~20-fold, and ~2900-fold, respectively) in the hPGCLCs relative to other clusters (Figure 4B), suggesting that the hPGCLCs were starting the process of X-reactivation^36^, which in hPGCs is associated with high expression of both *XIST* and *XACT*^37^. The X-chromosomal *HPRT1* gene, known to be more highly expressed in cells with two active X chromosomes^38^, was also ~3-fold upregulated (Supplementary File 4).

A small number of *DAZL-*positive cells were also detected by scRNAseq. Interestingly, these cells did not cluster with the hPGCLCs, and lacked expression of pluripotency markers such as *POU5F1* and *NANOG.* There were also a small number of cells expressing a more advanced germ cell marker *DDX4* in samples from day 4 and beyond. Overall, the fraction of hPGCLCs within ovaroids increased from days 2 to 4, but declined thereafter, whereas the fraction of *DAZL+* and *DDX4+* cells also increased from days 2 to 4, but remained roughly constant from days 4 to 14 (Figure 6C).

## Discussion

This work establishes a rapid and efficient method to produce granulosa-like cells by TF-mediated differentiation of hiPSCs. We demonstrate that these granulosa-like cells express granulosa marker genes and can model several key ovarian phenotypes, including hormonal signaling, germ cell maturation, and follicle formation. Additionally, we characterize the regulatory effects of granulosa-related TFs and determine that co-expression of NR5A1 and RUNX1 or RUNX2 is necessary and sufficient to generate granulosa-like cells from hiPSCs. We anticipate that this system will prove useful as a model for the study of ovarian development and function, and that further optimization could allow for human *in vitro* oogenesis.

Our system has several advantages in comparison with existing *in vitro* models of the human ovary. Several studies have used mouse gonadal cells to support hPGCLC maturation^2,34^, but this process is much slower (~77 days vs. ~4 days in our protocol) and the mouse cells may not behave the same as human cells. Other studies have reported growth-factor based protocols to differentiate human pluripotent stem cells to ovarian granulosa-like cells^8–10^. However, the efficiency of such protocols is low (4–12%, or not reported) and those studies did not demonstrate that the cells could support germ cell maturation. Using monoclonal hiPSC lines, our induction efficiency is sufficient to form ovaroids without the need for sorting FOXL2+ cells. Finally, many studies have used cancer cell lines such as COV434 to study various aspects of granulosa cells^25,26^. However, these cancer cells may be phenotypically aberrant. In particular, we show that COV434 cells do not express granulosa marker genes and do not respond to FSH. This is in agreement with a recent study reporting that the COV434 cell line was mis-identified and actually originates from small cell carcinoma of the ovary, not a granulosa tumor^27^. These results highlight the need for improved *in vitro* models of granulosa cells.

While our results represent a step forward for the *in vitro* study of granulosa cells and germ cell maturation, future research will be necessary to construct a full model of the human ovary. Although we can efficiently generate granulosa-like cells and form ovaroids in combination with germ cells, improved methods will be required for maintaining these ovaroids in culture and allowing their development to later stages by preventing the death and/or off-target differentiation of DAZL+ germ cells. Additionally, many important processes in ovarian development, such as sex determination, take place in the early gonad before the differentiation of granulosa cells. Studying these processes will require generating cells at a developmentally earlier stage than what our system produces. Finally, the ovary consists of more cell types than merely germ cells and granulosa cells, and these other cells are also functionally important. For example, our granulosa-like cells produced estradiol only in the presence of androstenedione, which is normally produced by ovarian theca cells. Thus, developing a method to generate theca cells (or earlier-stage gonadal cells that differentiate into both granulosa and theca cells) will provide a more complete model of ovarian biology. We anticipate that our overall approach of identifying TFs to drive the differentiation of hiPSCs will be broadly applicable to generate these cell types of interest, as well as many others.

## Materials and methods

### Cell culture

Two parental hiPSC lines were used in this study: ATCC-BXS0116 female hiPSCs, which we refer to as the F3 line, and the F66 line, an in-house hiPSC line derived from the NIA Aging Cell Repository fibroblast line AG07141 using Epi5 footprint-free episomal reprogramming. The karyotypes of parental lines, as well as engineered reporter lines, were verified by Thermo Fisher Cell ID + Karyostat, and pluripotency was assessed by Thermo Fisher Pluritest. All lines were identified as normal.

hiPSCs were cultured in mTESR Plus medium (Stemcell Technologies) on standard polystyrene plates coated with hESC-qualified Matrigel (Corning). Medium was changed daily. Passaging was performed using 0.5 mM EDTA, or TRYPLE for experiments requiring single-cell dissociation. hiPSCs were treated with 10 μM Y-27632 (Ambeed) for 24 hours after each passage. COV434 cells were cultured in DMEM + 10% FBS + 1X GlutaMax (Gibco). Passaging was performed with TRYPLE (Gibco). hPGCLCs were cultured in S-CM medium as previously described^34^, and passaged with Accutase (Stemcell Technologies). Mycoplasma testing was performed by PCR every 3 months; all cells tested negative.

### Electroporations

Electroporations were performed using a Lonza Nucleofector with 96-well shuttle, with 200,000 cells in 20 μL of P3 buffer. Pulse setting CA-137 was used for all electroporations. Selection with the appropriate agent was begun 48 hours after electroporation and continued for 5 days. For the agents used in this study, this time was sufficient to give a high-purity final cell population.

### Reporter construction

Homology arms for *FOXL2* were amplified by PCR from genomic DNA. A targeting plasmid, containing an in-frame C-terminal T2A-tdTomato reporter, as well as a Rox-PGK-PuroTK-Rox selection cassette (Figure S1A), was constructed by Gibson assembly. The plasmid backbone additionally had an MC1-DTA marker to select against random integration. sgRNA oligos targeting the C-terminal region of *FOXL2* were cloned into pX330 (Addgene #42230). For generation of the reporter lines, 1 μg donor plasmid and 1 μg sgRNA plasmid were co-electroporated into hiPSCs, which were subsequently plated in one well of a 6-well plate. After selection with puromycin (400 ng/mL), colonies were picked manually with a P20 pipette. The hiPSC lines generated were genotyped by PCR for the presence of wild-type and reporter alleles. Homozygous clones were further verified by PCR amplification of the entire *FOXL2* locus (Figure S1B) and Sanger sequencing.

To excise the selection cassette, hiPSCs were electroporated with pCAGGS-Dre (1 μg). Selection was performed with ganciclovir (4 μM) and colonies were picked as described above. The excision of the selection cassette was verified by genotyping. Primers used in this study are listed in Supplementary File 1.

### TF plasmid construction

TF cDNAs were obtained from the TFome^11^ or the ORFeome^39^ as Gateway entry clones. These were cloned into a barcoded Dox-inducible expression vector (Addgene #175503) using MegaGate cloning^40^. The final expression constructs were verified by Sanger sequencing, which also served to determine the barcode sequences for each TF. Two unique barcodes were used per TF during library pooling.

### TF screening for granulosa differentiation

A pooled library of barcoded TF plasmids was electroporated into FOXL2-tdTomato reporter hiPSCs, typically at 5 fmol library and 500 ng PiggyBac transposase expression plasmid (Systems Bio). These conditions were chosen to give an average copy number of approximately 5/cell. Some experiments were also performed at 50 fmol to explore the effects of higher copy numbers. For the screening data presented in Figure 1, two libraries were used: library B, containing 18 TFs, and library C, containing 35 TFs. Library B was used at both 5 fmol and 50 fmol, whereas library C was used only at 5 fmol.

After selection with puromycin (400 ng/mL), hiPSCs were treated with doxycycline (1 μg/mL) in mTESR Plus medium. In additional experiments, hiPSCs were first differentiated to mesoderm following a previously published protocol^41^ before doxycycline treatment. In both sets of experiments, doxycycline treatment continued for 5 days, after which the cells were dissociated with TRYPLE and reporter-positive cells were isolated by FACS. Genomic DNA was extracted (QIAamp DNA Micro kit) from reporter-positive and negative cells, as well as from the initial population before doxycycline treatment.

Barcodes were amplified by PCR (KAPA polymerase), using 10 ng input gDNA per reaction and typically 22 PCR cycles (95°C 15 sec. denature, 58 °C 20 sec. anneal/extend). PCR products were purified using ProNex beads, and a second round of PCR (NEB Q5 polymerase, 6 cycles of 98 °C 5 sec. denature, 61 °C 20 sec. anneal, 72 °C 5 sec. extend, final extension 72 °C 2 min) was performed to add Illumina indices. (Primers are given in **Table S1**). These amplicons were again purified using ProNex beads. Samples were normalized and pooled, and barcodes were sequenced on an Illumina MiSeq with 10% PhiX spike-in. To call barcodes, reads were aligned to the set of known barcode sequences.

### Flow cytometry / Cell sorting

Cells were dissociated by treatment with TRYPLE for 5 minutes, which was quenched with 4 volumes of ice-cold DMEM + 10% FBS. The suspension was passed through a 70 μm cell strainer. Cells were pelleted (200 g, 5 min) and resuspended in staining buffer (PBS + 3% FBS + antibodies, approx. 100 μL per million cells). Staining continued on ice in the dark for 30 minutes. The suspension was diluted with 9 volumes of PBS + 3% FBS. Cells were pelleted (200 g, 5 min) and resuspended in PBS + 3% FBS + 100 ng/mL DAPI. The suspension was kept on ice in the dark until analysis. Flow cytometry was performed on a BD LSRFortessa, and sorting was performed on a Sony SH800 with 100 μm chip.

Antibody capture beads (BD Biosciences, RRID AB_10051478), or hiPSCs expressing tdTomato, were used as compensation controls. Antibodies used are given in **Table S2**. Data analysis was performed using the Cytoflow Python package (version 1.0.0, https://github.com/cytoflow/cytoflow)

### Optimized protocol for granulosa differentiation

iPSCs were dissociated with TRYPLE, and plated in DK10 medium (DMEM-F12, 15 mM HEPES, 1X GlutaMax, 10% KSR) with Y-27632 (10 μM), CHIR99021 (3 μM), and doxycycline (1 μg/mL) at a cell density of 12,500/cm^2^ on Matrigel-coated polystyrene plates. For 24-well plates the medium volume per well was 0.5 mL; for 6-well plates it was 2 mL. 48 hours after plating, the medium was changed to DK10 + doxycycline (1 μg/mL), and the medium was subsequently changed every 24 hours. Cells were harvested on day 5 unless otherwise indicated. In the no-TF control differentiation for RNA-seq, the protocol was the same except the cells did not contain TF expression plasmids.

### RNA-seq

Total RNA was extracted from sorted FOXL2+ granulosa-like cells using the Arcturus PicoPure kit (Thermo Fisher), or from COV434 cells and hiPSCs using the Monarch Total RNA Miniprep kit (NEB). For experiments involving TF overexpression, TF expression plasmids were integrated into hiPSCs as described above (50 fmol / 200,000 cells). After selection with puromycin, TF expression was induced using doxycycline (1000 ng/mL).

Two biological replicates were collected for each sample (iPSC, hiPSC + individual TFs, sorted FOXL2+, no-TF differentiation, COV434). Libraries were prepared using the NEBNext Ultra II Directional kit following the manufacturer’s protocol, and sequenced on an Illumina NextSeq 500 (2 x 75bp paired-end reads). The TPM data shown in Figure 3 were generated using kallisto^42^ to pseudoalign reads to the reference human transcriptome (Ensembl GRCh38 v96). Differential expression analysis was performed using DESeq2. PantherDB^29^ was used to calculate gene ontology enrichment for significantly upregulated (log_2_fc > 3, p_adj_ < 0.05) and downregulated (log_2_fc < −3, p_adj_ < 0.05) genes for each sample relative to hiPSCs.

### Ovaroid formation with hPGCLCs and granulosa-like cells

F2 female hPGCLCs were maintained in long-term culture medium as previously described^34^ and harvested with Accutase. To form ovaroids, granulosa-like cells were harvested with TRYPLE, counted, and mixed with F2 hPGCLCs. For each ovaroid, 100,000 granulosa-like cells and 10,000 hPGCLCs were added to each well of a 96-well U-bottom low-bind plate (Corning #7007) in 200 μL of GK15 medium (GMEM, 15% KSR, with 1X GlutaMax, sodium pyruvate, and non-essential amino acids) supplemented with 10 mM Y-27632, 0.1 mM 2-mercaptoethanol, 1 μg/mL doxycycline, 100 ng/mL SCF, and 50 μg/mL primocin. The plate was centrifuged (100 g, 2 min) and incubated (37 °C, 5% CO_2_) for two days. Subsequently, ovaroids were transferred to Transwells (collagen-coated PTFE, 3 μm pore size, 24 mm diameter, Corning #3492) for air-liquid interface culture with aMEM, 10% KSR, 55 μM 2-mercaptoethanol, 1 μg/mL doxycycline, and 50 μg/mL primocin. Typically 5-6 ovaroids were cultured on each 6-well Transwell. The medium (1.5 mL) was changed every 2 days.

### Immunofluorescence

Ovaroids were washed with PBS and fixed with 1% PFA overnight at 4°C. After another PBS wash, ovaroids were detached from the Transwell. In preparation for cryosectioning, ovaroids were transferred to 10% sucrose in PBS. After 24 hr. at 4 °C, the 10% sucrose solution was removed and replaced with 20% sucrose in PBS. After an additional 24 hr. at 4 °C, the ovaroids were embedded in OCT compound and stored at −80 °C until sectioning.

The ovaroids were sectioned to 10 μm using a Leica CM3050S cryostat. Sections were transferred to Superfrost Plus slides, which were washed with PBS to remove OCT compound. The slides were washed with PBST (0.1% Triton X-100 in PBS) and sections were circled with a Pap pen. Slides were blocked for 30 min. at room temp. with blocking buffer (1% bovine serum albumin and 5% normal donkey serum [Jackson ImmunoResearch, #017-000-121, lot #152961] in PBST). The blocking buffer was removed and replaced with a solution of primary antibodies in blocking buffer, and the slides were incubated overnight at 4 °C. The antibody solution was removed and the slides were washed with PBST for 3 x 5 min. The slides were incubated with secondary antibody and DAPI solution in blocking buffer for 1 hr. at room temp. in the dark, followed by two 5-minute washes with PBST and one wash with PBS. After staining, samples were mounted in Prolong Gold medium and covered with coverslips. Imaging was performed on a Leica SP5 confocal microscope. Antibodies used are given in **Table S2.**

### scRNAseq

Ovaroids (6 ovaroids per sample, 2 samples per time point) were dissociated using the Miltenyi Embryoid Body Dissociation Kit (Miltenyi #130-096-348). The cells were passed through a 40 μm strainer, fixed using the Parse Biosciences fixation kit, and stored at −80 °C until all time points had been collected. Libraries were prepared using the Parse Biosciences WT Mega kit generating libraries of an average of 450bp. The ovaroids took up 8 of the 96 samples; the remaining kit capacity was used for other experiments. The libraries were sequenced on an Illumina NovaSeq 2×150bp S4 flow cell using single index, 6bp, libraries and a 5% PhiX spike-in. Data was demultiplexed into library fastq files and counts matrices were generated using Parse Bioscience’s analysis pipeline (version 0.9.6). Downstream data processing, such as doublet filtering, dimensionality reduction, and clustering, was performed using scanpy (version 1.8.2)^43^.

### Steroid hormone assays

Androstenedione (500 ng/mL) was added to the medium on day 4 of granulosa differentiation. FSH (0.25 IU/mL, BioVision #4781-50 lot 5F07L47810) or forskolin (100 μM, Sigma-Aldrich) were also added as indicated. The total medium volume was 0.5 mL per well of 24-well plate. After 24 hours, the medium was analyzed for estradiol content by ELISA (DRG International, EIA-2693). Concentrations were calculated with a 4-parameter logistic curve fit using the data from the standards provided in the kit. Samples outside the range of the calibration curve were diluted and re-run.

For measuring hormone production in ovaroids, ovaroids were aggregated as described above. Androstenedione (500 ng/mL) and/or FSH (0.25 IU/mL) were added to the aggregation medium (total volume 200 μL per ovaroid). After 3 days of culture, the medium was removed and analyzed by ELISA for estradiol (DRG International, EIA-2693) and progesterone (DRG International, EIA-1561). Hormone concentrations were calculated as described above.

## Supporting information

Supplement 1: Oligos

Supplement 2: Bulk RNA-seq Gene Expression

Supplement 3: GO Enrichment

Supplement 4: Cluster DGE

## Data and code availability

All data needed to evaluate the conclusions in the paper are present in the paper and supplementary tables and figures. Data analysis code can be found at: https://github.com/programmablebio/granulosa. Raw and processed sequencing data will be deposited to GEO upon publication.

## Author Contributions

M.P.S. and C.K. conceived, designed, and directed the study. M.P.S. performed reporter cell line creation, TF cloning, cell engineering, TF screening, ovaroid formation, steroidogenesis assays, confocal microscopy, scRNA-sequencing, NGS, and manuscript writing. C.K. performed TF cloning, scRNA-sequencing and data processing, hPGCLC culture, barcoded library generation, NGS library prep and sequencing, and creation of F66 hiPSC line. P.R.F. aided in confocal imaging and library preparation. P.C., E.T. and G.B. developed and executed TF prediction algorithms, and single cell and bulk analysis pipelines. M.K. and T.S aided in germ cell methodologies, E.D., and J.A., assisted in microscopy sample preparation and confocal imaging. P.C., R.E.K., T.S., and G.M.C supervised the study.

## Acknowledgements

This work was funded by the Synthetic Biology Platform at the Wyss Institute for Biologically Inspired Engineering. This work was additionally funded by the Gameto Sponsored Research Agreement at Harvard University and Colossal Sponsored Research Agreement at Harvard University. M.P.S. was supported by the National Science Foundation Graduate Research Fellowship.

## References

1. Yoshino, T. et al. Generation of ovarian follicles from mouse pluripotent stem cells. Science 373, eabe0237 (2021).

2. Yamashiro, C. et al. Generation of human oogonia from induced pluripotent stem cells in vitro. Science 362, 356–360 (2018).

3. Le Bouffant, R. et al. Meiosis initiation in the human ovary requires intrinsic retinoic acid synthesis. Human Reproduction 25, 2579–2590 (2010).

4. Su, Y.-Q. et al. Synergistic roles of BMP15 and GDF9 in the development and function of the oocyte–cumulus cell complex in mice: genetic evidence for an oocyte–granulosa cell regulatory loop. Developmental Biology 276, 64–73 (2004).

5. Peng, J. et al. Growth differentiation factor 9:bone morphogenetic protein 15 heterodimers are potent regulators of ovarian functions. Proceedings of the National Academy of Sciences 110, E776–E785 (2013).

6. Otsuka, F. & Shimasaki, S. A negative feedback system between oocyte bone morphogenetic protein 15 and granulosa cell kit ligand: Its role in regulating granulosa cell mitosis. Proceedings of the National Academy of Sciences 99, 8060–8065 (2002).

7. Findlay, J. K. et al. Follicle Selection in Mammalian Ovaries. in The Ovary (ed. Leung, P.) 1–21 (Elsevier, 2019).

8. Lipskind, S. et al. An Embryonic and Induced Pluripotent Stem Cell Model for Ovarian Granulosa Cell Development and Steroidogenesis. Reprod Sci 25, 712–726 (2018).

9. Lan, C.-W., Chen, M.-J., Jan, P.-S., Chen, H.-F. & Ho, H.-N. Differentiation of Human Embryonic Stem Cells Into Functional Ovarian Granulosa-like Cells. The Journal of Clinical Endocrinology & Metabolism 98, 3713–3723 (2013).

10. Gonen, N. et al. In-vitro *cellular reprogramming to model gonad development and its disorders*. http://biorxiv.org/lookup/doi/10.1101/2021.10.22.465384 (2021) doi:10.1101/2021.10.22.465384.

11. Ng, A. H. M. et al. A comprehensive library of human transcription factors for cell fate engineering. Nat Biotechnol 39, 510–519 (2021).

12. Liang, J. et al. Induction of Sertoli-like cells from human fibroblasts by NR5A1 and GATA4. eLife 8, e48767 (2019).

13. Irie, N. et al. SOX17 Is a Critical Specifier of Human Primordial Germ Cell Fate. Cell 160, 253–268 (2015).

14. Zhang, Y. et al. Transcriptome Landscape of Human Folliculogenesis Reveals Oocyte and Granulosa Cell Interactions. Molecular Cell 72, 1021–1034.e4 (2018).

15. Manuylov, N. L., Smagulova, F. O., Leach, L. & Tevosian, S. G. Ovarian development in mice requires the GATA4-FOG2 transcription complex. Development 135, 3731–3743 (2008).

16. Niu, W. & Spradling, A. C. Two distinct pathways of pregranulosa cell differentiation support follicle formation in the mouse ovary. Proc Natl Acad Sci USA 117, 20015–20026 (2020).

17. Voronina, E. et al. Ovarian granulosa cell survival and proliferation requires the gonad-selective TFIID subunit TAF4b. Developmental Biology 303, 715–726 (2007).

18. Richards, J. S. Perspective: The Ovarian Follicle—A Perspective in 2001. Endocrinology 142, 2184–2193 (2001).

19. Nicol, B. et al. RUNX1 maintains the identity of the fetal ovary through an interplay with FOXL2. Nat Commun 10, 5116 (2019).

20. Kramme, C. et al. An integrated pipeline for mammalian genetic screening. Cell Reports Methods 100082 (2021) doi:10.1016/j.crmeth.2021.100082.

21. Ottolenghi, C. et al. Foxl2 is required for commitment to ovary differentiation. Human Molecular Genetics 14, 2053–2062 (2005).

22. Cocquet, J. Evolution and expression of FOXL2. Journal of Medical Genetics 39, 916–921 (2002).

23. Oktay, K., Briggs, D. & Gosden, R. G. Ontogeny of Follicle-Stimulating Hormone Receptor Gene Expression in Isolated Human Ovarian Follicles. Journal of Clinical Endocrinology and Metabolism 82, 3748–3751.

24. Sybirna, A. et al. A critical role of PRDM14 in human primordial germ cell fate revealed by inducible degrons. Nature Communications 11, (2020).

25. Price, J. C., Cronin, J. & Sheldon, I. M. Toll-Like Receptor Expression and Function in the COV434 Granulosa Cell Line. Am J Reprod Immunol 68, 205–217 (2012).

26. Zhang, H. Characterization of an immortalized human granulosa cell line (COV434). Molecular Human Reproduction 6, 146–153 (2000).

27. Karnezis, A. N. et al. Re-assigning the histologic identities of COV434 and TOV-112D ovarian cancer cell lines. Gynecologic Oncology 160, 568–578 (2021).

28. Gustin, S. E. et al. WNT/β-catenin and p27/FOXL2 differentially regulate supporting cell proliferation in the developing ovary. Developmental Biology 412, 250–260 (2016).

29. Mi, H. et al. PANTHER version 16: a revised family classification, tree-based classification tool, enhancer regions and extensive API. Nucleic Acids Research 49, D394–D403 (2021).

30. Hakkarainen, J. et al. Hydroxysteroid (17β)-dehydrogenase 1-deficient female mice present with normal puberty onset but are severely subfertile due to a defect in luteinization and progesterone production. The FASEB Journal 29, 3806–3816 (2015).

31. Sasson, R. Gonadotrophin-induced gene regulation in human granulosa cells obtained from IVF patients. Modulation of steroidogenic genes, cytoskeletal genes and genes coding for apoptotic signalling and protein kinases. Molecular Human Reproduction 10, 299–311 (2004).

32. Welsh, T. H., Jia, X.-C. & Hsueh, A. J. W. Forskolin and phosphodiesterase inhibitors stimulate rat granulosa cell differentiation. Molecular and Cellular Endocrinology 37, 51–60 (1984).

33. Nicholls, P. K. et al. Mammalian germ cells are determined after PGC colonization of the nascent gonad. Proc. Natl. Acad. Sci. U.S.A. 116, 25677–25687 (2019).

34. Kobayashi, M. et al. Expanding homogeneous culture of human primordial germ cell-like cells maintaining germline features without serum or feeder layers. Stem Cell Reports S221367112200056X (2022) doi:10.1016/j.stemcr.2022.01.012.

35. Anderson, R. A., Fulton, N., Cowan, G., Coutts, S. & Saunders, P. T. Conserved and divergent patterns of expression of DAZL, VASA and OCT4 in the germ cells of the human fetal ovary and testis. BMC Developmental Biology 7, 136 (2007).

36. Vallot, C. et al. XACT Noncoding RNA Competes with XIST in the Control of X Chromosome Activity during Human Early Development. Cell Stem Cell 20, 102–111 (2017).

37. Chitiashvili, T. et al. Female human primordial germ cells display X-chromosome dosage compensation despite the absence of X-inactivation. Nat Cell Biol 22, 1436–1446 (2020).

38. Epstein, C. J. Expression of the Mammalian X Chromosome before and after Fertilization. Science 175, 1467–1468 (1972).

39. The ORFeome Collaboration. The ORFeome Collaboration: a genome-scale human ORF-clone resource. Nat Methods 13, 191–192 (2016).

40. Kramme, C. et al. MegaGate: A toxin-less gateway molecular cloning tool. STAR Protoc 2, 100907 (2021).

41. Morizane, R. & Bonventre, J. V. Generation of nephron progenitor cells and kidney organoids from human pluripotent stem cells. Nat Protoc 12, 195–207 (2017).

42. Bray, N. L., Pimentel, H., Melsted, P. & Pachter, L. Near-optimal probabilistic RNA-seq quantification. Nat Biotechnol 34, 525–527 (2016).

43. Wolf, F. A., Angerer, P. & Theis, F. J. SCANPY: large-scale single-cell gene expression data analysis. Genome Biol 19, 15 (2018).

