## Supplementary Information for "Directed Differentiation of Human iPSCs to Functional Ovarian Granulosa-Like Cells via Transcription Factor Overexpression"

### Supplementary tables:

S1. List of screened TFs
S2. Antibodies used in this study

### Supplementary figures:

S1. Construction of the FOXL2-T2A-tdTomato reporter hiPSC line.
S2. Control of TF expression plasmid copy number delivered to hiPSCs.
S3. Evaluation of monoclonal hiPSC lines for yield and quality of granulosa-like cells.
S4. Monoclonal hiPSC lines with integrated TFs allow the efficient production of granulosa-like cells in response to doxycycline.
S5. DAZL and OCT4 expression observed by immunofluorescence in day 16 ovaroids.

### Supplementary files:

File 1. List of oligonucleotides (.xlsx)
File 2. Bulk RNA-seq gene expression data
File 3. GO term enrichment
File 4. Gene expression in clusters observed in ovaroid scRNAseq

**Supplementary Table 1: List of screened TFs**

| TF | Library | Reason for inclusion |
|---|---|---|
| FOS | B and C | DEG analysis |
| CEBPD | B and C | DEG analysis |
| ELK1 | B and C | DEG analysis |
| FOXL2 | B and C | DEG analysis |
| GATA4 | B and C | DEG analysis |
| JUN | B and C | DEG analysis |
| KLF2 | B and C | DEG analysis |
| NR2F2 | B and C | DEG analysis |
| NR4A1 | B and C | DEG analysis |
| NR5A1 | B and C | DEG analysis |
| TCF21 | B and C | DEG analysis |
| RUNX1 | B and C | Literature |
| ZFPM2 | B and C | Literature |
| EGR1 | B and C | Network analysis |
| KLF4 | B and C | Network analysis |
| PPARG | B and C | Network analysis |
| RUNX2 | B and C | Network analysis |
| YBX1 | B and C | Network analysis |
| ATF4 | C | DEG analysis |
| EMX2 | C | DEG analysis |
| FOSB | C | DEG analysis |
| HOPX | C | DEG analysis |
| HOXC9 | C | DEG analysis |
| JUNB | C | DEG analysis |
| MAFF | C | DEG analysis |
| TSC22D3 | C | DEG analysis |
| WT1 | C | DEG analysis |
| ZBTB16 | C | DEG analysis |
| LHX1 | C | Literature |
| LHX9 | C | Literature |
| TAF4B | C | Literature |
| KLF6 | C | Network analysis |
| MYC | C | Network analysis |
| NR1H2 | C | Network analysis |
| TOX2 | C | Network analysis |

**Supplementary Table 2: Antibodies used in this study**

**Primary antibodies for immunofluorescence**

| Target | Antibody type | Supplier | Cat# | RRID | Dilution |
|---|---|---|---|---|---|
| AMHR2 | Rabbit IgG, polyclonal | Thermo Fisher | PA5-13902 | AB_2305463 | 1:100 |
| DAZL | Rabbit IgG, monoclonal | Abcam | ab215718 | AB_2893177 | 1:500 |
| FOXL2 | Goat IgG, polyclonal | Novus | NB100-1277 | AB_2106187 | 1:250 |
| OCT4 | Mouse IgG, monoclonal | BD Biosciences | 611202 | AB_398736 | 1:200 |
| SOX17 | Goat IgG, polyclonal | Novus | AF1924 | AB_355060 | 1:500 |
| TFAP2C | Mouse IgG, monoclonal | Abcam | ab110635 | AB_10858471 | 1:250 |

**Secondary antibodies for immunofluorescence**

| Target | Fluorophore | Antibody type | Supplier | Cat# | RRID | Dilution |
|---|---|---|---|---|---|---|
| Mouse IgG | AF647 | Donkey IgG | Fisher | A31571 | AB_162542 | 1:250 |
| Goat IgG | AF568 | Donkey IgG | Fisher | A11057 | AB_2534104 | 1:500 |
| Rabbit IgG | AF488 | Donkey IgG F(ab')$_2$ | Jackson | 711-546-152 | AB_2340619 | 1:500 |

**Antibodies for flow cytometry**

| Target | Fluorophore | Antibody type | Supplier | Cat# | RRID | Dilution |
|---|---|---|---|---|---|---|
| AMHR2 | FITC | Rabbit IgG | Biorbyt | orb37457 | AB_10992015 | 1:60 |
| CD82 | PerCP-Cy5.5 | Mouse IgG | BioLegend | 342111 | AB_2750124 | 1:60 |
| EpCAM | APC-Cy7 | Mouse IgG | BioLegend | 324245 | AB_2783193 | 1:60 |
| FSHR | APC | Mouse IgG | R&D Systems | FAB65591A | AB_2920602 | 1:60 |

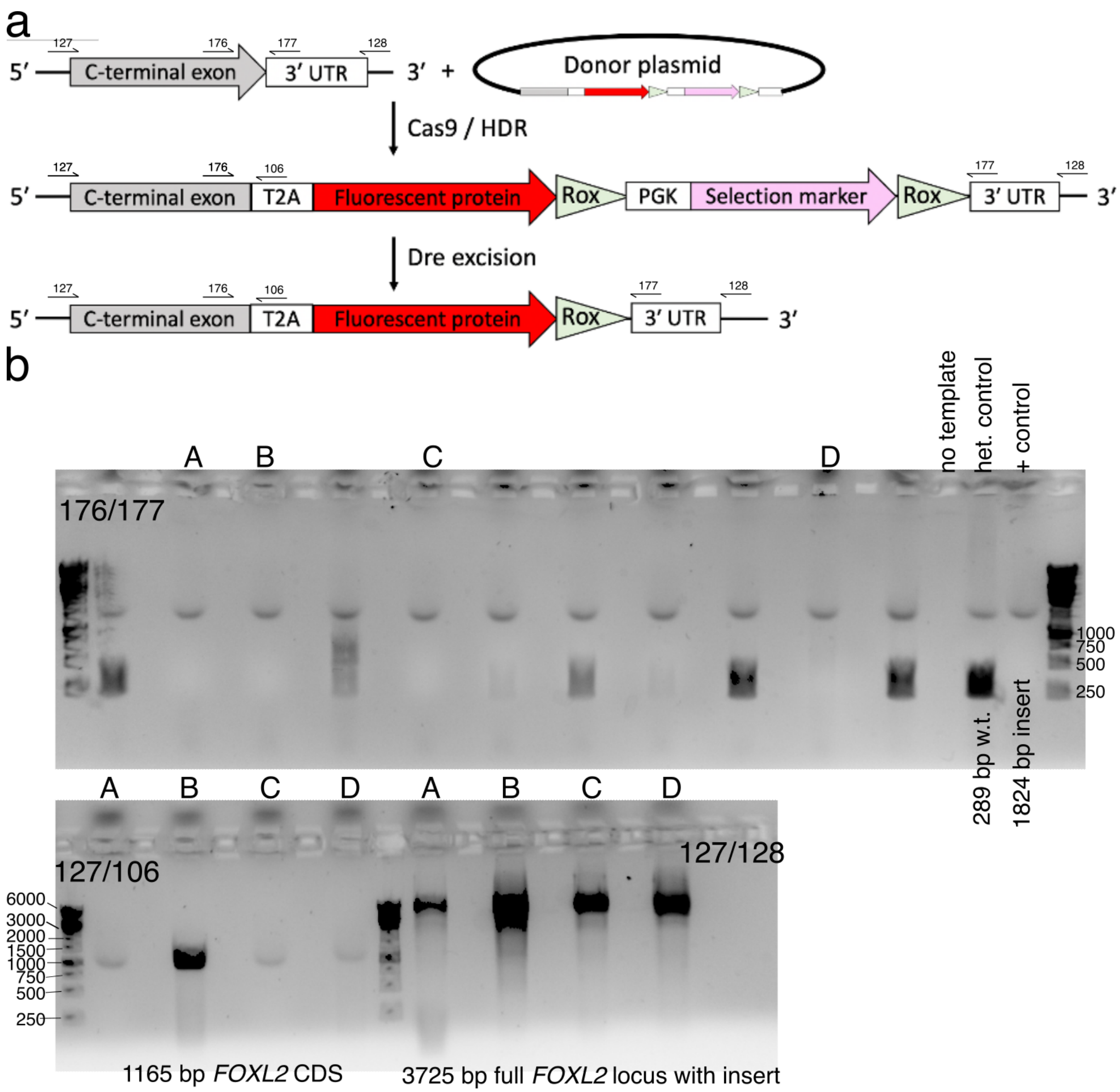


**Figure S1.** Construction of the FOXL2-T2A-tdTomato reporter hiPSC line. **(A)** Schematic for Cas9/HDR knock-in with a donor plasmid followed by selection marker excision. Primer binding sites for genotyping are also shown (not to scale). **(B)** Genotyping to verify homozygous editing. Initial screening was performed with primers 176/177. Candidate clones (denoted A, B, C, D) were further verified by additional genotyping. Note that primers 127/128 bind outside of the region used as homology arms for the donor plasmid.

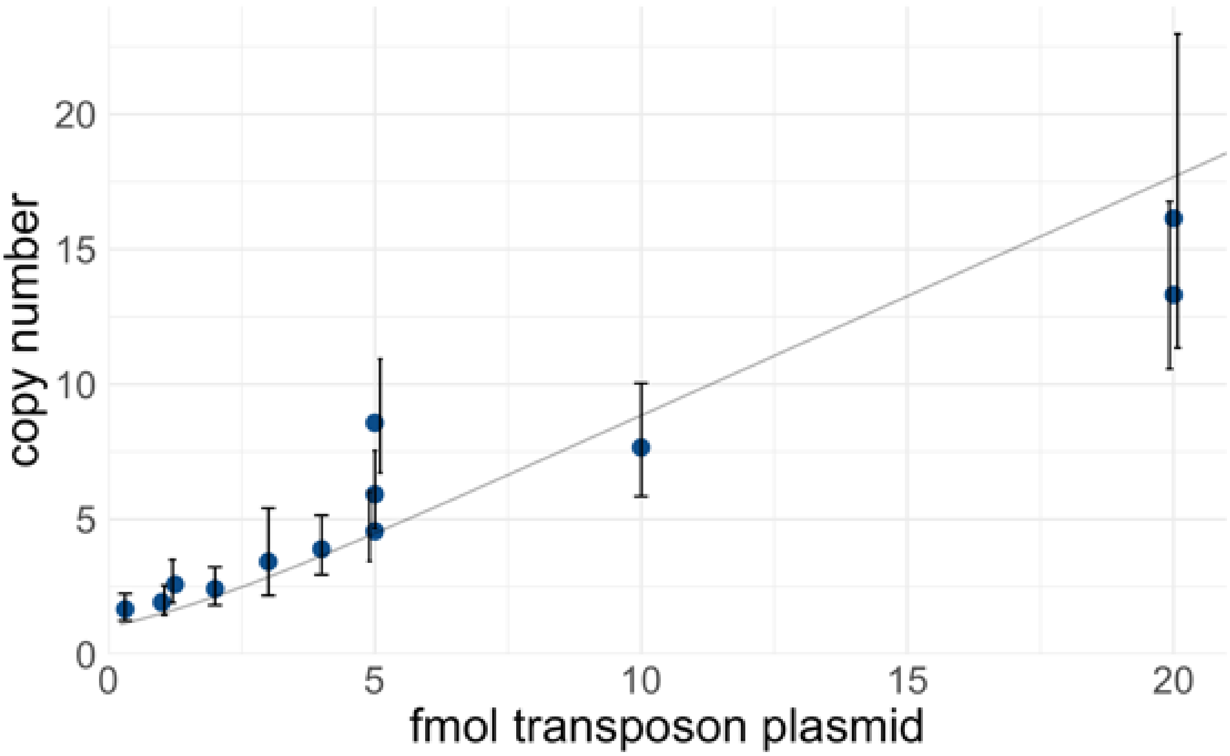


**Figure S2.** Control of TF expression plasmid copy number delivered to hiPSCs. A PiggyBac transposon library of barcoded doxycycline-inducible TF cDNAs was electroporated into iPSCs at varying concentrations. The amount of transposase plasmid was held constant (500 ng). After selection, gDNA was extracted and the transposon copy number was measured by qPCR using primer pairs CK107/108 (amplifies *RPP30* genomic control) and qMPS015 (amplifies *TET3G*, present in all TF expression plasmids). To calculate the copy number, a standard curve was generated using a control plasmid with a known 1:1 ratio of *RPP30* and *TET3G*. Each point represents a biological replicate, and error bars represent 95% confidence intervals of the technical replicates. The curve in light gray is a weighted-least-squares fit of a zero-truncated Poisson mean, which represents the theoretical relationship.

a

| Cells | CD82+ FOXL2+ EPCAM– | EPCAM+ | FSHR+ |
|---|---|---|---|
| 1A | 0.24 | 0.61 | 0.07 |
| 1B | 0.14 | 0.81 | 0.03 |
| 1C | 0.13 | 0.73 | 0.03 |
| 1D | 0.07 | 0.73 | 0.00 |
| 1E | 0.26 | 0.52 | 0.03 |
| 1F | 0.71 | 0.00 | 0.00 |
| 2A | 0.19 | 0.06 | 0.00 |
| 2B | 0.25 | 0.34 | 0.01 |
| 2C | 0.72 | 0.03 | 0.00 |
| 2D | 0.59 | 0.00 | 0.00 |
| 2E | 0.08 | 0.68 | 0.02 |
| 2F | 0.16 | 0.78 | 0.03 |
| 3A | 0.11 | 0.69 | 0.01 |
| 3B | 0.13 | 0.49 | 0.01 |
| 3C | 0.11 | 0.53 | 0.00 |
| 3D | 0.08 | 0.88 | 0.01 |
| 3E | 0.08 | 0.78 | 0.02 |
| 3F | 0.08 | 0.70 | 0.00 |
| 4B | 0.19 | 0.46 | 0.00 |
| 4C | 0.67 | 0.13 | 0.00 |
| 4D | 0.07 | 0.57 | 0.00 |
| 4E | 0.12 | 0.78 | 0.03 |
| 4F | 0.05 | 0.89 | 0.00 |
| No TFs | 0.07 | 0.74 | 0.03 |

b

| Cells | EPCAM– CD82+ | EPCAM+ CD82– | Estradiol ng/mL | AMHR2 ΔΔCt vs GAPDH |
|---|---|---|---|---|
| A1 | 0.25 | 0.24 | 5.03 | -3.7 |
| A2 | 0.17 | 0.10 | 3.67 | |
| A3 | 0.53 | 0.05 | 0.09 | -4.3 |
| A4 | 0.10 | 0.27 | 5.23 | |
| A5 | 0.11 | 0.29 | 4.61 | |
| A6 | 0.03 | 0.41 | 3.37 | |
| **A7** | **0.36** | **0.10** | **2.93** | **-9.2** |
| B1 | 0.53 | 0.08 | 0.09 | -8.7 |
| B2 | 0.08 | 0.30 | 3.00 | |
| B3 | 0.60 | 0.15 | 3.78 | -4.9 |
| B4 | 0.50 | 0.07 | 0.00 | -7.3 |
| B5 | 0.28 | 0.27 | 4.01 | |
| B6 | 0.79 | 0.02 | 0.03 | -3.6 |
| B7 | 0.10 | 0.34 | 4.88 | |
| C1 | 0.23 | 0.22 | 0.57 | |
| C2 | 0.70 | 0.02 | 0.01 | -4.7 |
| C3 | 0.57 | 0.02 | 0.40 | -2.0 |
| C4 | 0.12 | 0.32 | 0.29 | |
| C5 | 0.34 | 0.17 | 0.01 | |
| C6 | 0.08 | 0.21 | 4.04 | -9.3 |
| C7 | 0.22 | 0.22 | 0.39 | |
| D1 | 0.16 | 0.36 | 5.69 | -4.9 |
| D2 | 0.48 | 0.09 | 0.01 | -5.1 |
| D3 | 0.33 | 0.13 | 0.19 | |
| **D4** | **0.71** | **0.00** | **0.02** | **-5.7** |
| D5 | 0.30 | 0.07 | 0.22 | |
| D6 | 0.45 | 0.03 | 0.07 | |
| D7 | 0.46 | 0.01 | 0.00 | |
| **E1** | **0.42** | **0.08** | **2.80** | **-9.0** |
| E2 | 0.25 | 0.31 | 4.37 | |
| E3 | 0.61 | 0.09 | 0.00 | -6.4 |
| E4 | 0.28 | 0.29 | 2.99 | |
| E5 | 0.02 | 0.42 | 4.40 | |
| E6 | 0.05 | 0.36 | 5.29 | |
| E7 | 0.14 | 0.21 | 5.03 | -9.8 |
| No TFs | 0.02 | 0.58 | 0 | **0.0** |

c

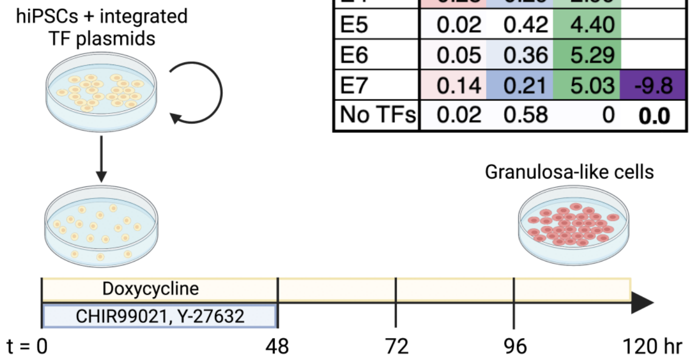


**Figure S3.** Evaluation of monoclonal hiPSC lines for yield and quality of granulosa-like cells. **(A)** Lines originating from the F3/FOXL2-T2A-tdTomato hiPSC reporter line were evaluated by flow cytometry for FOXL2, CD82, EPCAM, and FSHR. **(B)** Lines originating from the F66 wild-type hiPSC line were evaluated by flow cytometry for CD82 and EPCAM, as well as for estradiol production and qPCR to measure *AMHR2* expression (the no-TF control was used as a reference for calculating ΔΔCt). **(C)** Method of inducing granulosa-like cells from hiPSCs.

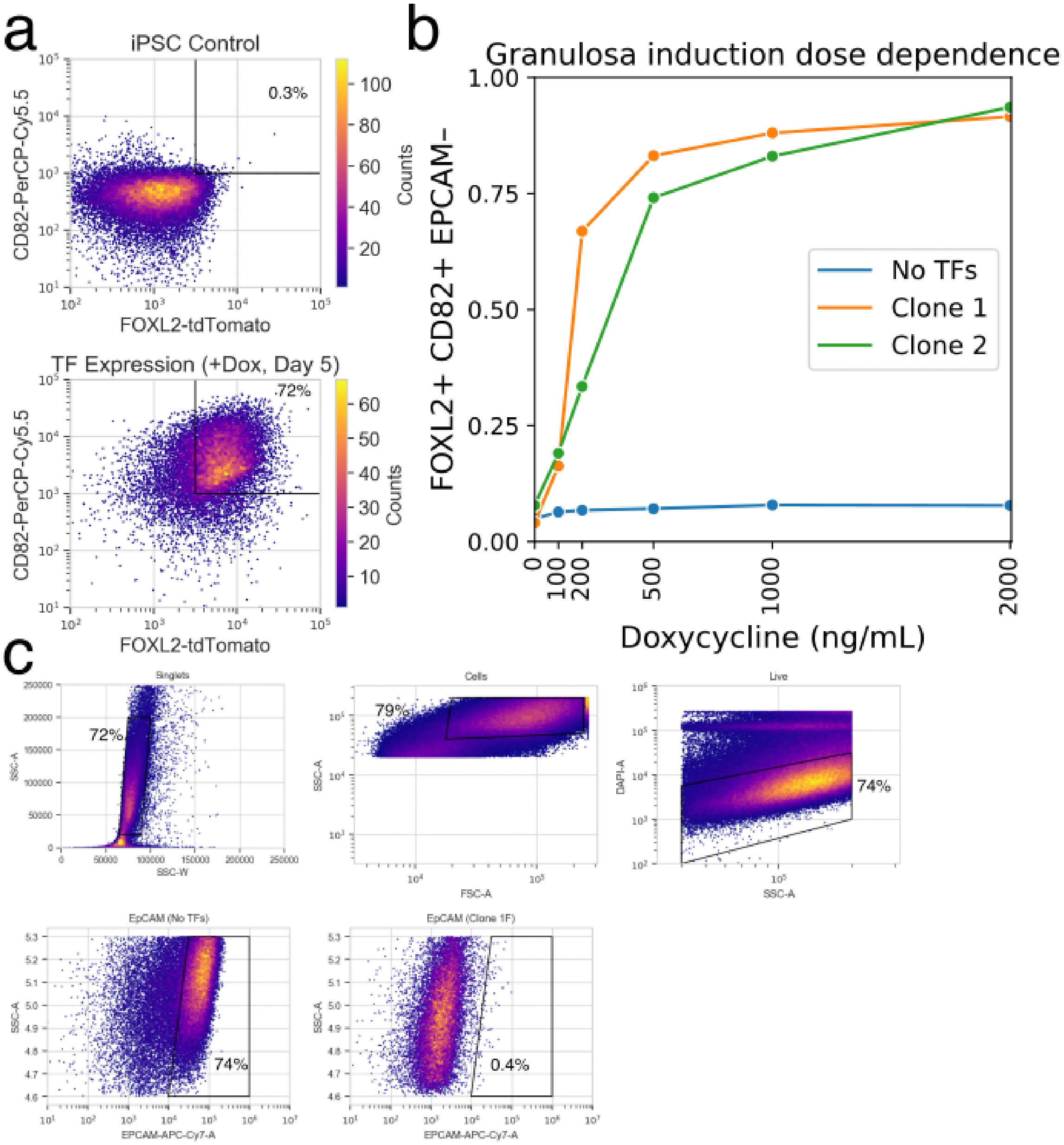


**Figure S4.** Monoclonal hiPSC lines with integrated TFs allow the efficient production of granulosa-like cells in response to doxycycline. **(A)** Flow cytometry showing FOXL2 and CD82 expression after 5 days of TF overexpression with 1000 ng/mL doxycycline. **(B)** Dose-dependence for the production of FOXL2+ CD82+ EPCAM– granulosa-like cells, shown in two monoclonal lines. Clone 1F contains NR5A1, TCF21, and RUNX1 expression vectors, whereas clone 2D has RUNX2 instead of RUNX1. Granulosa-like cells are efficiently induced from both clones (but not from control cells lacking TF expression vectors) in a doxycycline-dependent manner. **(C)** Representative gating strategy to analyze flow cytometry data.

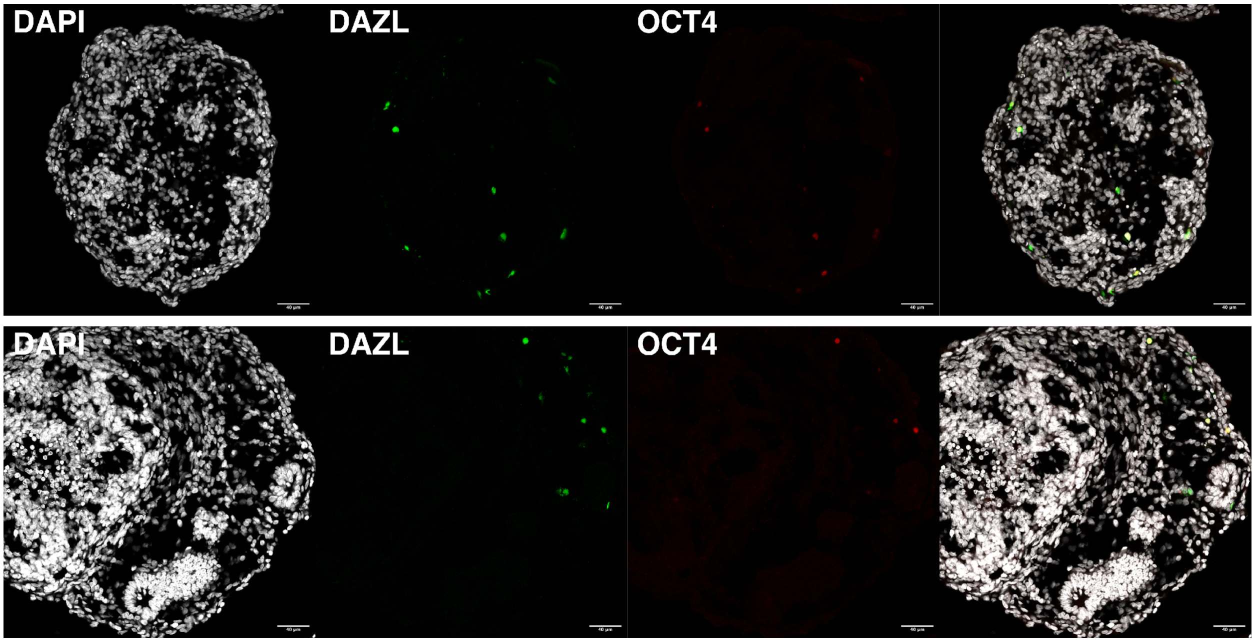


**Figure S5.** DAZL and OCT4 expression observed by immunofluorescence in day 16 ovaroids. Some DAZL+ OCT4– cells are visible, as well as DAZL+ OCT4+ cells. Ovaroids are also beginning to form follicle-like morphology. Scale bar is 40 µm.
