## Supplementary material for "Directed Differentiation of Human iPSCs to Functional Ovarian Granulosa-Like Cells via Transcription Factor Overexpression": Source Data: barcode_plots.ipynb

```
# %% [code]
import pandas as pd

counts_data = pd.read_csv("2021-12-09_reanalysis_for_figure.csv", index_col = "Unnamed: 0")
counts_data

# %% [code]
#Combine reads from different barcodes of the same TF
counts_data.drop(["Barcode"], axis = 1, inplace = True)
counts_data = counts_data[counts_data["Plasmid"]!= "Unmatched"]
counts_data = counts_data.groupby('Plasmid').sum()

# %% [code]
counts_data_normalized = counts_data

for col in counts_data:
    if col == "Barcode" or col == "Plasmid": continue
    
    
    total = sum(counts_data_normalized[col][0:len(counts_data_normalized[col])])
    
    counts_data_normalized[col] = counts_data_normalized[col] / total
    
counts_data_normalized

# %% [code]
#Remove very small read frequency values
include_thresh = 0.0001
cdn = counts_data_normalized.clip(lower = include_thresh)
cdn.replace({include_thresh:float("NaN")}, inplace = True)
cdn

# %% [code]
#initialize the fold-change summary

def add_FC_col(df, output_df, col_denominator, col_numerator, new_colname, thresh = 0.0005):
    #initialize to NaNs
    new_column = df[col_denominator]*float("NaN")
    new_column[df[col_denominator] > thresh] = df[df[col_denominator] > thresh][col_numerator] / df[df[col_denominator] > thresh][col_denominator]
    
    df_new = output_df.copy()
    df_new[new_colname] = new_column
    return df_new

fc_summary = pd.DataFrame(index = cdn.index)

#mTESR minus vs day 0
fc_summary = add_FC_col(cdn, fc_summary, "S01", "S08", "B50_mTESR_minus_vs_day0", thresh = 0.01)
fc_summary = add_FC_col(cdn, fc_summary, "S02", "S09", "N50_mTESR_minus_vs_day0", thresh = 0.01)
fc_summary = add_FC_col(cdn, fc_summary, "S03", "S10", "U50_mTESR_minus_vs_day0", thresh = 0.01)
fc_summary = add_FC_col(cdn, fc_summary, "S04", "S11", "B5_mTESR_minus_vs_day0", thresh = 0.01)
fc_summary = add_FC_col(cdn, fc_summary, "S05", "S12", "N5_mTESR_minus_vs_day0", thresh = 0.01)
fc_summary = add_FC_col(cdn, fc_summary, "S06", "S13", "U5_mTESR_minus_vs_day0", thresh = 0.01)
fc_summary = add_FC_col(cdn, fc_summary, "S07", "S14", "C5_mTESR_minus_vs_day0", thresh = 0.01)

#Meso minus vs day 0
fc_summary = add_FC_col(cdn, fc_summary, "S01", "S15", "B50_meso_minus_vs_day0", thresh = 0.01)
fc_summary = add_FC_col(cdn, fc_summary, "S02", "S16", "N50_meso_minus_vs_day0", thresh = 0.01)
fc_summary = add_FC_col(cdn, fc_summary, "S03", "S17", "U50_meso_minus_vs_day0", thresh = 0.01)
fc_summary = add_FC_col(cdn, fc_summary, "S04", "S18", "B5_meso_minus_vs_day0", thresh = 0.01)
fc_summary = add_FC_col(cdn, fc_summary, "S05", "S19", "N5_meso_minus_vs_day0", thresh = 0.01)
fc_summary = add_FC_col(cdn, fc_summary, "S06", "S20", "U5_meso_minus_vs_day0", thresh = 0.01)
fc_summary = add_FC_col(cdn, fc_summary, "S07", "S21", "C5_meso_minus_vs_day0", thresh = 0.01)
fc_summary.to_csv("negative_vs_day0.csv")
fc_summary

# %% [code]
fc_summary_pos_day0= pd.DataFrame(index = cdn.index)

#mTESR plus vs day 0
fc_summary_pos_day0 = add_FC_col(cdn, fc_summary_pos_day0, "S01", "S22", "B50_mTESR", thresh = 0.01)
#fc_summary_pos_day0 = add_FC_col(cdn, fc_summary_pos_day0, "S02", "S23", "N50_mTESR", thresh = 0.01)
#fc_summary_pos_day0 = add_FC_col(cdn, fc_summary_pos_day0, "S03", "S24", "U50_mTESR", thresh = 0.01)
fc_summary_pos_day0 = add_FC_col(cdn, fc_summary_pos_day0, "S04", "S25", "B5_mTESR", thresh = 0.01)
#fc_summary_pos_day0 = add_FC_col(cdn, fc_summary_pos_day0, "S05", "S26", "N5_mTESR", thresh = 0.01)
#fc_summary_pos_day0 = add_FC_col(cdn, fc_summary_pos_day0, "S06", "S27", "U5_mTESR", thresh = 0.01)
fc_summary_pos_day0 = add_FC_col(cdn, fc_summary_pos_day0, "S07", "S28", "C5_mTESR", thresh = 0.01)

#Meso plus vs day 0
fc_summary_pos_day0 = add_FC_col(cdn, fc_summary_pos_day0, "S01", "S29", "B50_meso", thresh = 0.01)
#fc_summary_pos_day0 = add_FC_col(cdn, fc_summary_pos_day0, "S02", "S30", "N50_meso", thresh = 0.01)
#fc_summary_pos_day0 = add_FC_col(cdn, fc_summary_pos_day0, "S03", "S31", "U50_meso", thresh = 0.01)
fc_summary_pos_day0 = add_FC_col(cdn, fc_summary_pos_day0, "S04", "S32", "B5_meso", thresh = 0.01)
#fc_summary_pos_day0 = add_FC_col(cdn, fc_summary_pos_day0, "S05", "S33", "N5_meso", thresh = 0.01)
#fc_summary_pos_day0 = add_FC_col(cdn, fc_summary_pos_day0, "S06", "S34", "U5_meso", thresh = 0.01)
fc_summary_pos_day0 = add_FC_col(cdn, fc_summary_pos_day0, "S07", "S35", "C5_meso", thresh = 0.01)
fc_summary_pos_day0.to_csv("reanalysis_positive_vs_day0.csv")
fc_summary_pos_day0

# %% [code]
fc_summary_pos_neg= pd.DataFrame(index = cdn.index)

#mTESR plus vs neg
fc_summary_pos_neg = add_FC_col(cdn, fc_summary_pos_neg, "S08", "S22", "B50_mTESR_plus_vs_neg", thresh = 0.005)
fc_summary_pos_neg = add_FC_col(cdn, fc_summary_pos_neg, "S09", "S23", "N50_mTESR_plus_vs_neg", thresh = 0.005)
fc_summary_pos_neg = add_FC_col(cdn, fc_summary_pos_neg, "S10", "S24", "U50_mTESR_plus_vs_neg", thresh = 0.005)
fc_summary_pos_neg = add_FC_col(cdn, fc_summary_pos_neg, "S11", "S25", "B5_mTESR_plus_vs_neg", thresh = 0.005)
fc_summary_pos_neg = add_FC_col(cdn, fc_summary_pos_neg, "S12", "S26", "N5_mTESR_plus_vs_neg", thresh = 0.005)
fc_summary_pos_neg = add_FC_col(cdn, fc_summary_pos_neg, "S13", "S27", "U5_mTESR_plus_vs_neg", thresh = 0.005)
fc_summary_pos_neg = add_FC_col(cdn, fc_summary_pos_neg, "S14", "S28", "C5_mTESR_plus_vs_neg", thresh = 0.005)

#Meso plus vs neg
fc_summary_pos_neg = add_FC_col(cdn, fc_summary_pos_neg, "S15", "S29", "B50_meso_plus_vs_neg", thresh = 0.005)
fc_summary_pos_neg = add_FC_col(cdn, fc_summary_pos_neg, "S16", "S30", "N50_meso_plus_vs_neg", thresh = 0.005)
fc_summary_pos_neg = add_FC_col(cdn, fc_summary_pos_neg, "S17", "S31", "U50_meso_plus_vs_neg", thresh = 0.005)
fc_summary_pos_neg = add_FC_col(cdn, fc_summary_pos_neg, "S18", "S32", "B5_meso_plus_vs_neg", thresh = 0.005)
fc_summary_pos_neg = add_FC_col(cdn, fc_summary_pos_neg, "S19", "S33", "N5_meso_plus_vs_neg", thresh = 0.005)
fc_summary_pos_neg = add_FC_col(cdn, fc_summary_pos_neg, "S20", "S34", "U5_meso_plus_vs_neg", thresh = 0.005)
fc_summary_pos_neg = add_FC_col(cdn, fc_summary_pos_neg, "S21", "S35", "C5_meso_plus_vs_neg", thresh = 0.005)
fc_summary_pos_neg.to_csv("reanalysis_positive_vs_neg.csv")
fc_summary_pos_neg

# %% [code]
import matplotlib.pyplot as plt
import seaborn as sns
import numpy as np

#log_fc=log_fc[log_fc["Plasmid"] != CK544_GFP]

# %% [code]
log_fc = np.log2(fc_summary)
log_fc_pos_day0 = np.log2(fc_summary_pos_day0)
log_fc_pos_neg = np.log2(fc_summary_pos_neg)

# %% [code]
log_fc_pos_day0["Sum"] = log_fc_pos_day0.sum(axis = 1)
log_fc_pos_day0.sort_values(ascending=False, by = "Sum", inplace = True)
log_fc_pos_day0.drop(columns = ["Sum"], inplace = True)

# %% [code]
f, ax = plt.subplots(figsize=(5, 10))
sns.heatmap(log_fc_pos_day0, xticklabels=True, yticklabels=True, annot=True, cmap='coolwarm',fmt=".1f", vmin = -2, vmax = 2)
plt.title("TF Log2 Fold Enrichment (FOXL2+ vs. Day 0)", fontsize =14)
plt.xlabel("Experimental Replicate")
plt.ylabel("TF Barcode")
plt.savefig("reanalysis_2021-12-09_heatmap2.png",bbox_inches='tight', dpi = 128)

# %% [code]
f, ax = plt.subplots(figsize=(6, 10))
sns.heatmap(log_fc_pos_neg, xticklabels=True, yticklabels=True, annot=True, cmap='coolwarm',fmt=".1f", vmin = -2, vmax = 2)
plt.title("Plasmid Log2 Fold Enrichment")
plt.xlabel("Experimental Replicate")
#plt.savefig("2021-06-01_heatmap3.png",bbox_inches='tight', dpi = 128)

# %% [code]
```
