## Supplementary material for "Directed Differentiation of Human iPSCs to Functional Ovarian Granulosa-Like Cells via Transcription Factor Overexpression": Source Data: 2022-06-07_replot_ELISA.ipynb

```
# %% [code]
#Source: https://people.duke.edu/~ccc14/pcfb/analysis.html

import numpy as np
import numpy.random as npr
import matplotlib.pyplot as plt
from scipy.optimize import leastsq

def logistic4(x, A, B, C, D):
    """4PL logistic equation."""
    return ((A-D)/(1.0+((x/C)**B))) + D

def residuals(p, y, x):
    """Deviations of data from fitted 4PL curve"""
    A,B,C,D = p
    err = y-logistic4(x, A, B, C, D)
    return err

def peval(x, p):
    """Evaluated value at x with current parameters."""
    A,B,C,D = p
    return logistic4(x, A, B, C, D)

# Make up some data for fitting and add noise
# In practice, y_meas would be read in from a file
x = np.linspace(0,20,20)
A,B,C,D = 0.5,2.5,8,7.3
y_true = logistic4(x, A, B, C, D)
y_meas = y_true + 0.2*npr.randn(len(x))

# Initial guess for parameters
p0 = [0, 1, 1, 1]

# Fit equation using least squares optimization
plsq = leastsq(residuals, p0, args=(y_meas, x))

# Plot results
plt.plot(x,peval(x,plsq[0]),x,y_meas,'o',x,y_true)
plt.title('Least-squares 4PL fit to noisy data')
plt.legend(['Fit', 'Noisy', 'True'], loc='upper left')
for i, (param, actual, est) in enumerate(zip('ABCD', [A,B,C,D], plsq[0])):
    plt.text(10, 3-i*0.5, '%s = %.2f, est(%s) = %.2f' % (param, actual, param, est))
#plt.savefig('logistic.png')

# %% [code]
import pandas as pd
std = pd.read_csv("standards.csv")
std

# %% [code]
y1 = std['A450_corrected']
x1 = std['pg_mL']

fit = leastsq(residuals, p0, args=(y1, x1))


test = np.linspace(0,2000, 50)
test

plt.plot(test,peval(test,fit[0]),x1,y1,'o')
plt.savefig('2021-10-07_standards_logistic_4param.png')

# %% [code]
#Interpolate using the inverse function
#x=p(3).*(((p(1)-p(4))./(y-p(4)))-1).^(1/p(2));
def inv_logistic4(p, y): #p = model parameters
    x = p[2] * (((p[0]-p[3])/(y-p[3]))-1)**(1/p[1])
    return x

test2 = np.linspace(0.3, 2, 10)
plt.plot(test,peval(test,fit[0]),inv_logistic4(fit[0], test2),test2,'o')

# %% [code]
data = pd.read_csv("2021-10-07_ELISA_data.csv")
data["pg_uL_raw"] = inv_logistic4(fit[0], data["A450_adj"])
standards_vol = 25
data["pg_uL"] = data["pg_uL_raw"]/data["Vol"] * standards_vol
data

# %% [code]
data.to_csv("2021-10-07_ELISA_results.csv")

# %% [code]
import seaborn as sns
import matplotlib.pyplot as plt

# %% [code]
plt.rcParams['font.size'] = 18#12#'24'
sns.catplot(data = data, x = "Cells", y = "pg_uL", hue = "Condition", kind = 'bar', hue_order=["Minus", "FK", "FSH", "FSH+FK"])
plt.ylabel("Estradiol (pg/mL)")
plt.title("Estradiol production: response to stimulus")
plt.xticks(rotation = 90)
plt.savefig("2021-10-07_ELISA_results_replot2.svg")
plt.savefig("2021-10-07_ELISA_results_replot2.png", dpi = 200, bbox_inches = "tight")

# %% [code]
plt.rcParams['figure.figsize'] = [8, 4]
plt.rcParams['font.size'] = 12#'24'

data2 = data[data["Cells"] != "4F"]
sns.catplot(data = data2, x = "Cells", y = "pg_uL", hue = "Condition", kind = 'bar', hue_order=["Minus", "FK", "FSH", "FSH+FK"])
plt.ylabel("Estradiol (pg/mL)", fontsize = 14)
plt.title("Estradiol production: response to stimulus", fontsize = 14)
plt.savefig("2021-10-07_ELISA_results_no_4F_rereplot.svg")
plt.savefig("2021-10-07_ELISA_results_no_4F_rereplot.png", dpi = 200, bbox_inches = "tight")

# %% [code]
```
