## Supplementary material for "Directed Differentiation of Human iPSCs to Functional Ovarian Granulosa-Like Cells via Transcription Factor Overexpression": Source Data: 2022-03-08_ELISA_analysis.ipynb

```
# %% [code]
#Source: https://people.duke.edu/~ccc14/pcfb/analysis.html

import numpy as np
import numpy.random as npr
import matplotlib.pyplot as plt
from scipy.optimize import leastsq

def logistic4(x, A, B, C, D):
    """4PL logistic equation."""
    return ((A-D)/(1.0+((x/C)**B))) + D

def residuals(p, y, x):
    """Deviations of data from fitted 4PL curve"""
    A,B,C,D = p
    err = y-logistic4(x, A, B, C, D)
    return err

def peval(x, p):
    """Evaluated value at x with current parameters."""
    A,B,C,D = p
    return logistic4(x, A, B, C, D)

# Make up some data for fitting and add noise
# In practice, y_meas would be read in from a file
x = np.linspace(0,20,20)
A,B,C,D = 0.5,2.5,8,7.3
y_true = logistic4(x, A, B, C, D)
y_meas = y_true + 0.2*npr.randn(len(x))

# Initial guess for parameters
p0 = [0, 1, 1, 1]

# Fit equation using least squares optimization
plsq = leastsq(residuals, p0, args=(y_meas, x))

# Plot results
plt.plot(x,peval(x,plsq[0]),x,y_meas,'o',x,y_true)
plt.title('Least-squares 4PL fit to noisy data')
plt.legend(['Fit', 'Noisy', 'True'], loc='upper left')
for i, (param, actual, est) in enumerate(zip('ABCD', [A,B,C,D], plsq[0])):
    plt.text(10, 3-i*0.5, '%s = %.2f, est(%s) = %.2f' % (param, actual, param, est))
#plt.savefig('logistic.png')

# %% [code]
import pandas as pd
std_E = pd.read_csv("standards_estradiol.csv")
std_P = pd.read_csv("standards_progesterone.csv")
std_P

# %% [code]
y1_E = std_E['A450_corrected']
x1_E = std_E['pg_mL']

guess = [2, 1, 500, 0]

fit_E = leastsq(residuals, guess, args=(y1_E, x1_E))


test = np.linspace(0,2000, 50)
test

plt.plot(test,peval(test,fit_E[0]),x1_E,y1_E,'o')
#plt.savefig('2021-10-07_standards_logistic_4param.png')

# %% [code]
fit_E

# %% [code]
y1_P = std_P['A450_corrected']
x1_P = std_P['pg_mL']

guess = [2, 1, 500, 0]

fit_P = leastsq(residuals, guess, args=(y1_P, x1_P))


test = np.linspace(0,40000, 50)
test

plt.plot(test,peval(test,fit_P[0]),x1_P,y1_P,'o')
#plt.savefig('2021-10-07_standards_logistic_4param.png')

# %% [code]
#Interpolate using the inverse function
#x=p(3).*(((p(1)-p(4))./(y-p(4)))-1).^(1/p(2));
def inv_logistic4(p, y): #p = model parameters
    x = p[2] * (((p[0]-p[3])/(y-p[3]))-1)**(1/p[1])
    return x

test2 = np.linspace(0.3, 2, 10)
plt.plot(test,peval(test,fit_E[0]),inv_logistic4(fit_E[0], test2),test2,'o')

# %% [code]
data = pd.read_csv("data_all.csv")
data["pg_uL_estradiol"] = inv_logistic4(fit_E[0], data["A450_adj_estradiol"])
data["pg_uL_progesterone"] = inv_logistic4(fit_P[0], data["A450_adj_progesterone"])
data

# %% [code]
data.to_csv("2022-03-08_hormones.csv")

# %% [code]
```
