## Supplementary material for "Directed Differentiation of Human iPSCs to Functional Ovarian Granulosa-Like Cells via Transcription Factor Overexpression": Source Data: scRNAseq_D4_ovaroids_only_2022-06-07.ipynb

```
# %% [code]
import numpy as np
import pandas as pd
import scanpy as sc
import scipy
import os
import scipy.io as sio
import anndata
import matplotlib.pyplot as plt
import seaborn as sns

data_path = 'split_DGE/'

sc.settings.verbosity = 1 # verbosity: errors (0), warnings (1), info (2), hints (3)
sc.settings.set_figure_params(dpi=100, fontsize=10, dpi_save=300, figsize=(5,4), format='png')
sc.settings.figdir = 'scanpy-figures-D4-ovaroids-only_v2022-06-07'

# %% [code]
diff_key = pd.read_csv("ovaroid_diff_key.csv")
diff_key

# %% [code]
# reading in gene and cell data
gene_data = pd.read_csv('all_genes.csv')

sample_list = ['4','5','7','8','10','11','13','14']#,]

cell_meta = pd.concat([pd.read_csv(data_path + '1mil-'+ i +'_cell_metadata.csv') for i in sample_list])
cell_meta

# %% [code]
# The DGE_filtered folder contains the expression matrix, genes, and files 
adata = anndata.concat([sc.read_mtx(data_path + '1mil-'+i+'.mtx.gz') for i in sample_list], axis = 0)
#adata.raw = adata
adata.shape

# %% [code]
# add cell meta data to anndata object
adata.obs = cell_meta
adata.obs.set_index('bc_wells', inplace=True)
adata.obs.index.name = None
adata.obs_names_make_unique()

# %% [code]
#add sample info
adata.obs = adata.obs.merge(diff_key, on = 'sample', how = 'left')
adata.obs.index = adata.obs.index.map(str)
adata.obs

# %% [code]
# find genes with nan values and filter
gene_data = gene_data[gene_data.gene_name.notnull()]
notNa = gene_data.index
notNa = notNa.to_list()

# remove genes with nan values and assign gene names
adata = adata[:,notNa]
adata.var = gene_data
adata.var.set_index('gene_name', inplace=True)
adata.var.index.name = None
adata.var_names_make_unique()

# %% [code]
sc.pp.filter_cells(adata, min_genes=1000)
sc.pp.filter_genes(adata, min_cells=5)

# Returns the dimensions of the expression matrix (cells, genes)
adata.shape

# %% [code]
#Check for too many mitochondrial genes
adata.var['mt'] = adata.var_names.str.startswith('MT-')
sc.pp.calculate_qc_metrics(adata, qc_vars=['mt'], percent_top=None, log1p=False, inplace=True)

# Scanpy will prepend the string in the save argument with "violin"
# and save it to our figure directory defined in the first step.
sc.pl.violin(adata, ['n_genes_by_counts'], save='_n_genes', jitter=0.4)
sc.pl.violin(adata, ['total_counts'], save='_total_counts', jitter=0.4)
#plt.ylim([0,100000])
sc.pl.violin(adata, ['pct_counts_mt'], save='_mito_pct', jitter=0.4)

# %% [code]
# Filter the data
adata = adata[adata.obs.n_genes_by_counts < 6250,:]
adata = adata[adata.obs.total_counts < 40000,:]
adata = adata[adata.obs.pct_counts_mt < 10,:]
adata.shape # Checking number of cells remaining

# %% [code]
#Visualize QC
sc.pl.scatter(adata, x='total_counts', y='n_genes_by_counts', save='_gene_vs_transcript_counts')
print('median transcript count per cell: ' + str(adata.obs['tscp_count'].median(0)))
print('median gene count per cell: ' + str(adata.obs['gene_count'].median(0)))

# %% [code]
#CPM normalize
sc.pp.normalize_total(adata, target_sum=1e6)
sc.pp.log1p(adata)

adata.raw = adata #Save raw CPMs for plotting

# %% [code]
#Identify highly variable genes and regress out transcript counts

#NOTE: DON'T remove non-variable genes. Just regress out the counts.

#sc.pp.highly_variable_genes(adata, min_mean=0.0125, max_mean=8, min_disp=0.25)
#sc.pl.highly_variable_genes(adata, save='') # scanpy generates the filename automatically
#adata = adata[:, adata.var.highly_variable]
sc.pp.regress_out(adata, ['total_counts'])
sc.pp.scale(adata, max_value=10)

# %% [code]
#PCA
sc.tl.pca(adata, svd_solver='arpack')
sc.pl.pca_variance_ratio(adata, log=True, n_pcs=50, save='') # scanpy generates the filename automatically

# %% [code]
#Clustering
#sc.settings.figdir = 'scanpy-figures-sorted1F'
sc.pp.neighbors(adata, n_neighbors=10, n_pcs=30)
sc.tl.umap(adata)
sc.tl.leiden(adata, resolution=0.05)
sc.pl.umap(adata, color=['leiden'], legend_fontsize=8, save='_leiden005_regress.svg')

# %% [code]
sc.pl.umap(adata, color=['sample'], legend_fontsize=8, save='_sample_regress')

# %% [code]
sc.pl.umap(adata, color=['Day'], legend_fontsize=8, save='_Day_regress')

# %% [code]
#Rank markers by cluster
sc.tl.rank_genes_groups(adata, 'leiden', method='t-test')

# The head function returns the top n genes per cluster
top_markers = pd.DataFrame(adata.uns['rank_genes_groups']['names']).head(15)
print(top_markers)
top_markers.to_csv(os.path.join(sc.settings.figdir,"2022-06-07_D4_ovaroids_top_markers_regress.csv"))

# %% [code]
#Save the gene ranking for each cluster
names_df = pd.DataFrame(adata.uns['rank_genes_groups']['names'])

scores_df = pd.DataFrame(adata.uns['rank_genes_groups']['scores'])

pvals_adj_df = pd.DataFrame(adata.uns['rank_genes_groups']['pvals_adj'])

fc_df = pd.DataFrame(adata.uns['rank_genes_groups']['logfoldchanges'])

for i in range(0,3):
    j = str(i)
    cluster_df = pd.concat((names_df[j],scores_df[j],fc_df[j],pvals_adj_df[j]),axis = 1, keys = ('Gene name', 'score','logfc','p_adj'))
    cluster_df.to_csv(os.path.join(sc.settings.figdir,"2022-06-07_D4_ovaroids_cluster_"+j+".csv"))

# %% [code]
#Rank markers by sample
#sc.tl.rank_genes_groups(adata, 'sample', method='t-test')

# The head function returns the top n genes per sample
#top_markers = pd.DataFrame(adata.uns['rank_genes_groups']['names']).head(15)
#print(top_markers)

# %% [code]
#gene_list = ['FOXL2', 'NR5A1', 'AMHR2', 'WT1', 'GATA4', 'RUNX1', 'RUNX2', 'EGFR', 'STS', 'FSHR', 'STAR', 'AMH',
#             'KITLG', 'CYP19A1', 'CYP17A1', 'DHH', 'INHA', 'INHBA','WNT4', 'RSPO1', 'CD82', 'CXCL12', 'IGF2',
#             'IGFBP7', 'NR2F2', 'TCF21', 'ALDH1A2', 'POU5F1', 'VIM', 'TBXT', 'TBX6', 'FOXF1','OSR1','KRT19',
#             'ZFPM2','LGR5','SIX1','SIX4','PBX1','CBX2','LHX9','EMX2','CLDN11','HSD3B2','NR0B1','CDKN1B','GADD45G',
#             'FST','BOLL','LHX8','SOHLH1','ZNF281','FIGLA','DAZL','DDX4','SOX17','CD38','REC8','NPM2','STRA8','SYCP3','GDF9','ZP3','XACT']

#gene_list += ['XIST','TFAP2C','ZBTB16','SALL4','KIT','SSEA4','THY1','EPCAM','GFRA1','UTF1','NGN3']

#gene_list = ['FOXL2','WNT4','PRDM1','PRDM14','NANOG','KIT']

#gene_list = ['HOXC9']

gene_list = ['IRX3','IRX5']

for i in gene_list:
    try:
        sc.pl.umap(adata, color=i, color_map='viridis', legend_fontsize=8, use_raw = True, save='_regress_'+i)
    except Exception as e:
        print(e)
        continue

# %% [code]
#gene_list = ['FOXL2', 'NR5A1', 'AMHR2', 'WT1', 'GATA4', 'RUNX1', 'RUNX2', 'EGFR', 'STS', 'FSHR', 'STAR', 'AMH',
#             'KITLG', 'CYP19A1', 'CYP17A1', 'DHH', 'INHA', 'INHBA','WNT4', 'RSPO1', 'CD82', 'CXCL12', 'IGF2',
#             'IGFBP7', 'NR2F2', 'TCF21', 'ALDH1A2', 'POU5F1', 'VIM', 'TBXT', 'TBX6', 'FOXF1','OSR1','KRT19',
#             'ZFPM2','LGR5','SIX1','SIX4','PBX1','CBX2','LHX9','EMX2','CLDN11','HSD3B2','NR0B1','CDKN1B','GADD45G',
#             'FST','BOLL','LHX8','SOHLH1','ZNF281','FIGLA','DAZL','DDX4','SOX17','CD38','REC8','NPM2','STRA8','SYCP3','GDF9','ZP3',
#             'ZBTB16','SALL4','KIT','SSEA4','THY1','EPCAM','GFRA1','UTF1','NGN3','XACT']

gene_list2 = ['DAZL', 'DDX4']

for gene in gene_list2:
    try:
        adata.obs[gene + '+'] = (adata.raw[:,gene].X.todense() > 0)
    except Exception as e:
        print(e)
        print(gene + ' not found')
#adata.obs['DAZL_plus'] = adata.obs['DAZL_plus']#.astype("bool")
adata.obs.to_csv(os.path.join(sc.settings.figdir,"D4_obs.csv"))

# %% [code]
#Plot fraction of PGCLCs and DAZL+ cells by day
#PGCLCs are cluster 2 (double-check this if you change the clustering!)


day_df = adata.obs
day_df['PGCLC'] = day_df['leiden'] == '2'
grouped = day_df.groupby("Day").mean()#numeric_only = False)
grouped_plot = grouped['PGCLC']
grouped_plot = grouped[['DAZL+','DDX4+','PGCLC']]
#grouped_plot.rename({'DAZL_plus':'DAZL+','DDX4_plus':'DDX4+'})
grouped_plot.to_csv(os.path.join(sc.settings.figdir,"time_plot.csv"))

# %% [code]
grouped_plot

# %% [code]
melted = grouped_plot.reset_index().melt(id_vars='Day')
melted['percent'] = melted['value']*100
melted['Cells'] = melted['variable']
melted['Day'] = melted['Day'].astype(int) 

sns.set_theme(font_scale = 2, style = None)
sns.lineplot(data = melted, x = 'Day', y = 'percent', hue = 'Cells', marker = 'o')#, legend = False)
plt.ylim([0,3])
plt.legend(loc='upper right', prop={'size': 20})
plt.xticks([2,4,8,14])
plt.yticks([0,1,2,3])
plt.grid(axis = 'x')
plt.ylabel("% of cells in ovaroids")
plt.savefig(os.path.join(sc.settings.figdir,"ovaroid_time_plot.svg"))
plt.savefig(os.path.join(sc.settings.figdir,"ovaroid_time_plot.png"), dpi = 200, bbox_inches = "tight")

# %% [code]
#Rank markers by cluster
#adata.obs['DAZL_plus'] = adata.obs['DAZL_plus'].astype("category")
#sc.tl.rank_genes_groups(adata, 'DAZL_plus', method='t-test')

# The head function returns the top n genes per cluster
#top_markers = pd.DataFrame(adata.uns['rank_genes_groups']['names'])
#print(top_markers)
#top_markers.to_csv("ovaroid_DAZL_plus_top_markers_names.csv")

# %% [code]
#pd.DataFrame(adata.X.to_dense()).to_csv("D4_ovaroid_dump.csv")

# %% [code]
markers = ['XACT','XIST','TSIX']
sc.pl.stacked_violin(adata, markers, groupby='leiden', figsize = (6,6), save = 'ChrX_lncRNA_cluster_violin2.svg', row_palette=sns.color_palette("tab10"))#, stripplot=True, jitter = True)#, use_raw = True)#, dendrogram=True)
#plt.savefig("")

# %% [code]
markers = ['FOXL2','WNT4','CD82','RUNX1','NR5A1','CDKN1B',"CD38","KIT", "PRDM1", "TFAP2C", "PRDM14", "NANOG",'POU5F1','XACT','XIST','TSIX']
sc.pl.stacked_violin(adata, markers, groupby='leiden', figsize = (12,6), save = 'genes_cluster_violin_long.svg', row_palette=sns.color_palette("tab10"))#, stripplot=True, jitter = True)#, use_raw = True)#, dendrogram=True)


# %% [code]
```
