## Supplementary material for "Directed Differentiation of Human iPSCs to Functional Ovarian Granulosa-Like Cells via Transcription Factor Overexpression": Source Data: plotdata_extended.Rmd

````
---
title: "qpCR_fit_plots"
output: html_document
---

```{r setup, include=FALSE}
#knitr::opts_chunk$set(echo = TRUE)
rm(list=ls())
```

```{r}
library(dplyr)
library(ggplot2)
library(svglite)
```

```{r}
data = read.csv("/Users/merrickpiersonsmela/Documents/Church_Lab/oogenesis/qPCR/Copynumber/2021-04-11_absolute_quant/qPCR_copy_data_extended.csv")

```

# Make a plot that is 1) plot data points 2) has a fitted curve 3) asymmetrical error bars

```{r}

k = 0.884
```

```{r}

ggplot(data) +
 aes(x = fmol, y = copies) +
 geom_point(size = 3L, colour = "#0c4c8a") +
  geom_errorbar(aes(ymin = data$errorLow, ymax = data$errorHigh), width=.3, position="dodge2") + 
  scale_x_continuous(expand = expansion(add = c(0,0)), limits = c(0,21)) +
  scale_y_continuous(expand = expansion(add = c(0,0)), limits = c(0,24)) +
  #scale_y_continuous(breaks = c(0,1,2,3,4,5,6,7), labels = c(0,1,2,3,4,5,6,7), limits = c(0,7.5))+
 labs(x = "fmol transposon plasmid", y = "copy number") +
 theme_minimal() + 
 stat_function(fun=function(x) (k * x / (1 - exp(-k * x))), colour = "#00000060") +
theme(text = element_text(size=20)) 
  
ggsave("qPCR_extended.svg")
ggsave("qPCR_extended.png")
```
````
