## Supplementary material for "Directed Differentiation of Human iPSCs to Functional Ovarian Granulosa-Like Cells via Transcription Factor Overexpression": Source Data: 2022-06-12-replot.ipynb

```
# %% [code]
from cytoflow import *

import pandas as pd
import numpy as np
from scipy.stats import gaussian_kde

%matplotlib inline
from matplotlib import ticker
import matplotlib.pyplot as plt
import matplotlib.patches as patches
#Matplotlib settings
plt.rcParams['figure.figsize'] = [4, 3]
plt.rcParams['font.size'] = 24#'24'
plt.rcParams['figure.dpi'] = 120

#Function to read tube metadata from a .csv, then import into CytoFlow
#The .csv must have a column named 'Path' containing the .fcs file path
#Subsequent columns will be interpreted as metadata
def csv_tubes_import(csv_path, channels = 'infer', verbose = True, fcs_folder = ''):

    csv_df = pd.read_csv(csv_path)

    conditions = {'Tube': 'category'} #initialize conditions dictionary

    #Read the column metadata and extract conditions
    for condition in csv_df.dtypes.iteritems():
        #if condition[0] == 'Path': continue

        dtype = condition[1]

        if dtype == 'int64': dtype = 'float64' #Convert int to float
        elif dtype == 'bool': dtype = 'bool' #Keep bool
        elif dtype == 'object': dtype = 'category' #Convert object to category

        conditions[condition[0]] = dtype
    
    if verbose: 
        print('Importing tubes from: ' + csv_path)
        print('using conditions: ')
        print(conditions)

    #Read the tubes
    tubes = []
    for index, tube in csv_df.iterrows():
        tubeConditions = {'Tube': str(index)}

        for condition in csv_df:
            #if condition == 'Path': continue
            tubeConditions[condition] = tube[condition]

        tubes.append(Tube(file = fcs_folder + tube['Path'], conditions = tubeConditions))
        if verbose: print('Read file: ' + fcs_folder + tube['Path'])
    
    if channels == 'infer': #infer channels. Must match!
        op_auto = ImportOp(conditions = conditions, tubes = tubes)
    else:
        op_auto = ImportOp(conditions = conditions, tubes = tubes, channels = channels)
    return op_auto.apply()

def flowPlotKDE(ex, channel_x = None, channel_y = None, subset = None, kde_N = 10000, xlabel = None, ylabel = None, title = None, logscale = False, cmap = 'plasma', cbar = False, pointsize = 2):
    
    if subset is not None:
        df = ex.data.query(subset)
    else:
        df = ex.data

    if logscale:
        #Drop negative or zero points
        df_pos = df[(df[channel_x] > 0) & (df[channel_y] > 0)]
        
        x = np.log10(df_pos[channel_x])
        y = np.log10(df_pos[channel_y])
        if xlabel is None: xlabel = 'log10(' + channel_x + ')'
        if ylabel is None: ylabel = 'log10(' + channel_y + ')' 
    else:
        x = df[channel_x]
        y = df[channel_y]
        if xlabel is None: xlabel = channel_x
        if ylabel is None: ylabel = channel_y

    # Calculate the point density (using a sample size of kde_N)
    xy = np.vstack([x,y])
    
    xy_subsample = xy[:, np.random.choice(xy.shape[1], size = kde_N, replace = False)]
    
    z = gaussian_kde(xy_subsample)(xy)
    
    #Make the plot
    plot = plt.scatter(x, y, c=z, s=pointsize, edgecolor='', cmap = cmap)
    
    #Make the colorbar
    if cbar: plt.colorbar(plot)
    
    #Add axis labels and title
    plt.xlabel(xlabel)
    plt.ylabel(ylabel)
    plt.title(title)
    
    #plt.show()
    
    return plot

def flowPlotHist(ex, channel_x = None, channel_y = None, subset = None, bins = [100, 100], xlabel = None, ylabel = None, title = None, logscale = False, cmap = 'plasma', cbar = False, pointsize = 2):
    
    if subset is not None:
        df = ex.data.query(subset)
    else:
        df = ex.data

    if logscale:
        #Drop negative or zero points
        df_pos = df[(df[channel_x] > 0) & (df[channel_y] > 0)]
        
        x = np.log10(df_pos[channel_x])
        y = np.log10(df_pos[channel_y])
        if xlabel is None: xlabel = 'log10(' + channel_x + ')'
        if ylabel is None: ylabel = 'log10(' + channel_y + ')' 
    else:
        x = df[channel_x]
        y = df[channel_y]
        if xlabel is None: xlabel = channel_x
        if ylabel is None: ylabel = channel_y

    # Calculate the point density
    hist, locx, locy = np.histogram2d(x, y, bins = bins)
    z = np.array([hist[np.argmax(a<=locx[1:]),np.argmax(b<=locy[1:])] for a,b in zip(x,y)])

    #Make the plot
    plot = plt.scatter(x, y, c=z, s=pointsize, edgecolor='', cmap = cmap)
    
    #Make the colorbar
    if cbar:
        bar = plt.colorbar(plot)
        for t in bar.ax.get_yticklabels():
             t.set_fontsize(14)
        bar.set_label('Counts')
        bar.ax.tick_params(labelsize=14)
        
    #Add axis labels and title
    plt.xlabel(xlabel)
    plt.ylabel(ylabel)
    plt.title(title)
    
    #plt.show()
    
    return plot

def flowPlotHist_gate(ex, channel_x = None, channel_y = None, subset = None, poly_gate = None, ellipse_gate = None, bins = [100, 100], xlabel = None, ylabel = None, title = None, logscale = False, cmap = 'plasma', cbar = False, pointsize = 2, cbar_fontsize = 14, xlim = None, ylim = None):
    
    if subset is not None:
        df = ex.data.query(subset)
    else:
        df = ex.data

    if logscale:
        #Drop negative or zero points
        df_pos = df[(df[channel_x] > 0) & (df[channel_y] > 0)]
        
        x = np.log10(df_pos[channel_x])
        y = np.log10(df_pos[channel_y])

    else:
        x = df[channel_x]
        y = df[channel_y]
        
    if xlabel is None: xlabel = channel_x
    if ylabel is None: ylabel = channel_y

    # Calculate the point density
    hist, locx, locy = np.histogram2d(x, y, bins = bins)
    z = np.array([hist[np.argmax(a<=locx[1:]),np.argmax(b<=locy[1:])] for a,b in zip(x,y)])

    #Make the plot
    plot = plt.scatter(x, y, c=z, s=pointsize, edgecolor='', cmap = cmap)
    
    #Make the colorbar
    if cbar:
        bar = plt.colorbar(plot)
        for t in bar.ax.get_yticklabels():
             t.set_fontsize(cbar_fontsize)
        bar.set_label('Counts', fontsize = cbar_fontsize)
    
    #Draw the gates
    ax = plt.gca()
    if poly_gate is not None:
        ax.add_patch(patches.Polygon(poly_gate, linewidth=1, fill = False, edgecolor = 'black'))
        
    if ellipse_gate is not None:
        ax.add_patch(patches.Ellipse(*ellipse_gate, linewidth=1, fill = False, edgecolor = 'red'))
        
    #Add axis labels and title
    plt.xlabel(xlabel)
    plt.ylabel(ylabel)
    plt.title(title)
    
    #if log scale, make the ticks
    if logscale:
        if ylim is not None:
            yticks_lim = [np.floor(ylim[0]).astype('int'), np.ceil(ylim[1]).astype('int')]
            
            ax.yaxis.set_major_formatter(ticker.StrMethodFormatter("$10^{{{x:.0f}}}$"))
            
            yticks_minor = [np.log10(x) for p in range(yticks_lim[0],yticks_lim[1]) for x in np.linspace(2 * 10**p, 9 * 10**p, 8)]
            ax.set_yticks(yticks_minor, minor=True)
            
            yticks_major = range(yticks_lim[0],yticks_lim[1]+1)
            ax.set_yticks(yticks_major)
            
        if xlim is not None:
            xticks_lim = [np.floor(xlim[0]).astype('int'), np.ceil(xlim[1]).astype('int')]
            
            ax.xaxis.set_major_formatter(ticker.StrMethodFormatter("$10^{{{x:.0f}}}$"))
            
            xticks_minor = [np.log10(x) for p in range(xticks_lim[0],xticks_lim[1]) for x in np.linspace(2 * 10**p, 9 * 10**p, 8)]
            ax.set_xticks(xticks_minor, minor=True)
    
            xticks_major = range(xticks_lim[0],xticks_lim[1]+1)
            ax.set_xticks(xticks_major)
    
    #Add axis limits
    plt.xlim(xlim)
    plt.ylim(ylim)
    
    return plot

def ellipse_to_poly(center = [0,0], len_x = 1, len_y = 1, rotation = 0, n_points = 24):
    
    angles = np.linspace(start = 0, stop = 360, num = n_points, endpoint = False)
    
    x = np.cos(np.deg2rad(angles)) * len_x / 2
    y = np.sin(np.deg2rad(angles)) * len_y / 2
    
    #perform the rotation
    c, s = np.cos(np.deg2rad(rotation)), np.sin(np.deg2rad(rotation))
    
    xy = np.array(((c, -s), (s, c))) @ np.array((x, y))

    return np.transpose(xy) + center

# %% [code]
fcs_folder = 'Cells/'
ex_0 = csv_tubes_import('flow_files_day5.csv', fcs_folder = fcs_folder)

# %% [code]
ssc_gate = [(65000,15000),(75000,200000),(120000,200000),(90000,15000)]
#see: https://cancer.wisc.edu/research/wp-content/uploads/2017/08/Flow_TechNotes_Doublet-Discrimination_20170918.pdf
flowPlotHist_gate(ex_0, channel_x = 'SSC-W', channel_y = 'SSC-A', poly_gate = ssc_gate, ylim = (0, 200000),
                  subset = "(Cells == '1F')")

# %% [code]
SingletOp = PolygonOp(
    name='singlets',
    xchannel='SSC-W',
    ychannel='SSC-A',
    vertices=ssc_gate)

#SingletOp.estimate(ex_auto2)
ex_1 = SingletOp.apply(ex_0)
#SingletOp.default_view().plot(ex_1)#, xlim=(1000, 300000.0), ylim=(None, 300000.0))

# %% [code]
cells_gate = [(10**4.25, 10**4.4), 
              (10**5.5, 10**4.5),
              (10**5.5, 10**5.2), 
              (10**4.3, 10**5)]

flowPlotHist_gate(ex_1, channel_x = 'FSC-A', channel_y = 'SSC-A', poly_gate = np.log10(cells_gate), xlim = (3.5, 5.5), ylim = (2.5, 5.5), logscale = True,
                  subset = "singlets and (Cells == '13D7')")

# %% [code]
flowPlotHist_gate(ex_1, channel_x = 'FSC-A', channel_y = 'SSC-A', poly_gate = np.log10(cells_gate), xlim = (3.5, 5.5), ylim = (2.5, 5.5), logscale = True,
                  subset = "singlets and (Cells == '1F')")

# %% [code]
CellsOp = PolygonOp(
    name='single_cells',
    xchannel='FSC-A',
    ychannel='SSC-A',
    vertices=cells_gate)

#SingletOp.estimate(ex_auto2)
ex_2 = CellsOp.apply(ex_1)

# %% [code]
flowPlotHist_gate(ex_2, channel_x = 'FSC-H', channel_y = 'FSC-W', subset = "Cells == '1F' and single_cells and singlets", xlim = (0, 300000))

# %% [code]
#Live plot
live_gate = [(10**4.4, 100),
              (10**5.2, 1000),
              (10**5.2, 10**4.5),
              (10**4.4, 10**3.75)]

# %% [code]
flowPlotHist_gate(ex_2, channel_x = 'SSC-A', channel_y = 'Indo-1 (Blue)-A', logscale = True, poly_gate = np.log10(live_gate),
                  subset = '(Cells == "1F") and single_cells and singlets')

# %% [code]
flowPlotHist_gate(ex_2, channel_x = 'SSC-A', channel_y = 'Indo-1 (Blue)-A', logscale = True, poly_gate = np.log10(live_gate),
                  subset = '(Cells == "13D7_iPSC") and single_cells and singlets')

# %% [code]
LiveOp = PolygonOp(
    name='Live',
    xchannel='SSC-A',
    ychannel='Indo-1 (Blue)-A',
    vertices=live_gate,
    xscale='log',
    yscale='log')

ex_3 = LiveOp.apply(ex_2)
#LiveOp.default_view(subset='(singlets == True)').plot(ex_2)

# %% [code]
BgOp = AutofluorescenceOp(
    channels=[
        'PE-Texas Red-A','APC-A', 'PE-Cy5-5-A', 'APC-Cy7-A'
    ],
    blank_file = "Compensation Panel/controls_F66_unstained_iPSCs_005.fcs"
)

BgOp.estimate(ex_3, subset='Live and singlets and single_cells')
ex_4 = BgOp.apply(ex_3)
BgOp.default_view().plot(ex_4)

# %% [code]
ex_3.metadata.values()

# %% [code]
CompOp = BleedthroughLinearOp(
    controls={
        'PE-Texas Red-A': 'Compensation Panel/controls_13D7_CRISPRa_unstained_iPSCs_004.fcs',
        'APC-A': 'Compensation Panel/controls_FSHR-APC_beads_003.fcs',
        'PE-Cy5-5-A': 'Compensation Panel/controls_CD82-PECy5-5_beads_001.fcs',
        'APC-Cy7-A': 'Compensation Panel/cells_F66_EPCAM_002.fcs'
    })

CompOp.estimate(ex_4)
ex_5 = CompOp.apply(ex_4)

# %% [code]
#tdT plot
tdT_gate = [(10**3.3, 10**4.3),
              (10**3.8, 10**5.4),
              (10**6, 10**5.4),
              (10**6, 10**4.3)]

flowPlotHist_gate(ex_5, channel_x = 'PE-Texas Red-A', channel_y = 'SSC-A', logscale = True, poly_gate = np.log10(tdT_gate),
                  subset = 'Live and single_cells and singlets and (Cells == "13D7")', xlim = (1,5))

# %% [code]
flowPlotHist_gate(ex_5, channel_x = 'PE-Texas Red-A', channel_y = 'SSC-A', logscale = True, poly_gate = np.log10(tdT_gate),
                  subset = 'Live and single_cells and singlets and (Cells == "1F")', xlim = (2,5))

# %% [code]
tdTOp = PolygonOp(
    name='tdT_plus',
    xchannel='PE-Texas Red-A',
    ychannel='SSC-A',
    vertices=tdT_gate,
    xscale='log',
    yscale='log')

#SingletOp.estimate(ex_auto2)
ex_6 = tdTOp.apply(ex_5)
#tdTOp.default_view(subset='Live and single_cells and singlets').plot(ex_4)

# %% [code]
#FSHR plot
FSHR_gate = [(10**3.4, 10**4.4),
              (10**3.4, 10**5.4),
              (10**6, 10**5.4),
              (10**6, 10**4.4)]

flowPlotHist_gate(ex_6, channel_x = 'APC-A', channel_y = 'SSC-A', logscale = False, poly_gate = np.log10(FSHR_gate),
                  subset = 'Live and single_cells and singlets and (Cells == "13D7")')

# %% [code]
#FSHR plot

flowPlotHist_gate(ex_6, channel_x = 'APC-A', channel_y = 'SSC-A', logscale = False, poly_gate = np.log10(FSHR_gate),
                  subset = 'Live and single_cells and singlets and (Cells == "1F")')

# %% [code]
#FSHR plot

flowPlotHist_gate(ex_6, channel_x = 'APC-A', channel_y = 'SSC-A', logscale = False, poly_gate = np.log10(FSHR_gate),
                  subset = 'Live and single_cells and singlets and (Cells == "COV434")')

# %% [code]
#FSHROp = PolygonOp(
#    name='FSHR_plus',
#    xchannel='APC-A',
#    ychannel='SSC-A',
#    vertices=FSHR_gate,
#    xscale='log',
#    yscale='log')

#SingletOp.estimate(ex_auto2)
#ex_7 = FSHROp.apply(ex_6)
#tdTOp.default_view(subset='Live and single_cells and singlets').plot(ex_4)
ex_7 = ex_6

# %% [code]
#CD82 plot
CD82_gate = [(10**2.6, 10**4.3),
              (10**2.6, 10**5.4),
              (10**6, 10**5.4),
              (10**6, 10**4.3)]

flowPlotHist_gate(ex_6, channel_x = 'PE-Cy5-5-A', channel_y = 'SSC-A', logscale = True, poly_gate = np.log10(CD82_gate),
                  subset = 'Live and single_cells and singlets and (Cells == "13D7_iPSC")', xlim = (1,4))

# %% [code]
CD82Op = PolygonOp(
    name='CD82_plus',
    xchannel='PE-Cy5-5-A',
    ychannel='SSC-A',
    vertices=CD82_gate,
    xscale='log',
    yscale='log')

#SingletOp.estimate(ex_auto2)
ex_8 = CD82Op.apply(ex_7)
#tdTOp.default_view(subset='Live and single_cells and singlets').plot(ex_4)

# %% [code]
EPCAM_gate = [(10**3.9, 10**4.4),
              (10**4.4, 10**5.2),
              (10**6, 10**5.2),
              (10**6, 10**4.4)]

flowPlotHist_gate(ex_6, channel_x = 'APC-Cy7-A', channel_y = 'SSC-A', logscale = True, poly_gate = np.log10(EPCAM_gate),
                  subset = 'Live and single_cells and singlets and (Cells == "13D7_iPSC")', xlim = (1,7))

# %% [code]
EPCAMOp = PolygonOp(
    name='EPCAM_plus',
    xchannel='APC-Cy7-A',
    ychannel='SSC-A',
    vertices=EPCAM_gate,
    xscale='log',
    yscale='log')

#SingletOp.estimate(ex_auto2)
ex_9 = EPCAMOp.apply(ex_8)
#tdTOp.default_view(subset='Live and single_cells and singlets').plot(ex_4)

# %% [code]
doublepos_gate = [(10**2.75, 10**2.75),
              (10**2.75, 10**5),
              (10**6, 10**5.4),
              (10**6, 10**2.75)]

flowPlotHist_gate(ex_8, channel_x = 'PE-Texas Red-A', channel_y = 'PE-Cy5-5-A', logscale = True, poly_gate = np.log10(doublepos_gate),
                  subset = 'Live and single_cells and singlets and (Cells == "1F") and Dox == 1000', xlim = (1,5), ylim = (0,5), cbar = True)
plt.xlabel("FOXL2-tdTomato", fontsize = 14)
plt.ylabel("CD82-PerCP-Cy5.5", fontsize = 14)
plt.title("1F +Dox (1000ng/mL)", fontsize = 16)
plt.savefig("2022-06-12_dox.png", dpi = 256, bbox_inches = "tight")

# %% [code]
doubleop = PolygonOp(
    name='double_pos',
    xchannel='PE-Texas Red-A',
    ychannel='PE-Cy5-5-A',
    vertices=doublepos_gate,
    xscale='log',
    yscale='log')

#SingletOp.estimate(ex_auto2)
ex_10 = doubleop.apply(ex_9)

# %% [code]
flow_df = ex_10.data

flow_df['tdTplus_EPCAMminus'] = flow_df['tdT_plus'] & ~flow_df['EPCAM_plus']
flow_df['CD82plus_EPCAMminus'] = flow_df['CD82_plus'] & ~flow_df['EPCAM_plus']
flow_df['tdTplus_CD82plus_EPCAMminus'] = flow_df['double_pos'] & ~flow_df['EPCAM_plus']

#flow_df.to_csv("2021-05-23_raw.csv")
samples = flow_df['Path'].unique()
singlets = flow_df[(flow_df['singlets'] == True) & (flow_df['single_cells'] == True)  & (flow_df['Live'] == True)]


print(samples)

# %% [code]
mean_stats = singlets.groupby("Path").mean(numeric_only = False)
mean_stats.to_csv("2021-07-31b_day5_means_singletfiltered.csv")

# %% [code]
# for sample in samples:
#     print('Cells: ' + sample)
#     singlets_count = singlets[singlets['Cells'] == sample]
#     PGCLC_count = singlets_count[singlets_count['GFP_plus'] == True]
#     print('Total live singlets: '+str(len(singlets_count)))
#     print('Total GFP+: ' + str(len(PGCLC_count)))
#     print('Percentage: {0}\n'.format( len(PGCLC_count)/(len(singlets_count)) * 100))

# %% [code]
mean_stats2 = singlets.groupby(["Cells","Dox"]).mean(numeric_only = False)
mean_stats2.dropna(inplace = True)
mean_stats2.to_csv("2021-07-31b_day5_stats_singletfiltered.csv")

# %% [code]
mean_stats3 = singlets.groupby(["Cells"]).mean(numeric_only = False)
mean_stats3.dropna(inplace = True)
mean_stats3.to_csv("2021-07-31b_day5_pooled.csv")

# %% [code]
#tdT plot
#
#flowPlotHist_gate(ex_8, channel_x = 'PE-Texas Red-A', channel_y = 'SSC-A', logscale = True, poly_gate = np.log10(tdT_gate),
#                  subset = 'Live and single_cells and singlets and (Cells == "13D7") and (Condition == 5)', xlim = (2,6))

# %% [code]
mean_stats_all = flow_df.groupby(["Cells","Dox"]).mean(numeric_only = False)
mean_stats_all.dropna(inplace = True)
mean_stats_all.to_csv("2021-07-31b_day5_stats_all.csv")

# %% [code]

flowPlotHist_gate(ex_10, channel_x = 'FSC-A', channel_y = 'SSC-A', logscale = True, poly_gate = np.log10(doublepos_gate),
                  subset = 'Live and single_cells and singlets and (Cells == "13D7" and double_pos)', xlim = (3.5, 5.5), ylim = (2.5, 5.5), cbar = True)


# %% [code]
plot_df = pd.read_csv("tdTPlus_CD82Plus.csv")
plot_df

# %% [code]
import seaborn as sns

# %% [code]
sns.lineplot(data = plot_df, x = "Dox", y = "double_pos", hue = "Clone")
plt.xlabel("Doxycycline (ng/mL)", fontsize = 14, color = "black")
plt.ylabel("FOXL2+ CD82+", fontsize = 14, color = "black")
plt.ylim([0,1])
plt.xlim([0,2000])
plt.xticks([0, 100, 200, 500, 1000, 2000], fontsize = 11)
plt.yticks(fontsize = 11)
plt.legend(fontsize = 12)
plt.title("Dox dose dependence for granulosa induction", fontsize = 16, color = "black")
plt.savefig("dose-dependence.png", dpi = 256, bbox_inches="tight")

# %% [code]
```
