## Supplementary figures and images for "Directed Differentiation of Human iPSCs to Functional Ovarian Granulosa-Like Cells via Transcription Factor Overexpression"

### 2021-10-07_ELISA_results.png

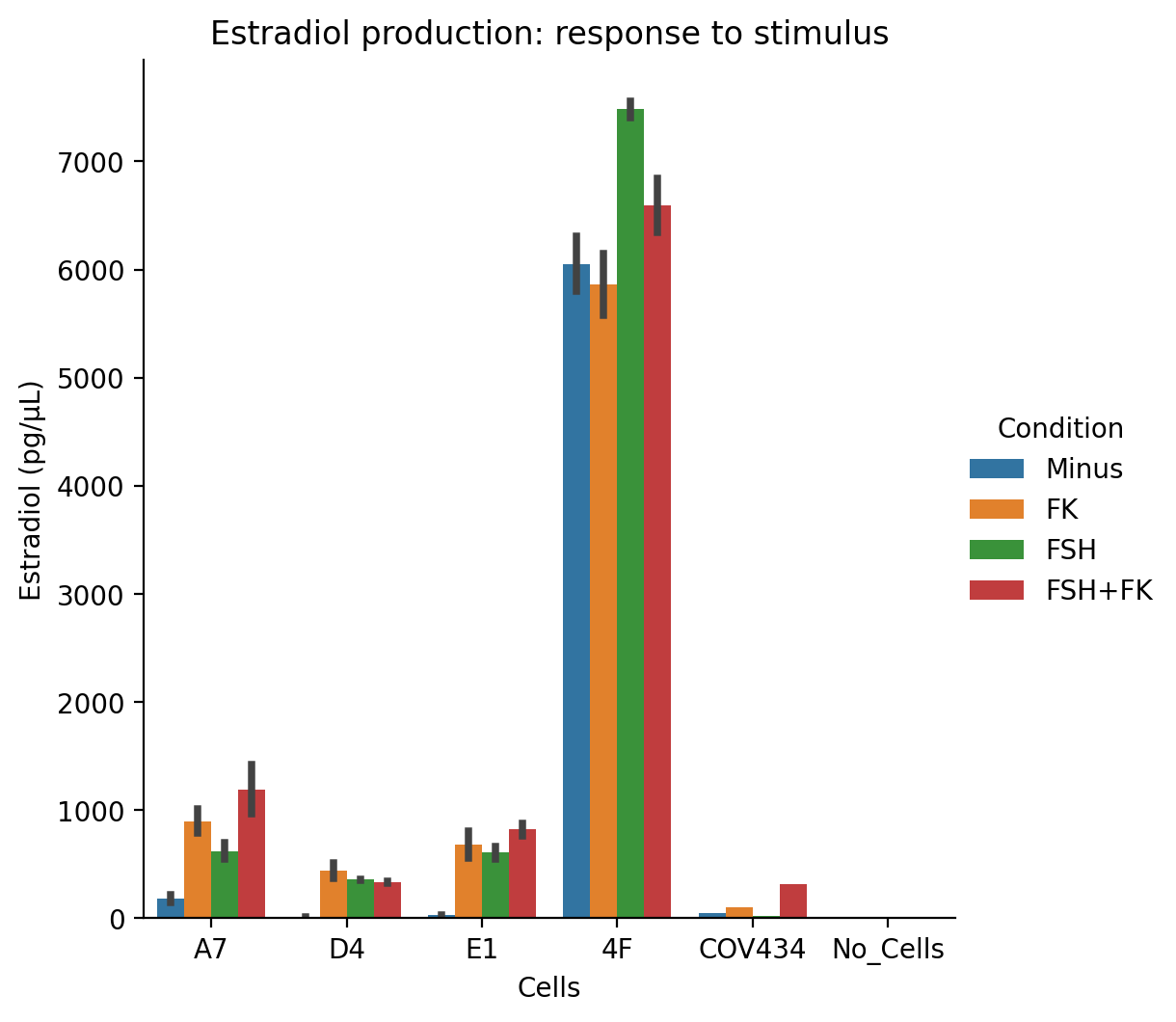

### 2021-10-07_ELISA_results_no_4F.png

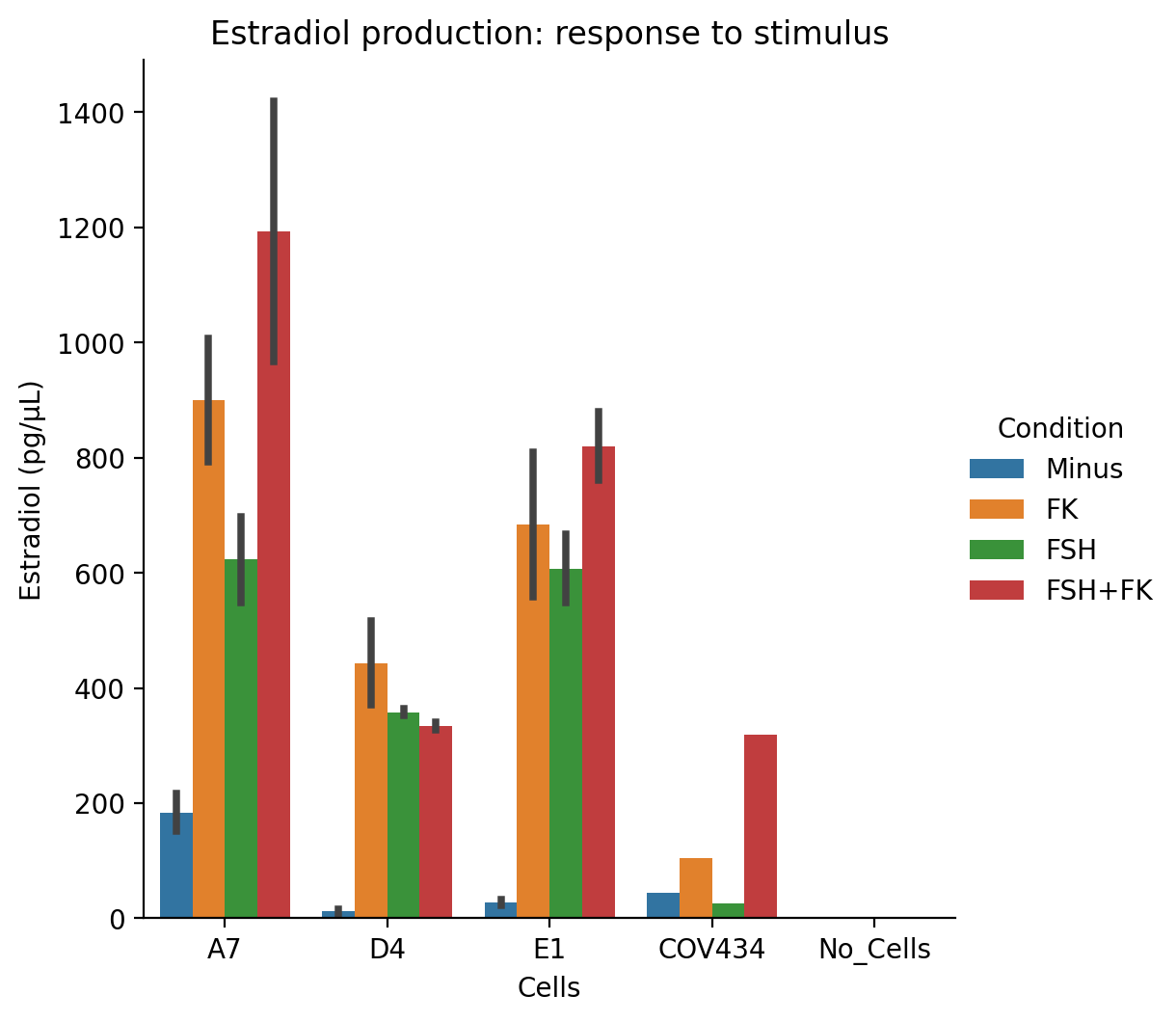

### 2021-10-07_ELISA_results_no_4F_replot.png

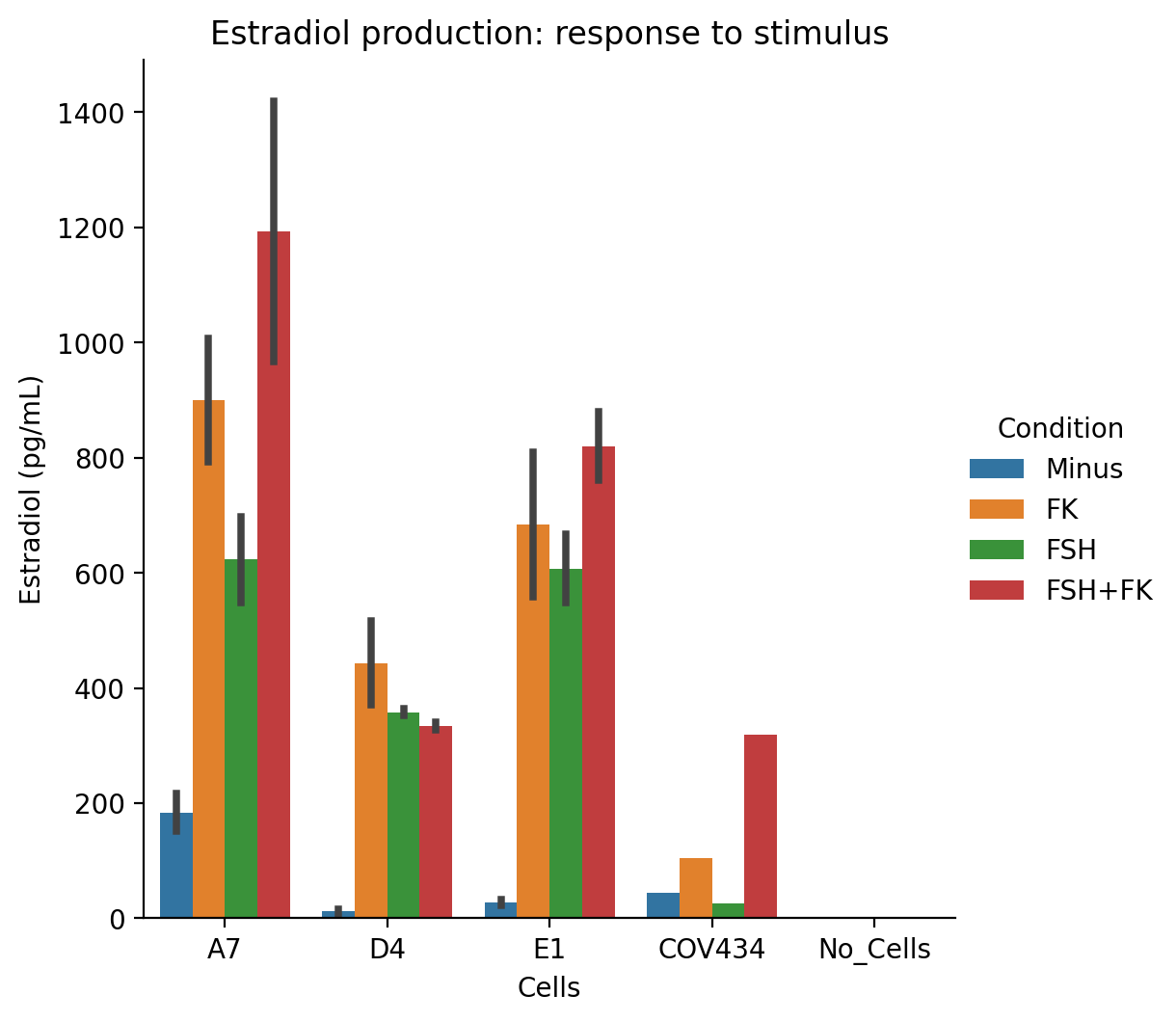

### 2021-10-07_ELISA_results_no_4F_rereplot.png

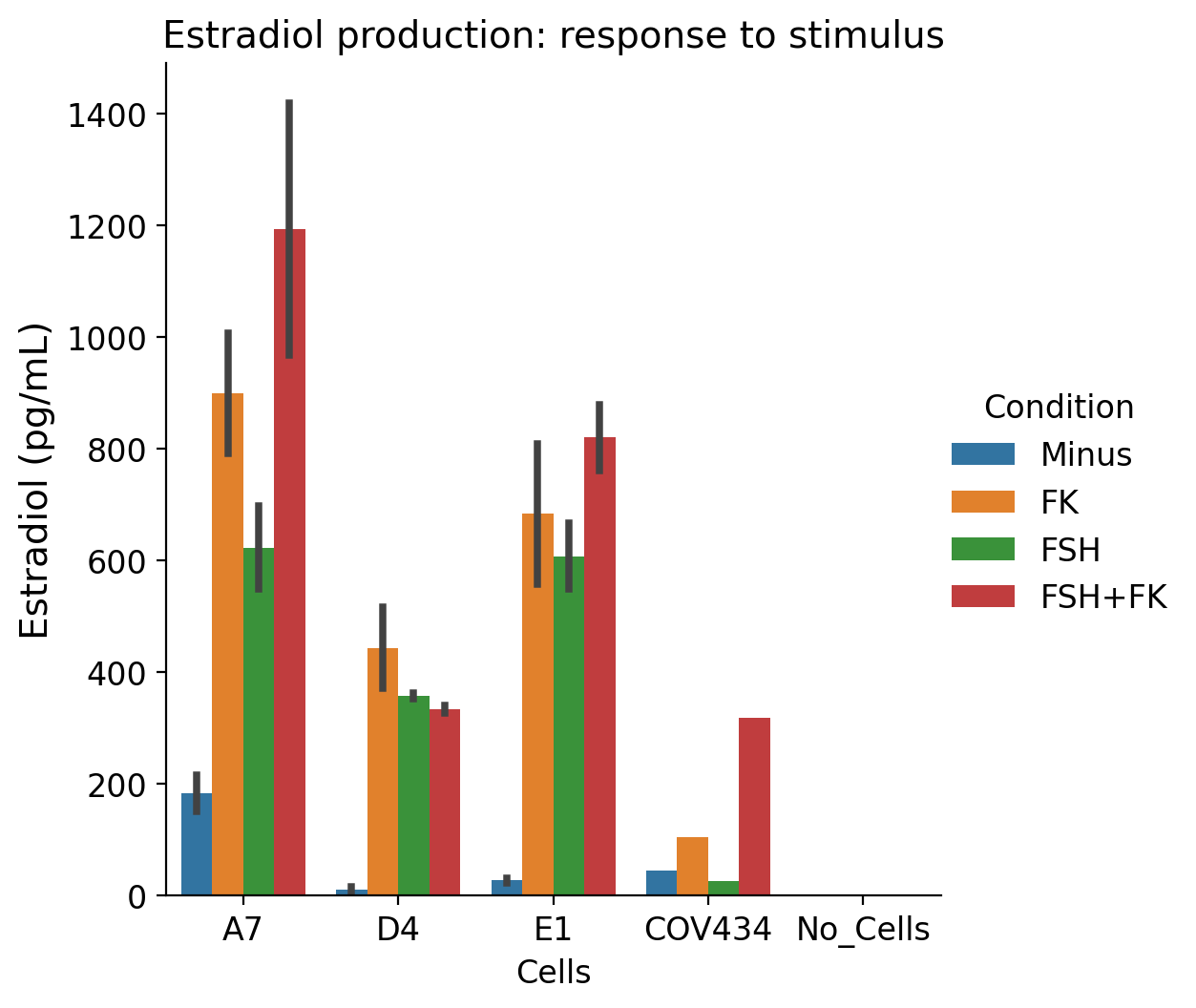

### 2021-10-07_ELISA_results_no_4F_rereplot_inkscape.png

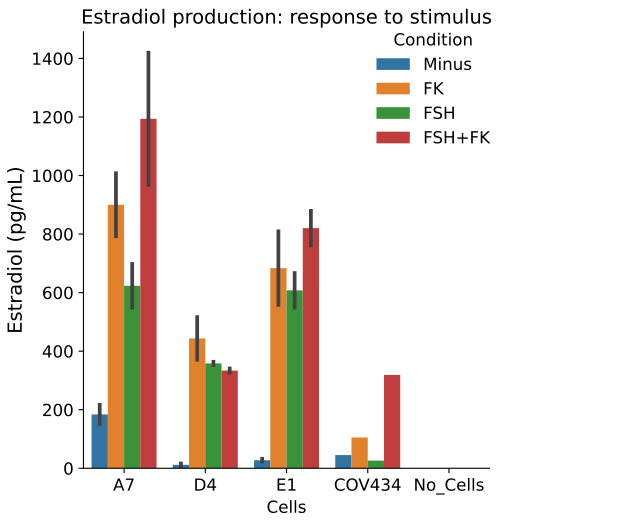

### 2021-10-07_ELISA_results_replot2.png

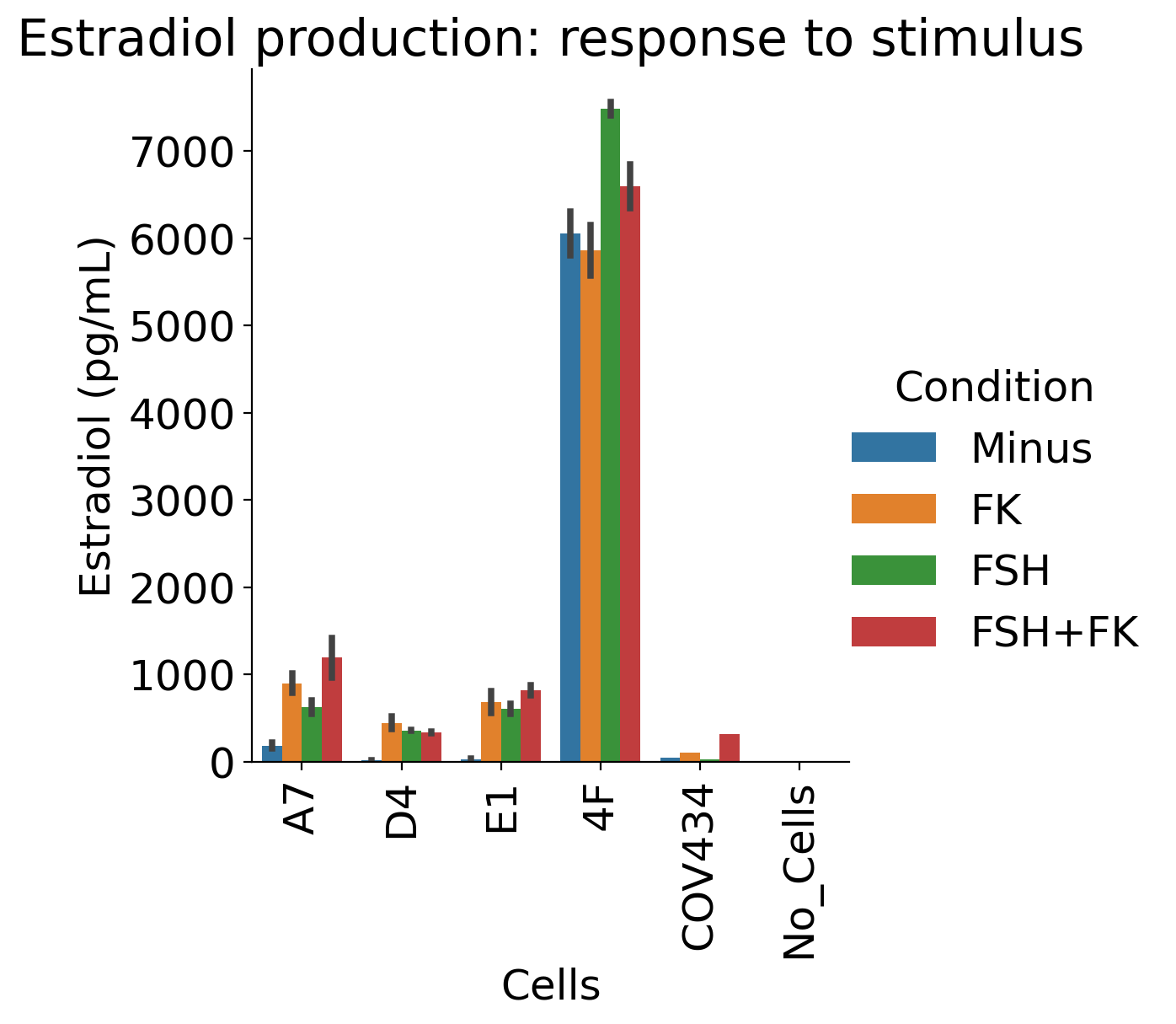

### 2021-10-07_ELISA_results_replot.png

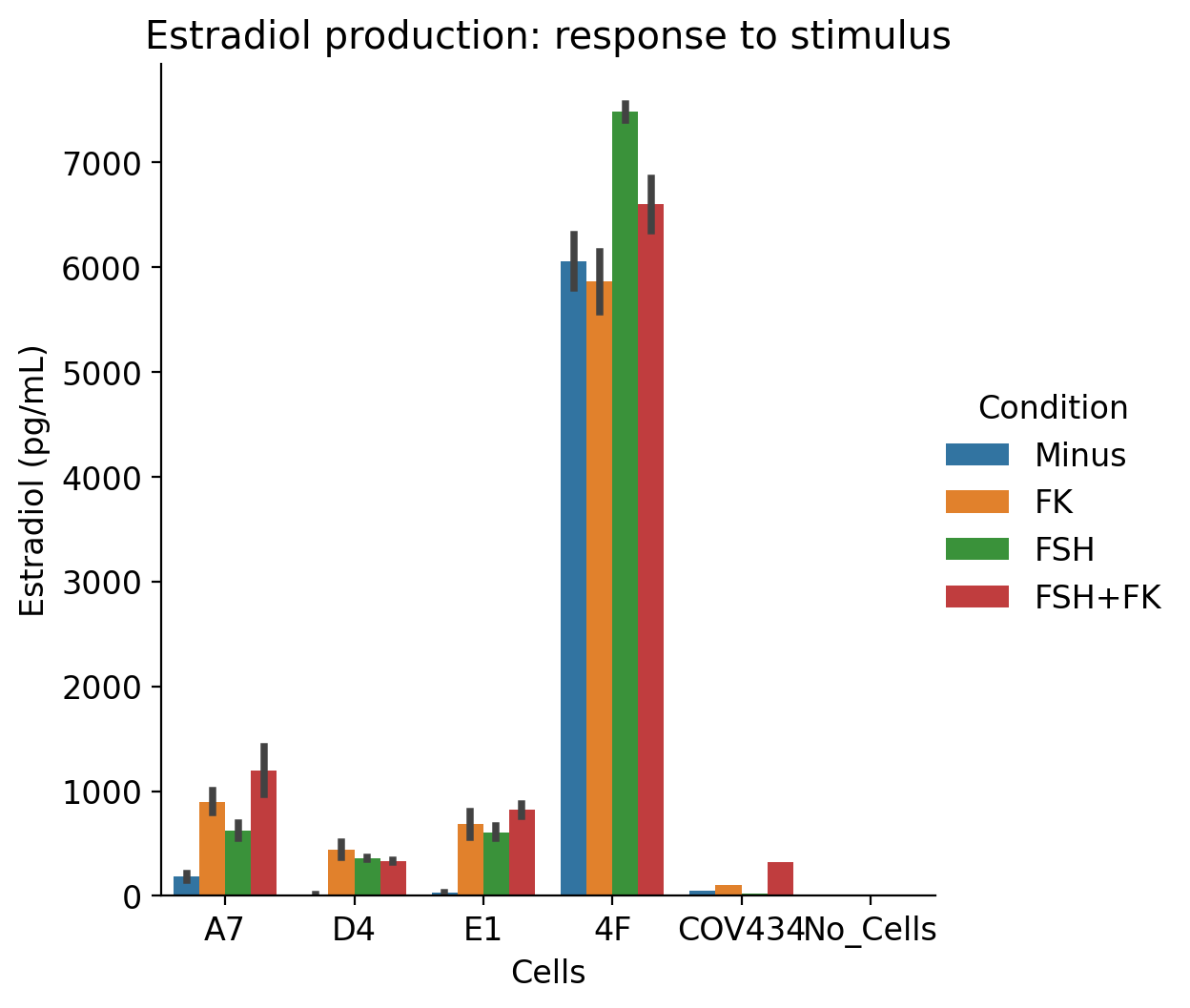

### 2021-10-07_standards_logistic_4param.png

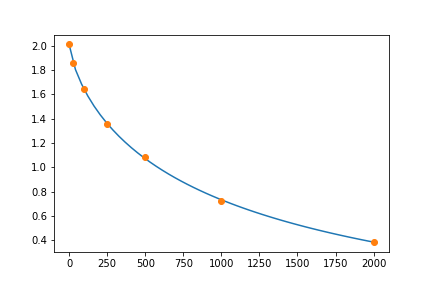

### _NEW_PLOT.png

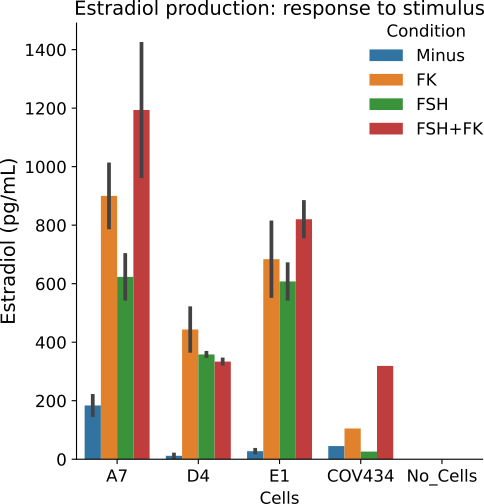

### ovaroid_time_plot.png

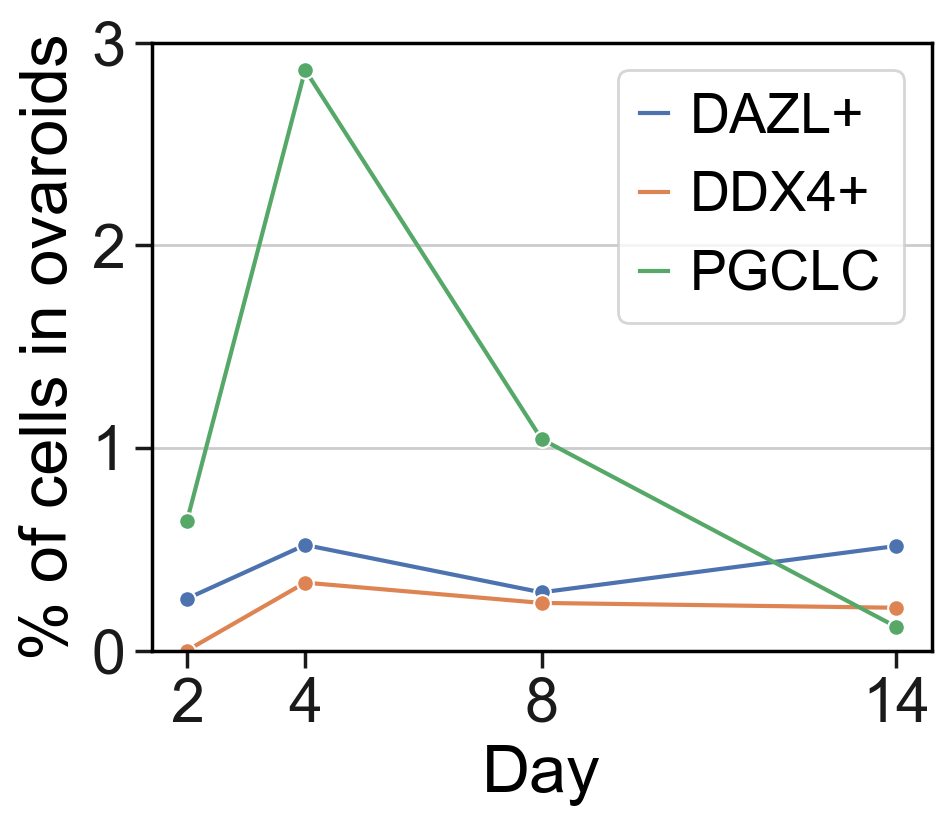

### pca_variance_ratio.png

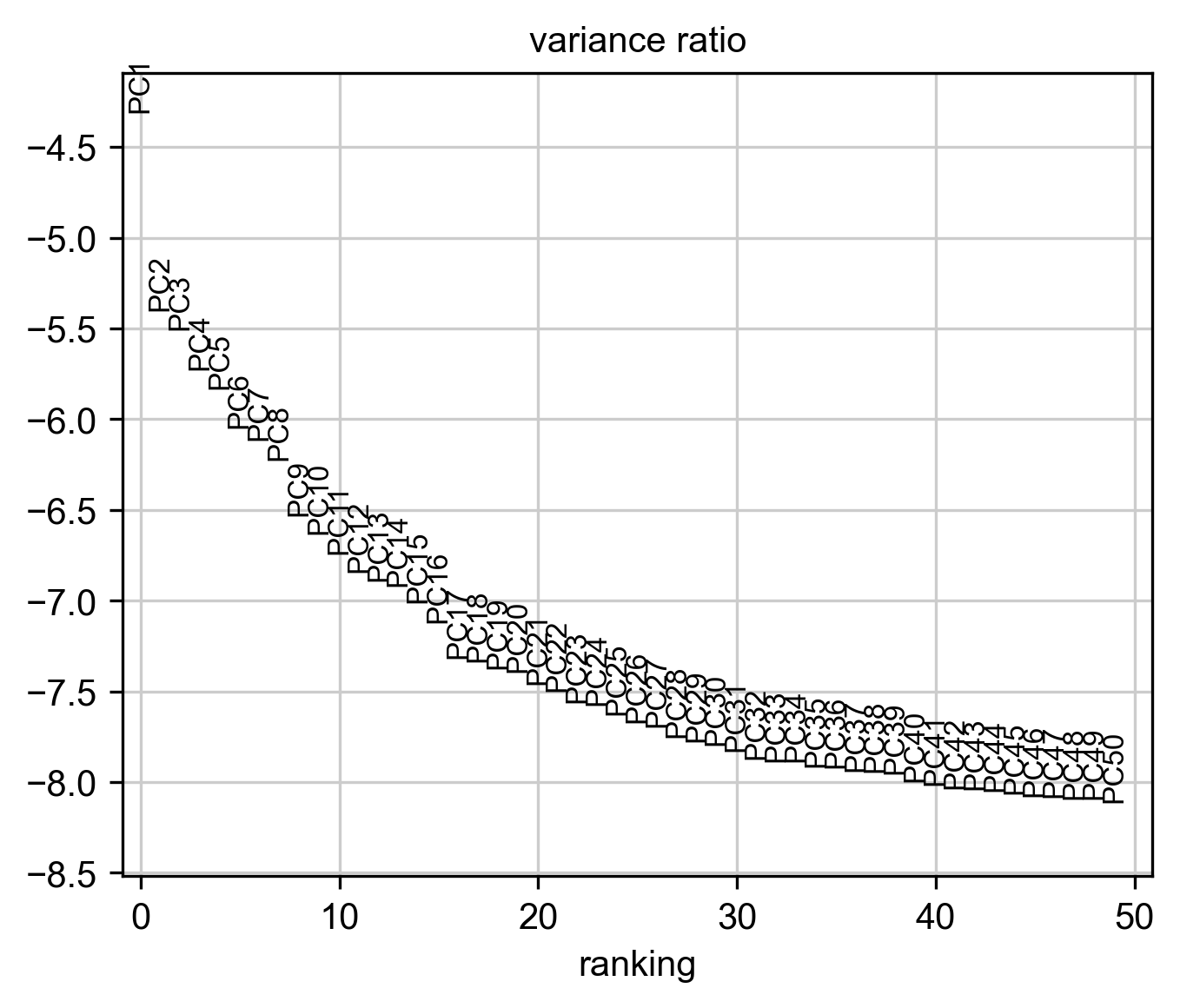

### scatter_gene_vs_transcript_counts.png

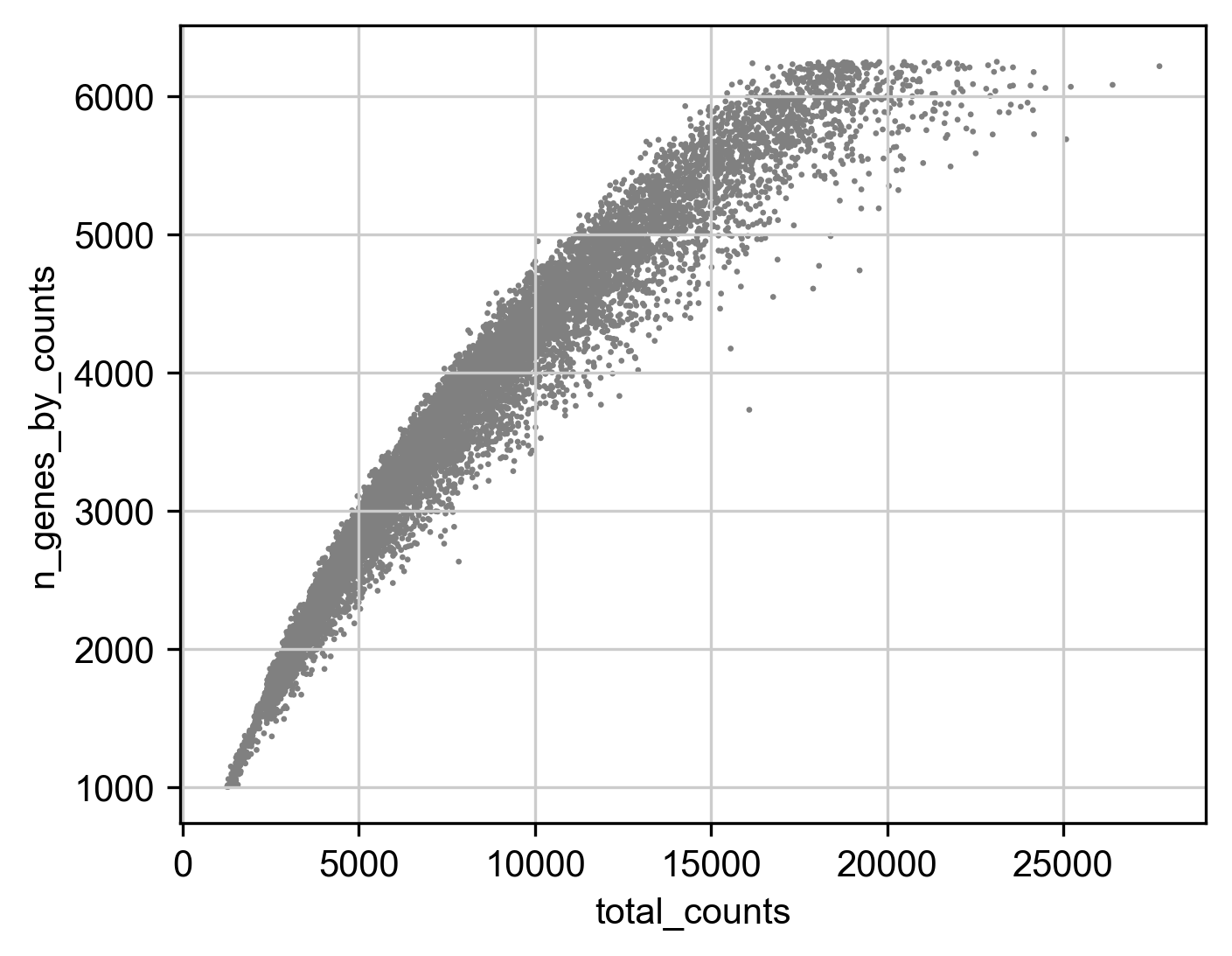

### umap_Day_regress.png

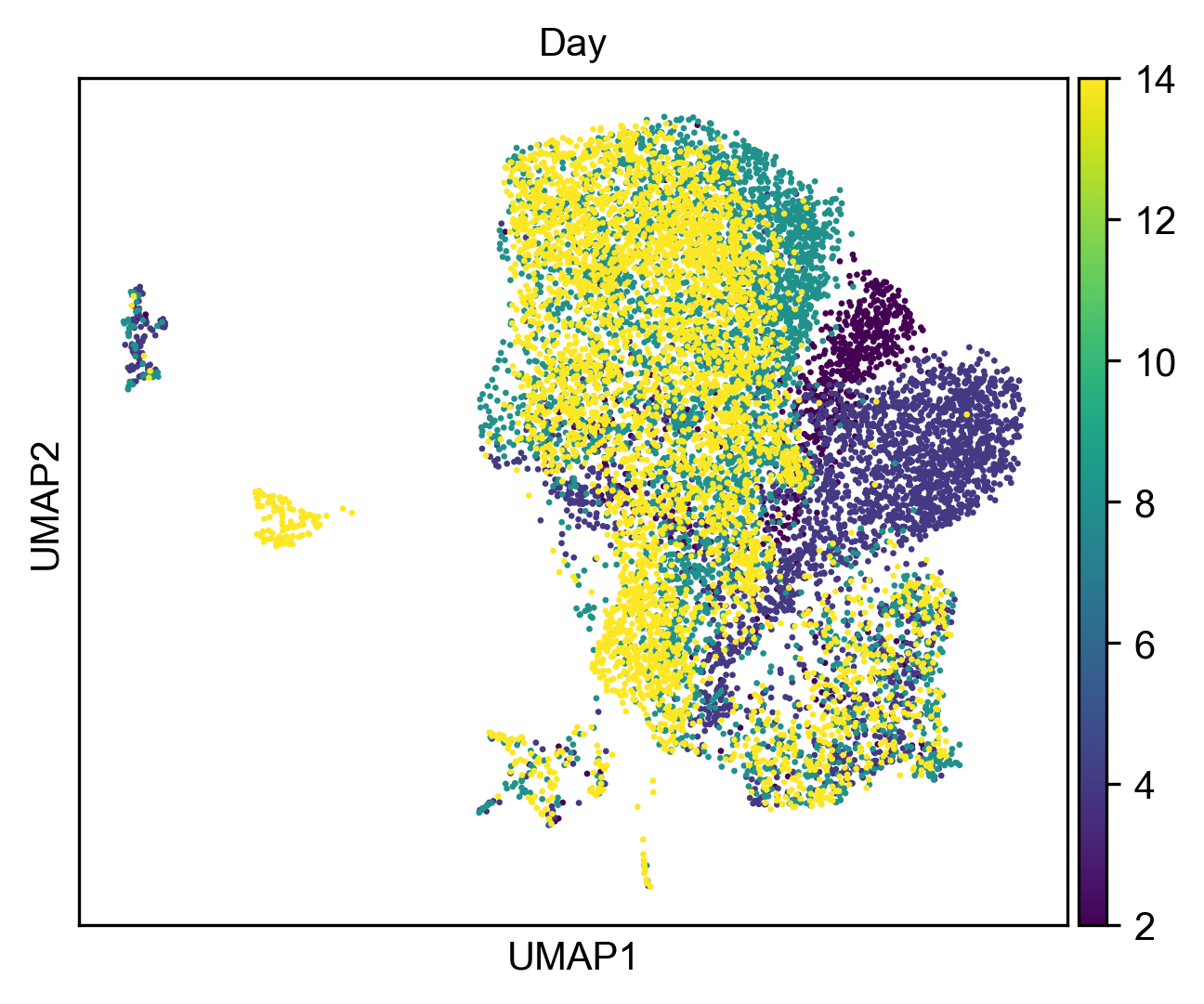

### umap_regress_FOXL2.png

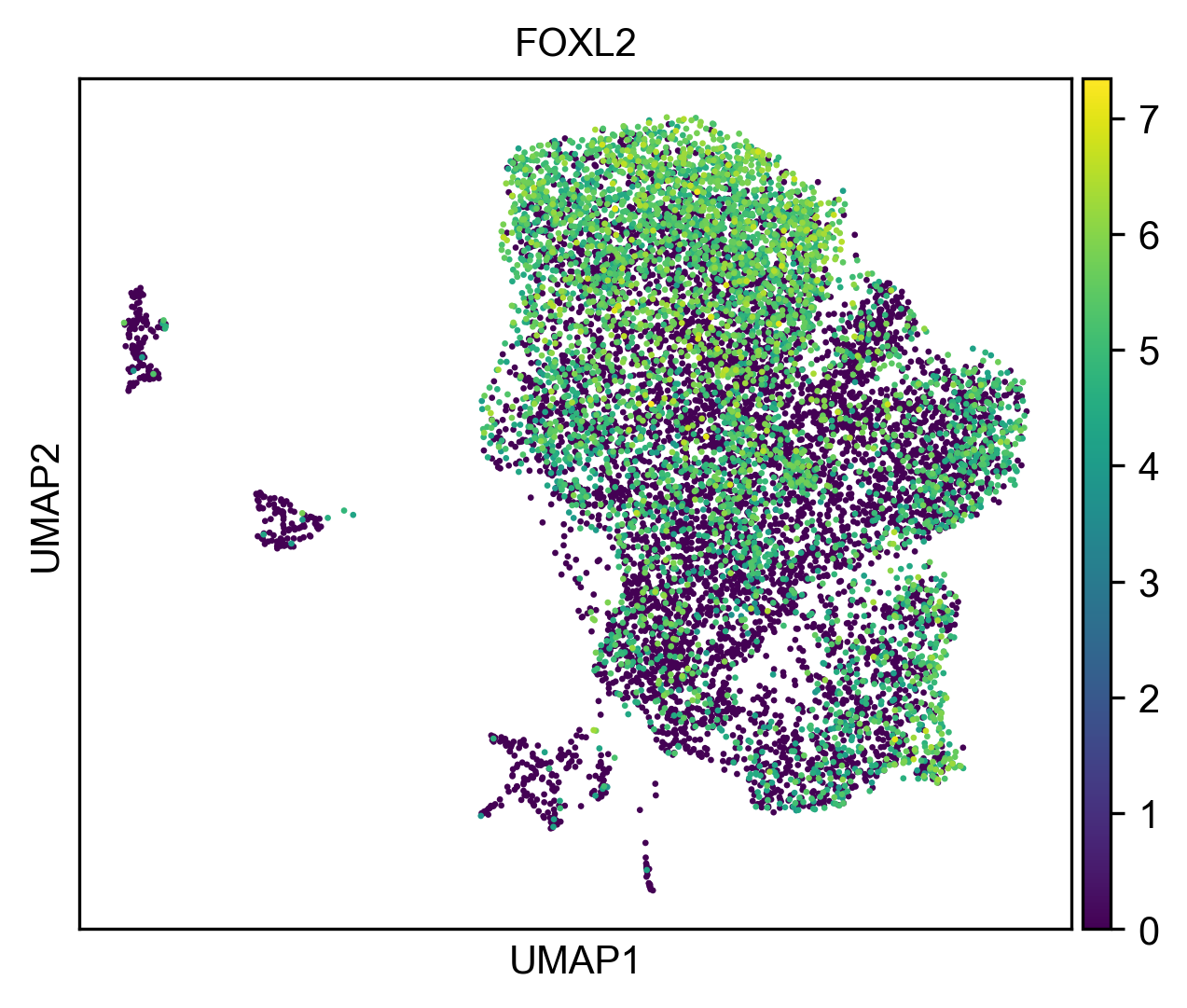

### umap_regress_KIT.png

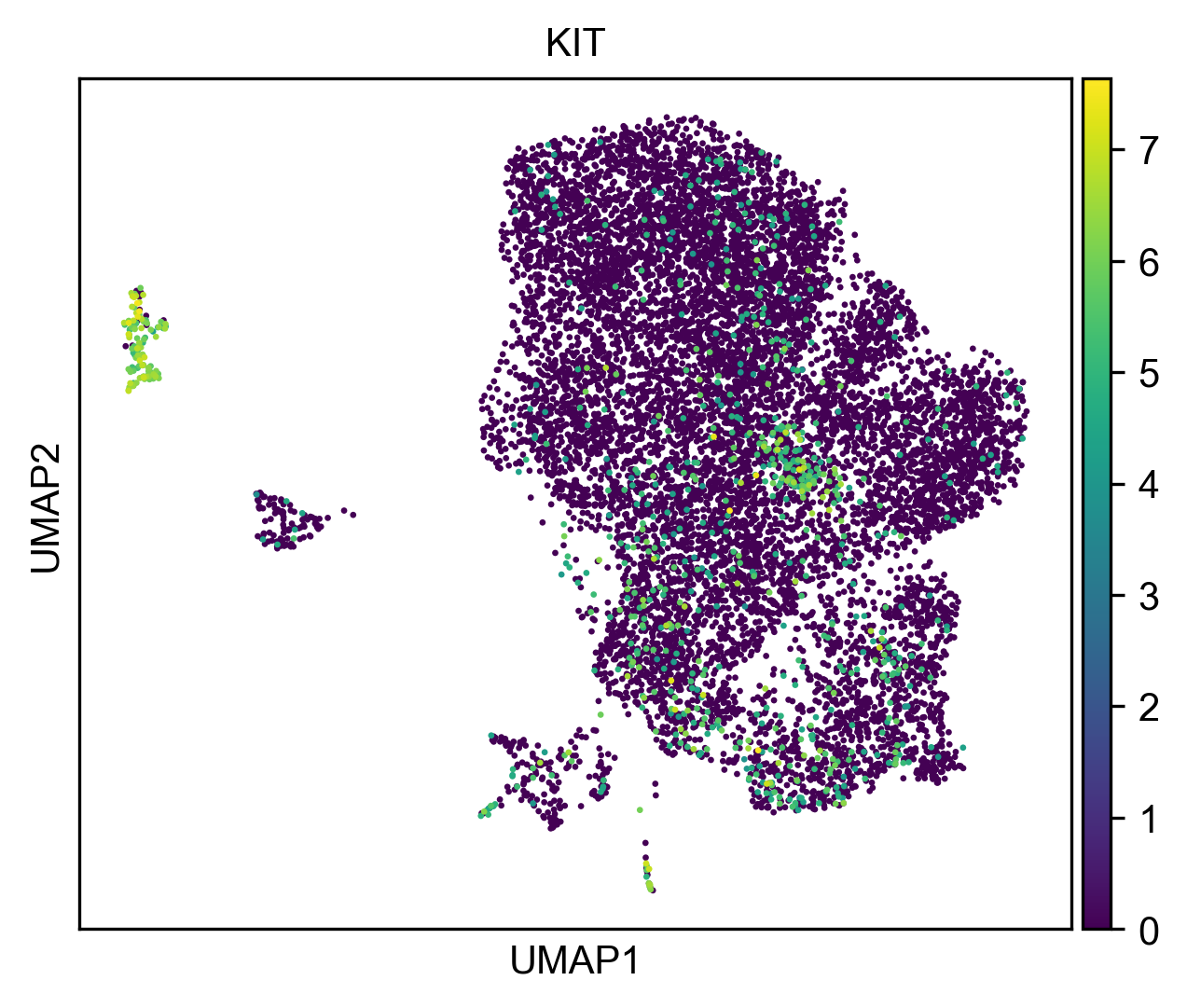

### umap_regress_NANOG.png

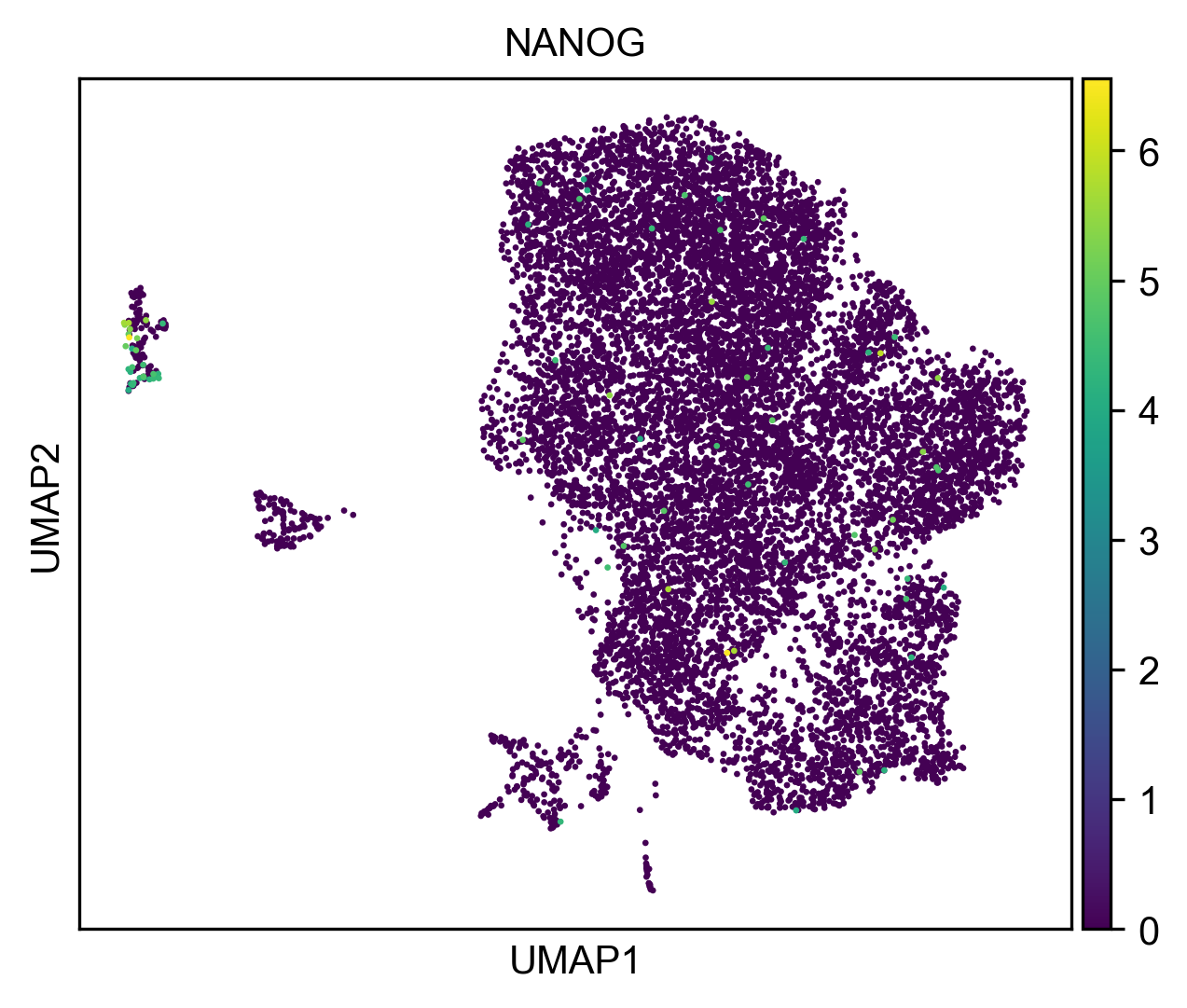

### umap_regress_PRDM1.png

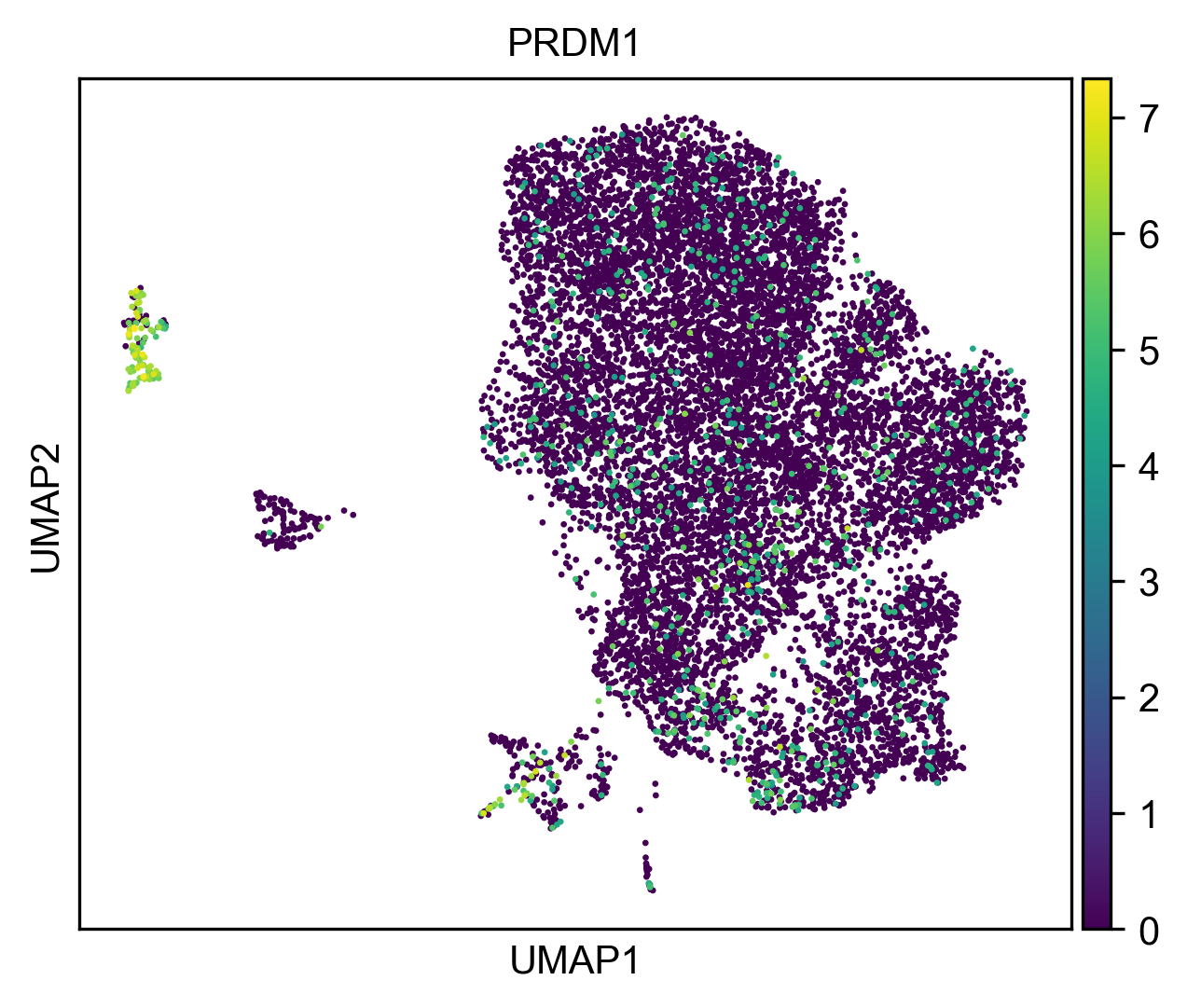

### umap_regress_WNT4.png

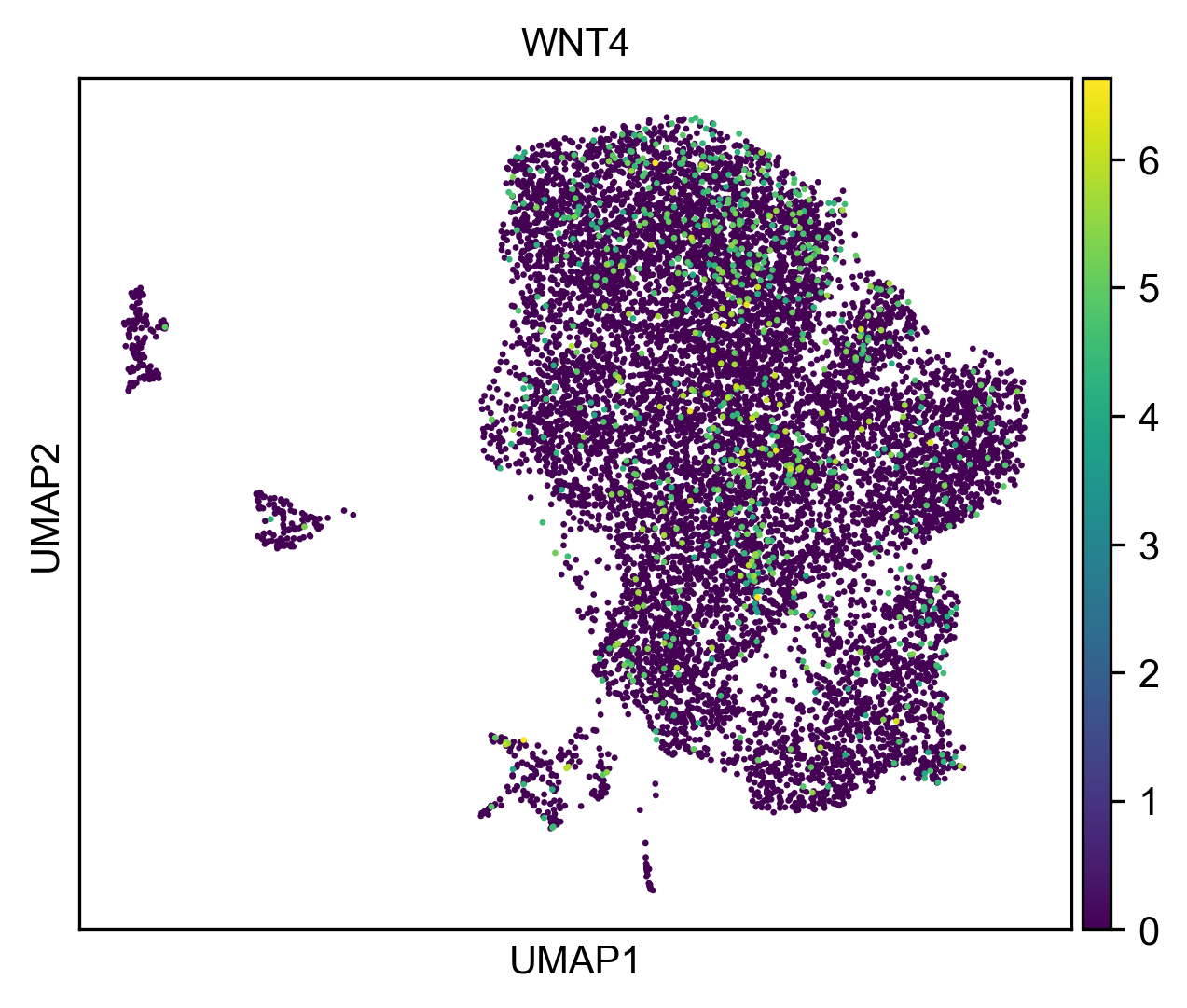

### umap_sample_regress.png

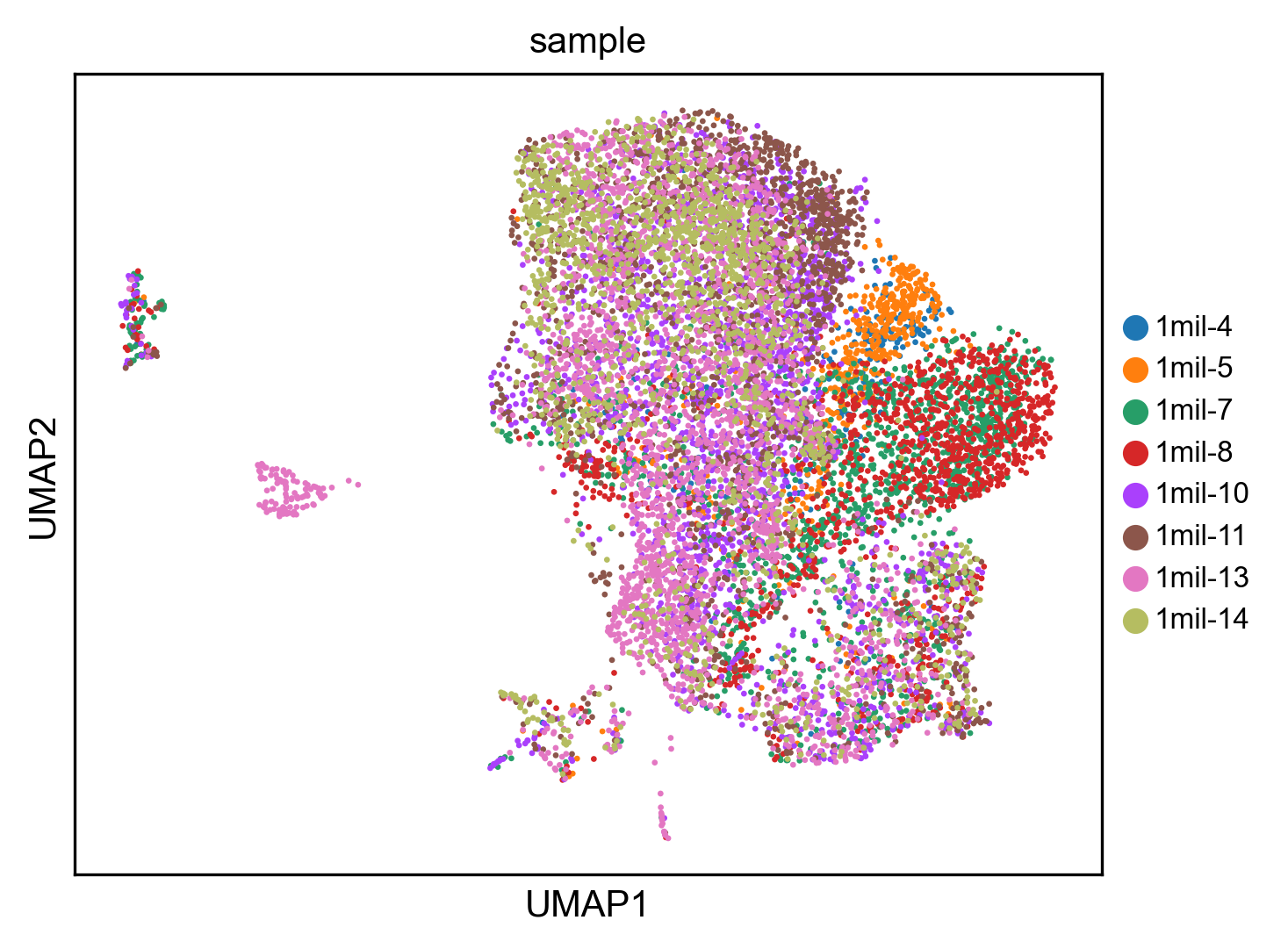

### violin_mito_pct.png

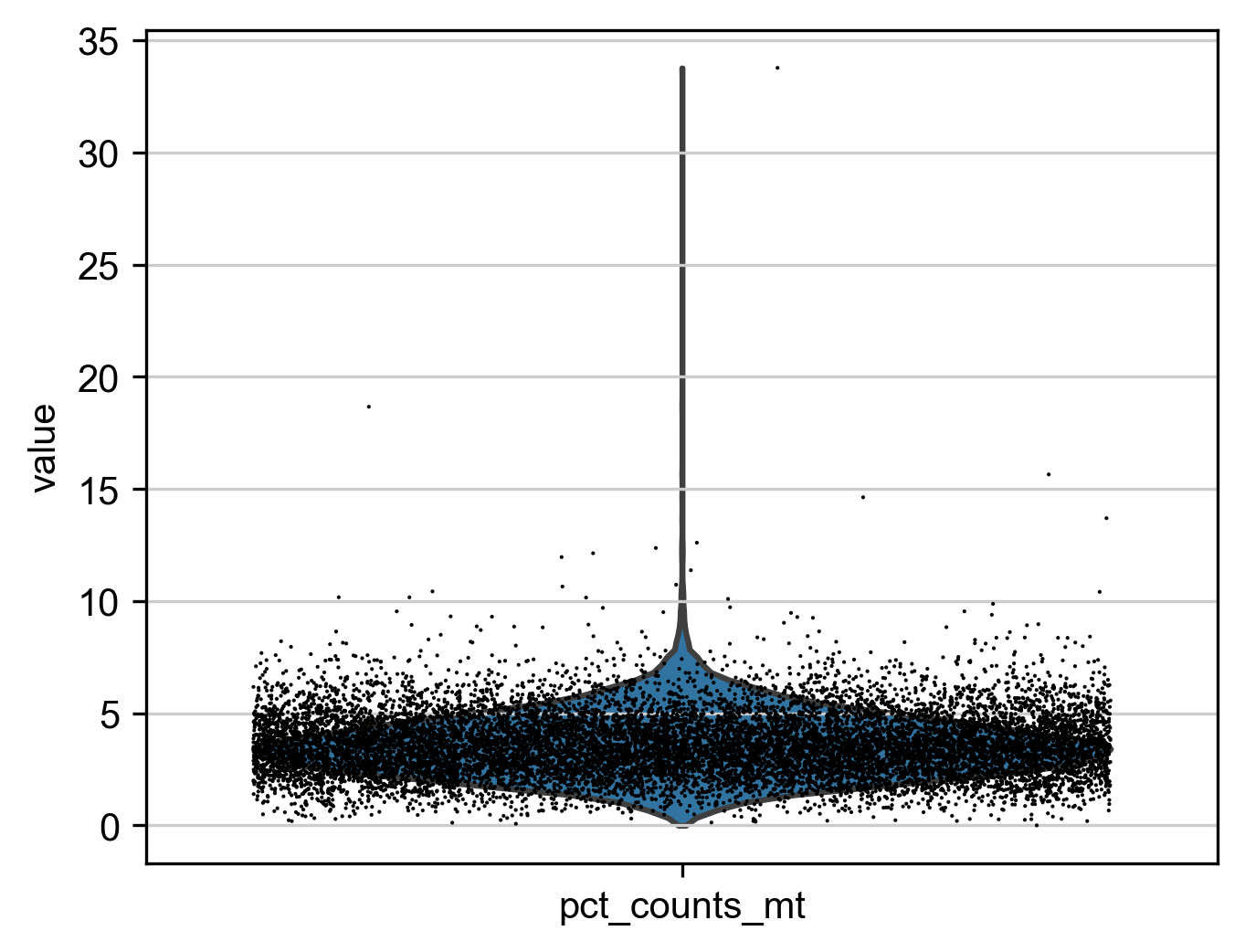

### violin_n_genes.png

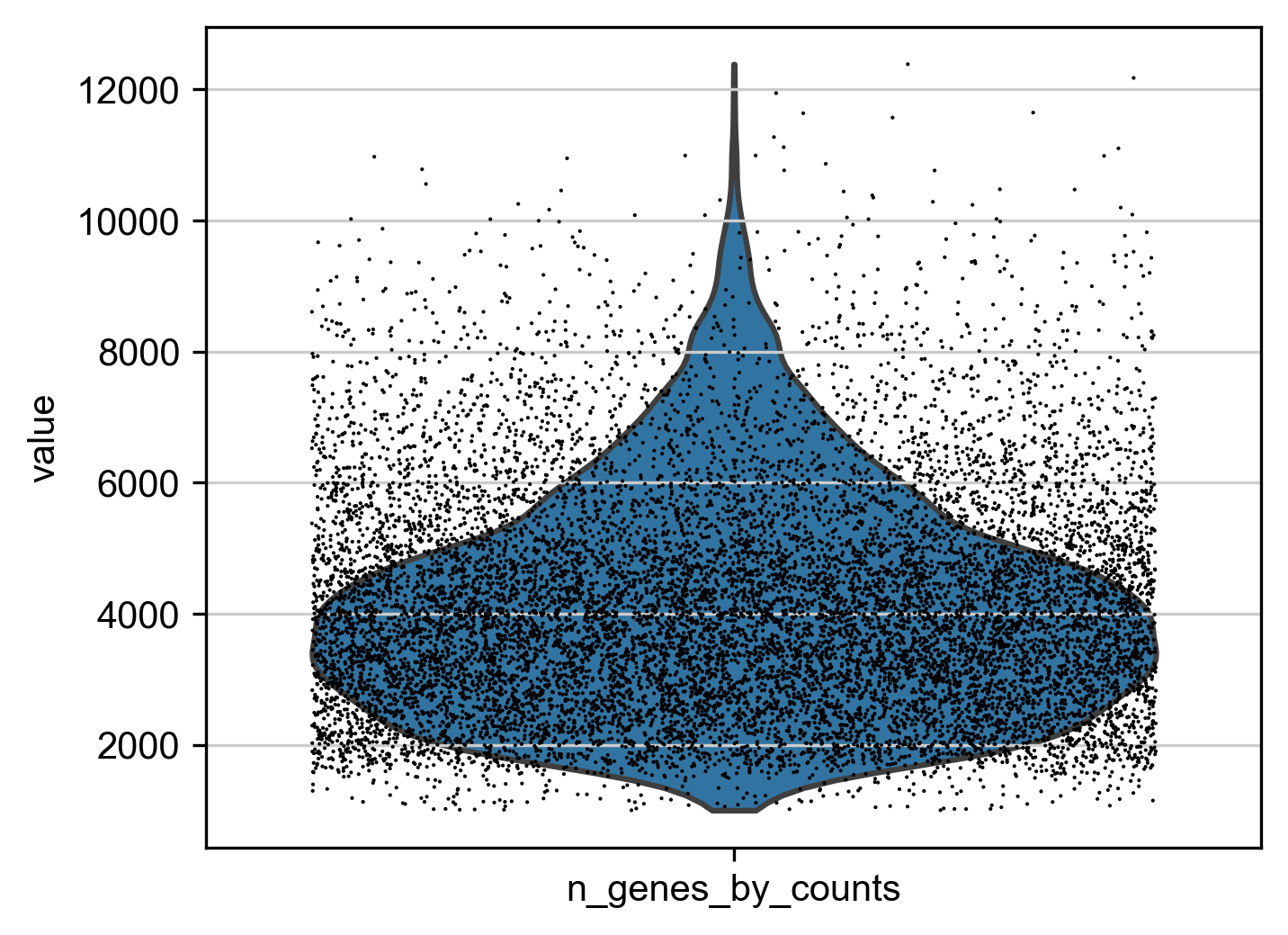

### violin_total_counts.png

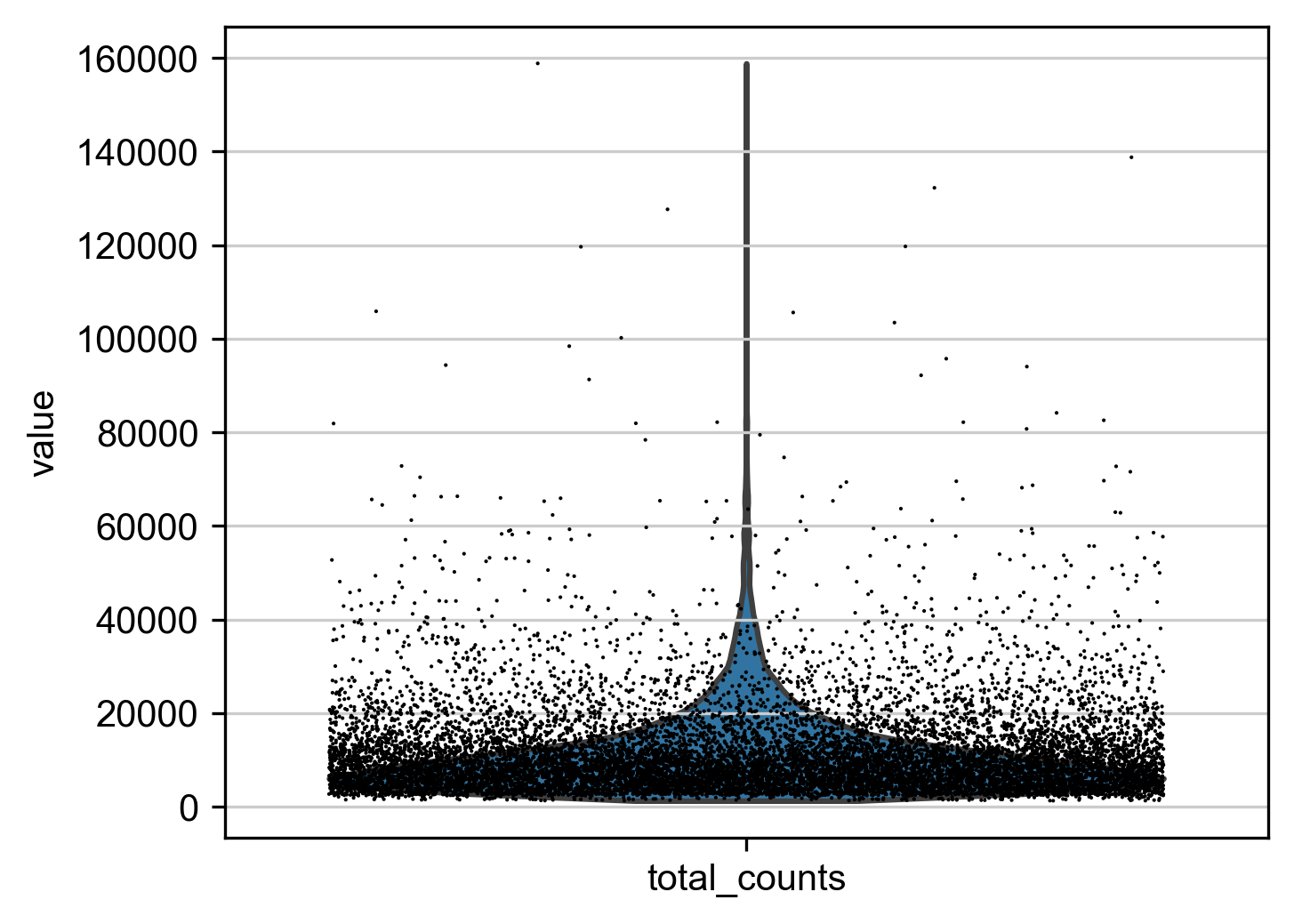

### volcano_FOXL2.png

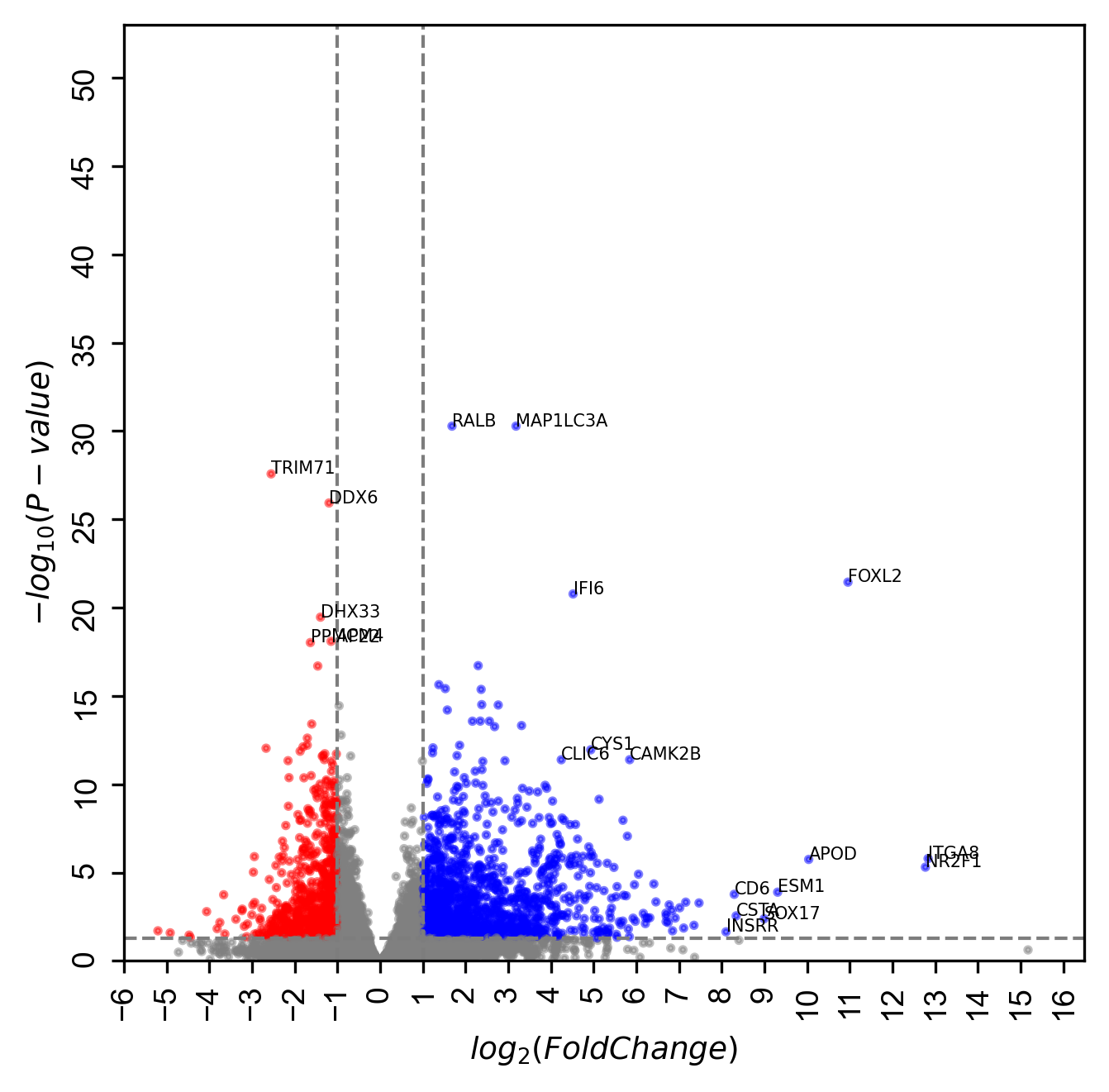

### volcano_GATA4.png

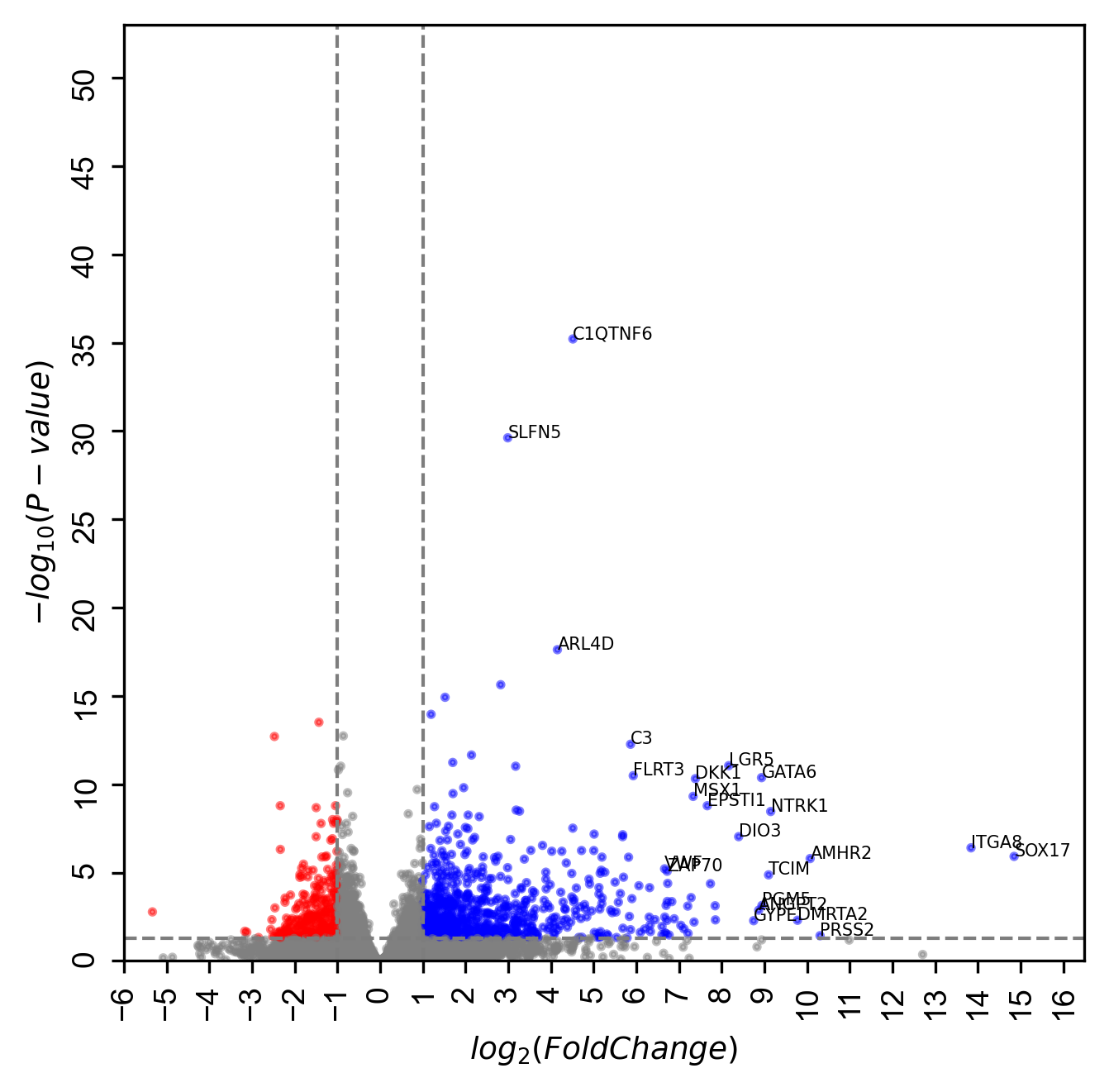

### volcano_NR5A1.png

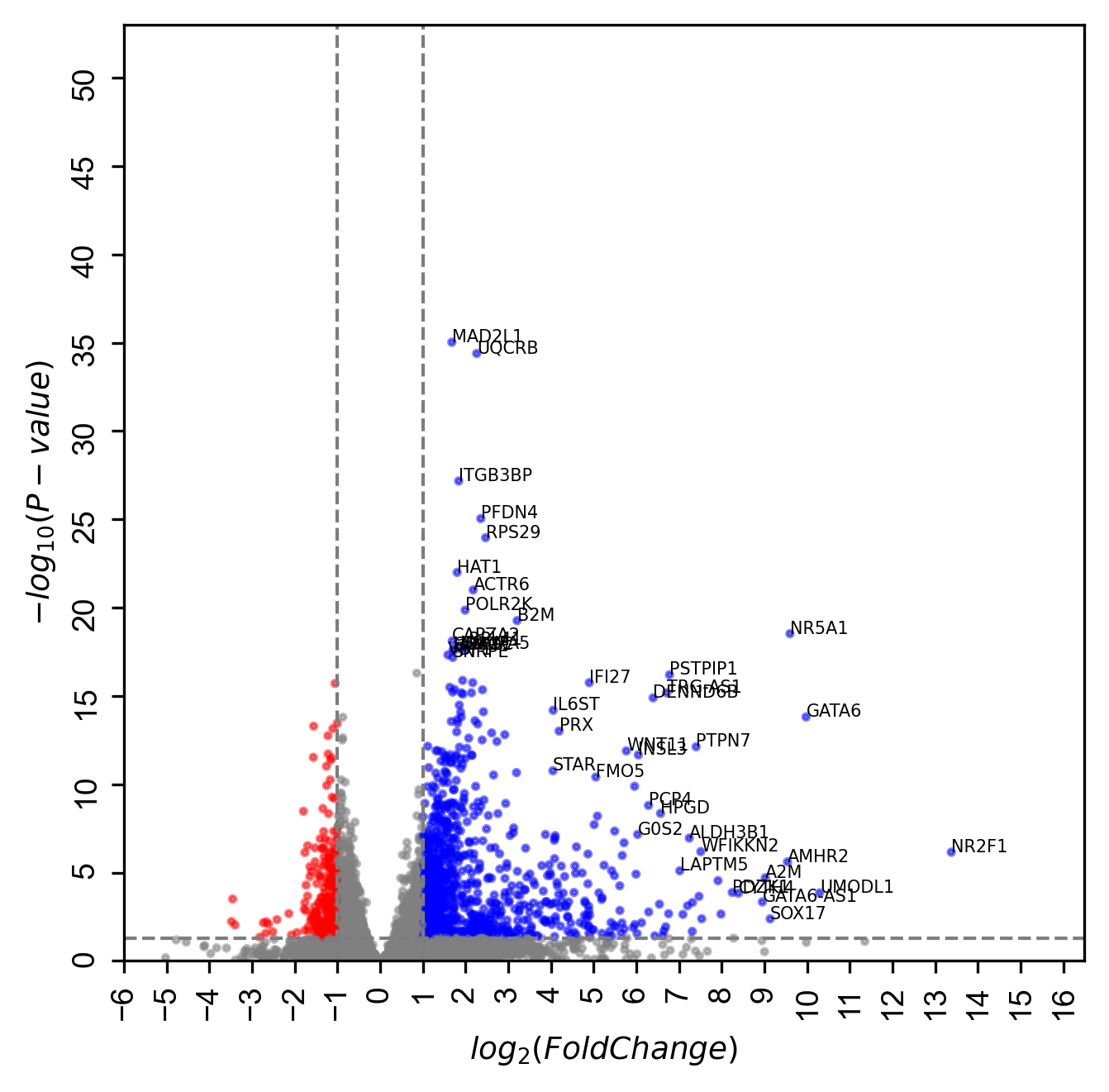

### volcano_RUNX1.png

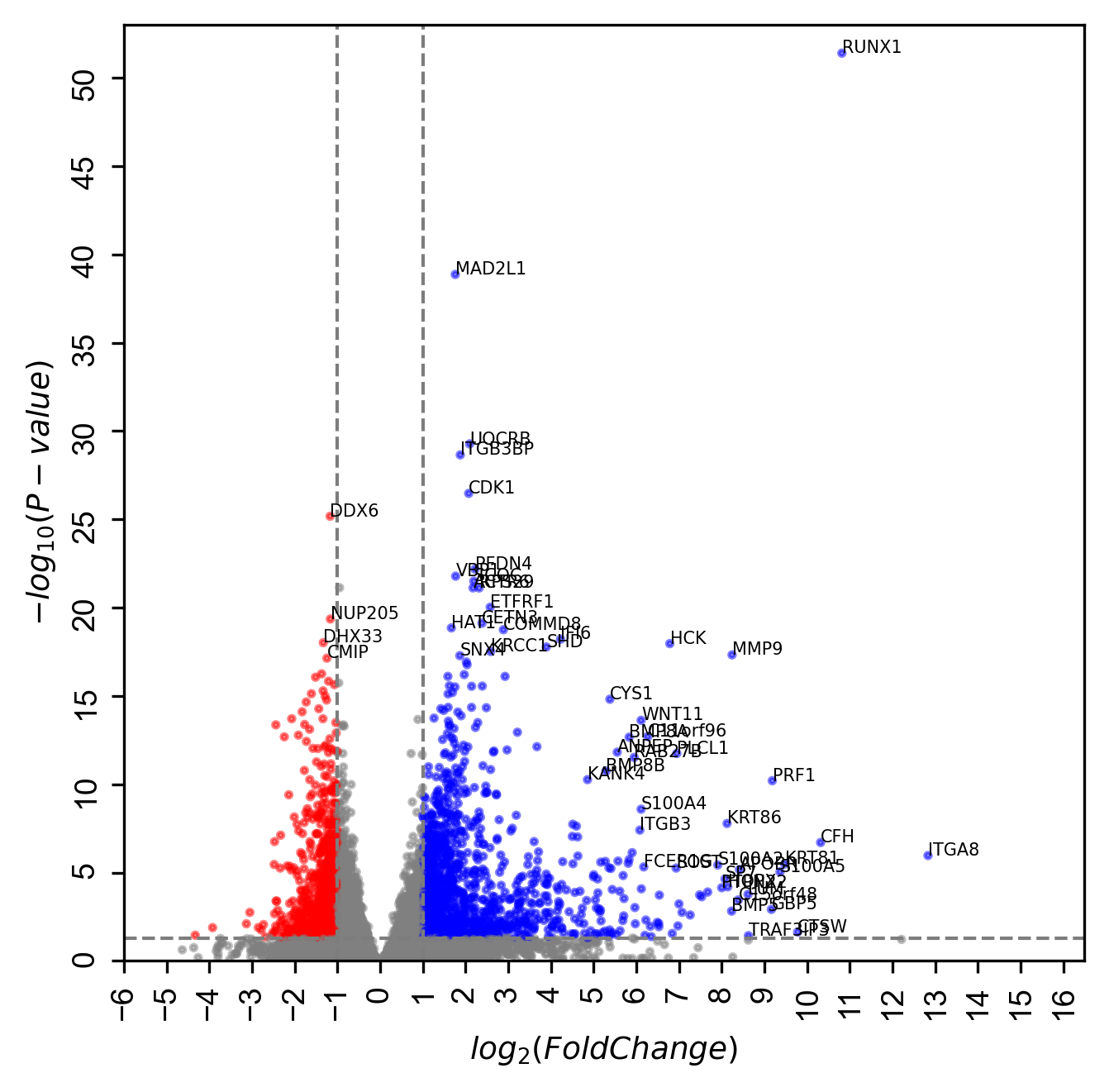

### volcano_TCF21.png

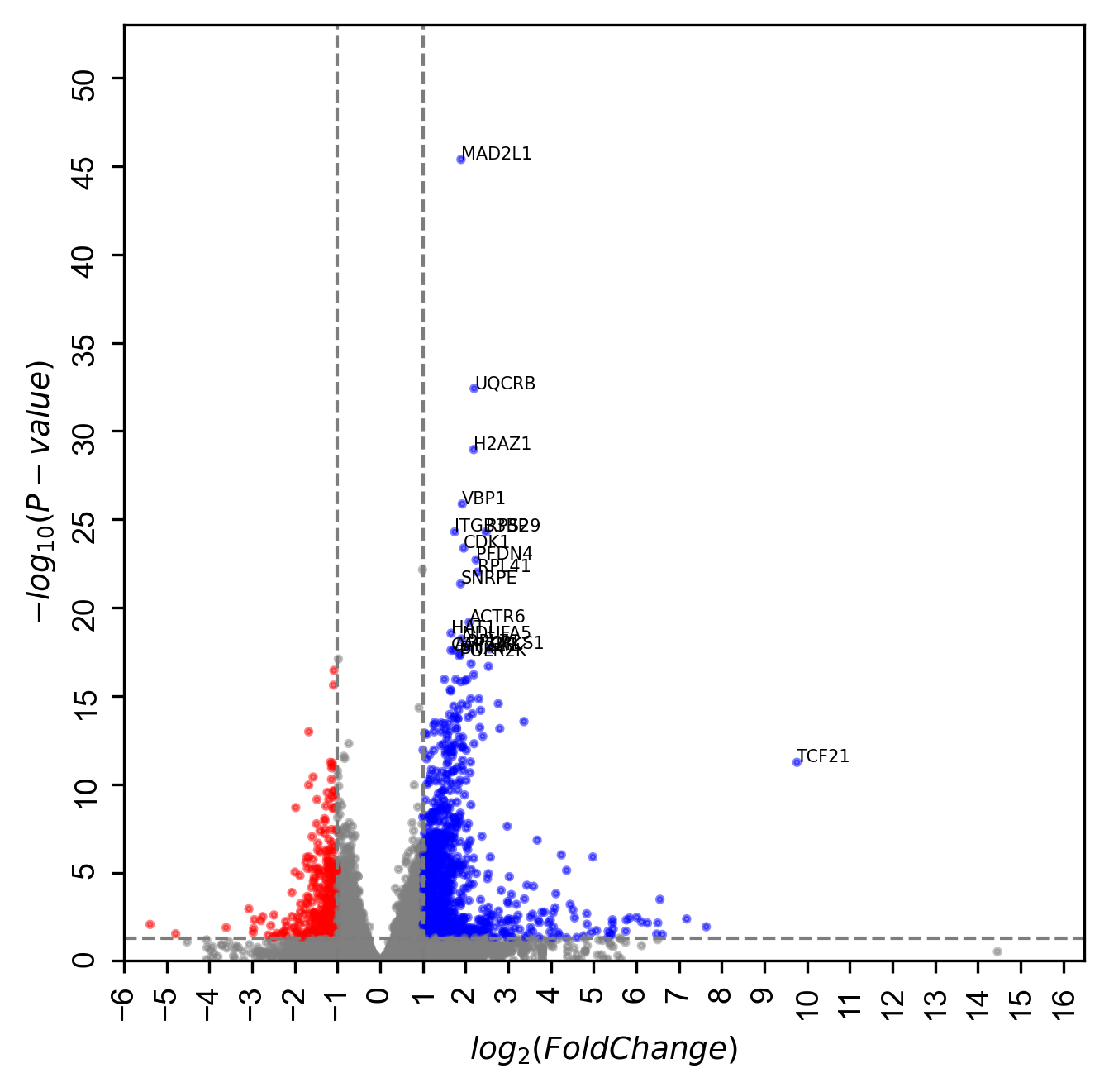

### volcano_WT1.png

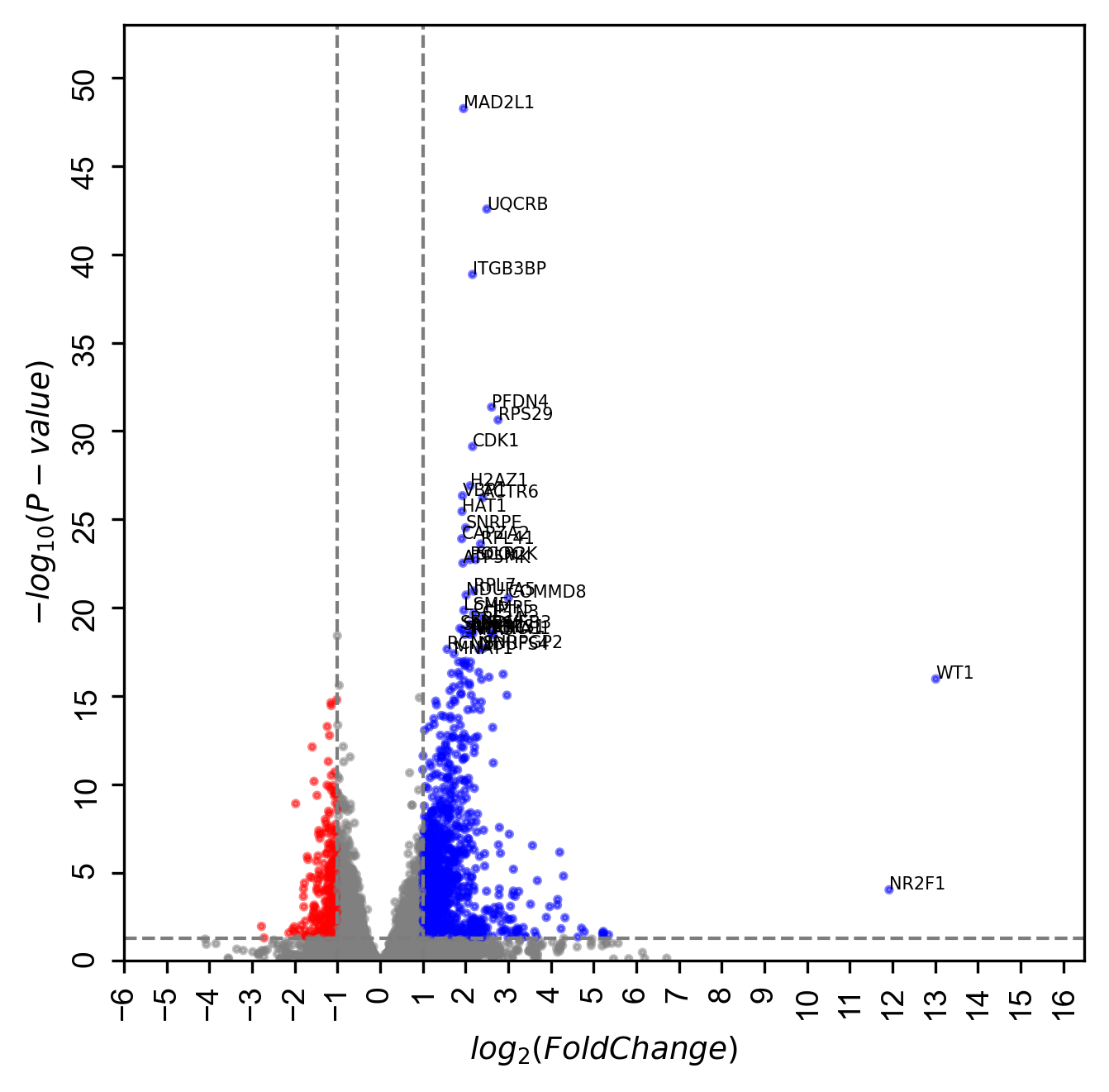
